# *Mycobacterium tuberculosis* diversity and macrophage heterogeneity dictate phagosomal acidification

**DOI:** 10.1101/2025.11.19.689035

**Authors:** Matteo Chiacchiaretta, Rita Sorrentino, Nadia Bresciani, Alessandra Aiello, Valentina Vanini, Samuel Zambrano, Davide Mazza, Federica Cugnata, Daniela Maria Cirillo, Alessandra Agresti, Delia Goletti, Paolo Miotto

## Abstract

Therapies counteracting pathogen-induced immune modulation of the host responses represent a promising improvement for tuberculosis treatments. Despite the documented role of host immune heterogeneity and bacterial genotypic background in determining infection outcomes, their interplay remains largely uncharacterized. We investigate the *Mycobacterium tuberculosis* (MTB)-human macrophage interaction considering both macrophage phenotypic variability and MTB genetic diversity.

Using single-cell techniques, we show how diverse MTB lineages fine-tune phagosome acidification differently based on macrophage phenotype (M1, M2), revealing very heterogeneous host-pathogen interactions.

Our findings underscore the multiplicity of outcomes due to the combinatorial interplay between MTB lineages and macrophage phenotypes, which may have implications in proper design for host-directed therapies.

**IMPORTANCE:** The intricate interplay among host, pathogen, and environmental factors significantly contributes to susceptibility and clinical presentation of tuberculosis. These multifaceted mechanisms are often overlooked in tuberculosis studies, hindering our comprehensive understanding of infection progression and impeding the development of effective host-directed therapies which offer benefits such as reduced drug resistance emergence and heightened host compatibility.

Contrary to the prevailing hypothesis, the results observed in this study demonstrate the nuanced response of different *M. tuberculosis* lineages within distinct macrophage phenotypes in terms of phagosome acidification. The findings of our investigation delineate a crucial yet under-characterized aspect of tuberculosis pathogenesis, emphasizing how the interactions between different *M. tuberculosis* lineages and macrophage phenotypes could significantly influence the efficacy of host-directed therapies, particularly those targeting phagolysosomal maturation. More broadly, recognizing the impact of both *M. tuberculosis* and macrophage heterogeneity is paramount in the development of effective strategies to combat tuberculosis.

## INTRODUCTION

While significant progress has been made in understanding the pathogenesis of tuberculosis (TB), we still need to reach the disease elimination goal. *Mycobacterium tuberculosis* (MTB) resides within macrophages, crucial components for the formation of the granuloma and serving as survival niche for the bacteria. Within macrophages, MTB encounters various stresses and evolved strategies to cope with them. Among others, MTB has the capacity to inhibit phagosome maturation and block acidification of the compartment, modulates apoptosis, and inhibits reactive oxygen species production (1).

MTB has been considered a highly clonal organism and the laboratory strain H37Rv is largely used as a reference for research studies. However, the MTB complex (MTBC) consists of lineages with differences in transmission capacity, proliferation in the host tissues, ability to evade host immunity and in the propensity to acquire drug resistance (2, 3). More broadly, the distribution of lineages also reflects the reciprocal host-pathogen selective pressure leading to co-evolution and adaptation (4). The MTB complex (MTBC) includes seven main lineages grouped in two main phylogenetic clades, ancient (L1, L5, L6, L7) and modern (L2, L3, L4). Ancient lineages are geographically restricted and less virulent compared to modern ones and are referred as specialist strains (SPEC). Modern lineages are most virulent and geographically widespread, and for this reason are referred as generalist strains (GEN). The modern lineages 2 (Beijing), 3 (CAS/Delhi) and 4 (Haarlem, H37Rv) bear a large genomic deletion known as MTB-specific deletion 1 region (TbD1) and are associated with major TB epidemics, and high transmission rate compared to Lineage 1 (L1) (Indo-oceanic), L5 and L6 (*M. africanum*). Modern MTB strains, in general, induce an early inflammatory response which is weaker than those produced by ancient ones (3, 5).

Despite this microbe-centric classification, pathogenicity and virulence are nowadays recognized as functions of the host’s susceptibility and resistance (6). In this context, host heterogeneity at various levels should be taken into consideration. Besides genetic variability among individuals, human alveolar macrophages have been reported to show a mixed, plastic M1/M2 macrophage phenotype (7, 8). However, only a few studies considered the polarization status of the host’s macrophages in *in vitro*/*ex vivo* infection models (9).

In this study, we explore the MTB-human macrophage interactions at early time points post-infection (p.i.), considering both macrophage phenotypic variability and MTB genetic diversity. Single-cell approaches allowed a deep characterization of the heterogeneous host-pathogen interactions. Even small numbers of MTB bacilli can initiate infection *in vivo*, and at low multiplicities of infection *in vitro* only a subset of phagocytes become infected while others remain unexposed. These heterogeneous dynamics create distinct populations of infected and bystander cells, making single-cell analyses uniquely suited to interrogate processes such as phagosomal acidification with high precision (10, 11). We evaluated the inhibition of acidification in pathogen-containing compartments, autophagy, apoptosis, and necrosis in classically (M1) or alternatively (M2) activated THP-1-derived macrophage-like cells infected by different MTB lineages, as well as the survival rates of both macrophages and bacteria.

Our study suggests that the infection outcomes are strictly dependent on the interactions between MTB lineages and macrophage phenotypes which modulate phagosome acidification. In addition, our data suggests a broader remodeling of lysosomal pathways, potentially reflecting macrophage activation statuses or remodeling of the endolysosomal system dynamics. These results suggest that the MTB lineage plays a crucial role when considering host pathways. The impact of different MTB strains on phagosome acidification is particularly relevant for host-directed therapies, as treatments targeting endolysosomal pathways and phagolysosome maturation – such as metformin, imatinib, or pravastatin – may have varying efficacy depending on the strain tested. Relying solely on laboratory reference strains could therefore limit the reliability of such studies. This study lays the foundation for advancing host-pathogen interaction research and exploring alternative strategies for evaluating host-directed therapies.

## METHODS

### *M. tuberculosis* strains

Two clinical isolates representative of ancient (L1, L6) and modern lineages (L2, L4) and previously characterized were kindly provided by Prof. Stefan Niemann (Research Center Borstel, Borstel, Germany) (12). *M. tuberculosis* H37Rv NCTC-7416, *M. tuberculosis* H37Ra NCTC-7417, and *M. bovis* BCG NCTC-5692 were also included. Strains were classified and grouped according to the definitions provided in Stucki et al. related to the ecological niche width of MTB strains (Table 1) (13), resulting in ≥3 independent biological replicates per group, ensuring both single-cell resolution and robust biological replication. This grouping has been referred to as “geographical distribution and host adaptation” (geo-host adaptation) throughout the manuscript. Additional classifications are also provided for simplified comparison with other studies. All mycobacterial strains were engineered to express mCherry (Supplementary 1). For comparative infection experiments, we selected two representative strains each from L2, L3, and L6. Each of these strains was tested in at least two independent biological replicates, defined as separate experiments using independently grown bacterial cultures. In parallel, reference strains including H37Rv, H37Ra, BCG, and CDC1551 were included as controls and tested across at least three independent experiments using separate cultures; as these strains are genetically identical across replicates, they are considered biological replicates of the same strain.

**Table 1:** Laboratory and clinical MTB strains considered in this study and classification used for the analysis of the results.

| Strain<sup>a</sup> | Lineage nomenclature | Grouping geo-host adaptation<sup>b</sup> | Lineage type | Virulence | Laboratory adaptation | Assay |  |  |  |  |
| --- | --- | --- | --- | --- | --- | --- | --- | --- | --- | --- |
|  |  |  |  |  |  | Phagolysosomal acidification | Autophagy<sup>c</sup> | Apoptosis<sup>c</sup> | Macrophage survival | Mycobacterial survival |
| *M. africanum* 10514/01<sup>d</sup> | 6 (Africanum) | Specialist (SPEC) | ancient | Less | Clinical | y | y | y | y | y |
| *M. africanum* 5468/02<sup>d</sup> | 6 (Africanum) | SPEC | ancient | Less | Clinical | y | y | y | y | y |
| *M. bovis* BCG | Bovis | Other (OTH) | animal | Less | Laboratory | y | y | n | y | n |
| *M. tuberculosis* 12594/02<sup>d</sup> | 2 (Beijing) | Generalist (GEN) | modern | More | Clinical | y | y | y | y | y |
| *M. tuberculosis* 1934/03<sup>d</sup> | 2 (Beijing) | GEN | modern | More | Clinical | y | y | y | y | y |
| *M. tuberculosis* CDC1551<sup>d</sup> | 4 | Intermediate (INT) | modern | More | Clinical | y | n | n | y | n |
| *M. tuberculosis* 1797/03<sup>d</sup> | 1 | SPEC | ancient | Less | Clinical | y | n | n | n | n |
| *M. tuberculosis* 4850/03<sup>d</sup> | 1 | SPEC | ancient | Less | Clinical | y | n | n | n | n |
| *M. tuberculosis* H37Ra NCTC 7417 | 4 | OTH | modern | Less | Laboratory | y | y | n | y | y |
| *M. tuberculosis* H37Rv NCTC 7416 | 4 | GEN | modern | More | Laboratory | y | y | y | y | y |

Assay performed: y (yes) n (no). <sup>a</sup> L1, L6, L2 and L4 clinical isolates from (12); <sup>b</sup> grouping according to (13); <sup>c</sup> provided as Supplementary 4 and 5. <sup>d</sup>
strains from (12).

### Cells preparation and infection

Macrophage-like cells were derived from THP-1 as previously described (14) (Supplementary 1). Given the lack of standard protocols for recapitulating the alveolar macrophage *in vitro* cell lines, we opted to consider the two extremes of the dynamic state of activation of the macrophage. Three days after removal of the PMA, macrophage-like cells were polarized in cRPMI medium without antibiotics for 16h with either lipopolysaccharide (50 ng/mL, Sigma-Aldrich, Missouri, USA) to induce an M1-like phenotype [M(LPS)] or human IL-4 [20 ng/mL, *E. coli*-derived human IL-4 protein (R&Dsystems, Minnesota, USA)] to induce an M2a-like phenotype [M(IL-4)] prior to the infection (14–16). Effective polarization of cells was evaluated by checking morphology and the expression of specific markers for M1 and M2 (TNFα, IL-1β, CD80, IL-6, CD206) (Figure S1). THP-1 derived macrophages were infected with mycobacteria at a theoretical multiplicity of infection (MOI) 10:1 for achieving robust and efficient infection, thus ensuring that most target cells are exposed to bacteria (Supplementary 1).

### Live imaging microscopy

All experiments were carried out using a Leica TCS SP5 confocal microscope equipped with an incubator chamber allowing to stably maintain cells at 37 °C in 5% CO_2_. Images were acquired as 16-bit, 1024×1024 resolution, TIFF files using a 63x, 1.40 NA immersion oil objective.

#### Phagosome acidification, live imaging

To assess acidification dynamics in infected macrophages, we employed LysoSensor staining combined with confocal microscopy and *z-*stack acquisition. Intracellular acidification was quantified using a size-filtered ROI mask to identify LysoSensor-positive structures, thereby excluding background signal and capturing a defined range of acidic compartments (Supplementary 1). For clarity, these LysoSensor-positive structures are hereafter referred to as acidified compartments, encompassing large lysosomes as well as phagolysosomes. To specifically assess phagosomal acidification, we performed colocalization analysis between LysoSensor fluorescence and internalized bacteria, allowing us to distinguish pathogen-associated acidified compartments from the broader lysosomal population (see Supplementary 1 for details). M1 and M2 macrophages were stained with Hoechst 33342 NucBlue^TM^ Live ReadyProbes^TM^ Reagent (Ex/Em 360-460 nm, ThermoFisher, Massachussets, USA) for nuclei, and with LysoSensor™ Green DND-189 (Ex/Em 443-505 nm, pK_a_ 5.2, ThermoFisher) for monitoring dynamics of acidified compartments and phagosome acidification according to manufacturer’s instructions. Imaging experiments were performed at 6- and 24-hours post-infection (p.i.). For each strain, 6 images (2 positions for each well out of 3 wells per strain) were acquired as *z-*stacks (0.89 µm x 7 stacks) in at least two independent experiments. All images have been acquired and analyzed using the same settings for allowing quantitative comparison of LysoSensor intensities (Supplementary 2). Biological and technical replicates are detailed in Supplementary 2. Induction of autophagosomes, apoptosis and necrosis in live imaging are provided as Supplementary 4 and 5, respectively.

### Data analysis

Images were processed for single-cell analysis with Fiji software(17). Stacks have been analyzed by applying the maximum intensity projection method. For each infected well, at least 5 cells with internalized mycobacteria and 5 without internalized (unless not available) were considered for each acquisition. Infected wells were compared toward “uninfected controls” (CTRL) defined as samples unexposed mycobacterial infection. Effective cell infection was visually inspected using orthogonal views. Single cells were manually segmented (Supplementary 1).

All images have been acquired and analyzed using the settings as reported in Supplementary 1. The Mann–Whitney test was performed to compare two independent groups, while in presence of more than two independent groups the Kruskal–Wallis test followed by post hoc analysis using Dunn’s test was used. In presence of dependent observations, the linear/logistic mixed-effects models (18, 19), linear quantile mixed models (20), or Zero-inflated Mixed-effects Poisson models (21) were performed (Supplementary 1). Inferential techniques were applied in presence of adequate sample sizes (n ≥ 5). For all analyses, the significance threshold was set at 0.05. Analyses were performed using R statistical software (22). Full statistics reports are available upon request.

### Macrophage survival

THP-1 were plated and differentiated in 96well plates (55,000 cells/well) as described above (M1 or M2) and *in vitro* infected with MTB strains at MOI 5:1 to ensure macrophage infection but reducing the risk of overwhelming the cells or triggering too much cell death. MTB strains used for this experiment include the laboratory strains H37Rv, H37Ra, *M. bovis* BCG NCTC-5692, CDC1551 and clinical strains 2x L2, 2x L6 (Table 1). After 48h and 5 days p.i., cells viability was assessed using the crystal violet assay as previously described (23) (Supplementary 1). The percentage of cell death compared to the uninfected control was considered.

### Intramacrophage MTB survival

For mycobacterial survival, cells were lysed with sterile water. Supernatants and cell lysate were serially diluted, and each dilution was plated on 7H10 agar plate to count MTB CFUs present at 4h, and 48h p.i.. After 3 weeks of incubation, colonies were counted. CFUs/mL obtained at 4h p.i. (macrophage entry) were then normalized on the CFU obtained from the inoculum, while the data obtained at 24 and 48h p.i. (survival capacity) were normalized on the CFUs/mL values obtained at 4h p.i. Analyses were performed using Prism-GraphPad software.

## RESULTS

The activity of acidifying drugs jointly used with antibiotics impair MTB survival (24). Phagosome acidifying mechanisms are cell-autonomous, therefore strictly dependent on the phenotype of both the host and the infecting MTB strain and their interactions. Therefore, we aimed at exploring the blocking ability of different strains in both M1 and M2 macrophages. LysoSensor Green stains acidic compartments broadly; therefore, we refer to “acidified compartments” throughout. To identify pathogen-containing acidified compartments we used colocalization between LysoSensor signal and internalized bacteria; this colocalization does not necessarily prove canonical phagolysosome identity (see Methods and Discussion).

Findings are described using the grouping generalists/specialists, as this classification encompasses different features such as phylogeny, geographical distribution and host adaptation. Phagosome acidification results for ungrouped samples are reported in Supplementary 2.

### Phagosomal acidification

LysoSensor Green was used to quantify the abundance and intensity of acidified compartments in THP-1-derived M1 and M2 macrophages infected with generalist (GEN), intermediate (INT), specialist (SPEC) and other (OTH) strains (Table 1) at 6 and 24h post infection (p.i.) (Figure 1). To specifically assess phagosomal acidification, LysoSensor fluorescence was analyzed in conjunction with intracellular bacterial localization. A total of 7384 single cells (4098 at 6h p.i.; 3286 at 24h p.i.) are considered for the analysis (on average, for each strain tested, M1: 178±67 uninfected cells + 172±44 infected cells; M2: 180±65 uninfected cells + 174±46 infected cells) (Supplementary 2). Bystander effect refers to a phenomenon where uninfected cells exhibit changes or behaviors typically associated with infected cells, due to signals or factors released by the infected cells. This means that even though some cells are not directly infected by MTB, they still show similar features as the infected cells because of their proximity or response to the infected cells’ influence. In M1 macrophages, GEN and SPEC strains do not block the LysoSensor-based acidification signal after 6h p.i., with the latter inducing also a bystander effect (increase of acidification in adjacent, uninfected macrophages) in M1 macrophages, although not statistically significant (Figure 2a). In contrast, the categories INT and OTH, display a reduction of acidification levels in large LysoSensor-positive compartments, suggesting interference with host endolysosomal dynamics (Figure 2a). On the contrary, the SPEC strains at 6h p.i. are the less efficient in blocking the acidification and induce the highest number of acidified macrophages (50% vs <25% of all the other categories) (Figure 2b). At 24h p.i., acidification levels are overall high, irrespective of the infecting MTB lineage (Figure 2c). However, at single cell level, only a small macrophage fraction displays such acidification (<25%) (Figure 2d). Analysis on the number of LysoSensor-positive compartments revealed a time-dependent shift in M1 macrophages infected with GEN versus SPEC strains. At 6h p.i., GEN-infected macrophages displayed significantly fewer acidified compartments than SPEC, whereas at 24h p.i., GEN-infected macrophages displayed significantly more acidified compartments, indicating a group-specific temporal modulation of phagosomal acidification (Supplementary 2). In M2 macrophages, none of the categories block the acidification of LysoSensor-positive compartments at 6h p.i. (Figure 3a), and the fraction of acidified macrophage ranges between 25% (OTH) and 50% (SPEC and GEN) (Figure 3b). The SPEC strains induce a bystander effect in uninfected M2 macrophages (approximately 40% of the SPEC uninfected cells show acidification; Figure 3a-3b). As the time of infection progresses (24h p.i.), GEN and INT strains block the acidification of compartments (Figure 3c) and the fraction of acidified macrophages decreases below 25%, except for the OTH category (50%) (Figure 3d). No significant differences were observed in the number of LysoSensor-positive compartments across the different MTB groups (Supplementary 2).

**Figure 1:**
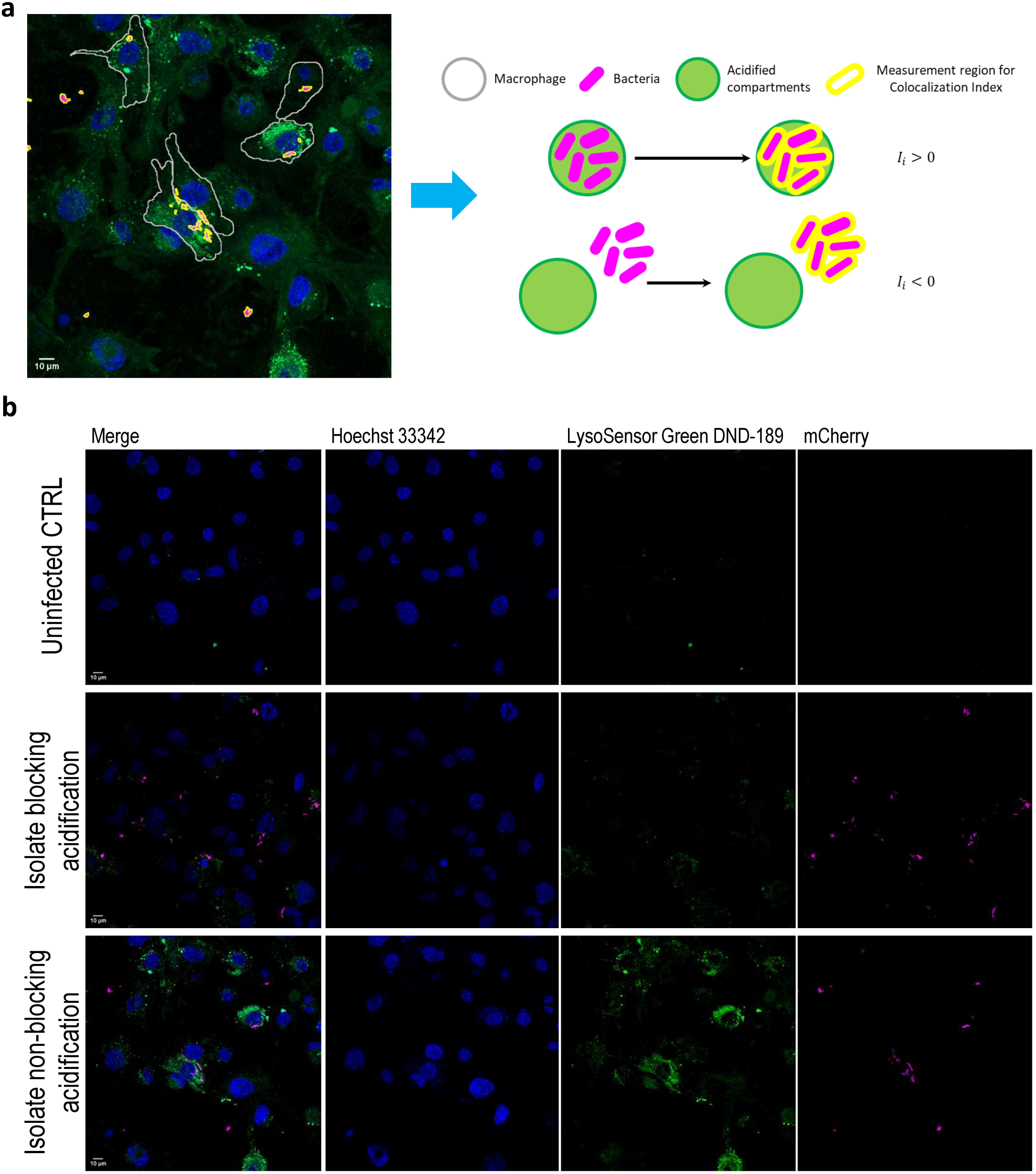
Phagosomal acidification in M1 cells. (a) Schematic representation of the elements considered for the analysis (see section “Images and data analysis - Phagolysosomal acidification” in Materials and Methods for details). (b) representative images of uninfected control macrophages (no exposure to mycobacteria, top), macrophages infected with isolate blocking phagosomal acidification (middle), and macrophages infected with isolate non-blocking phagosomal acidification (bottom); the fluorescence images were acquired in live imaging after staining with LysoSensor (green) and Hoechst (blue). Bacteria are reported in magenta (mCherry). Scale bar: 10µm.

**Figure 2:**
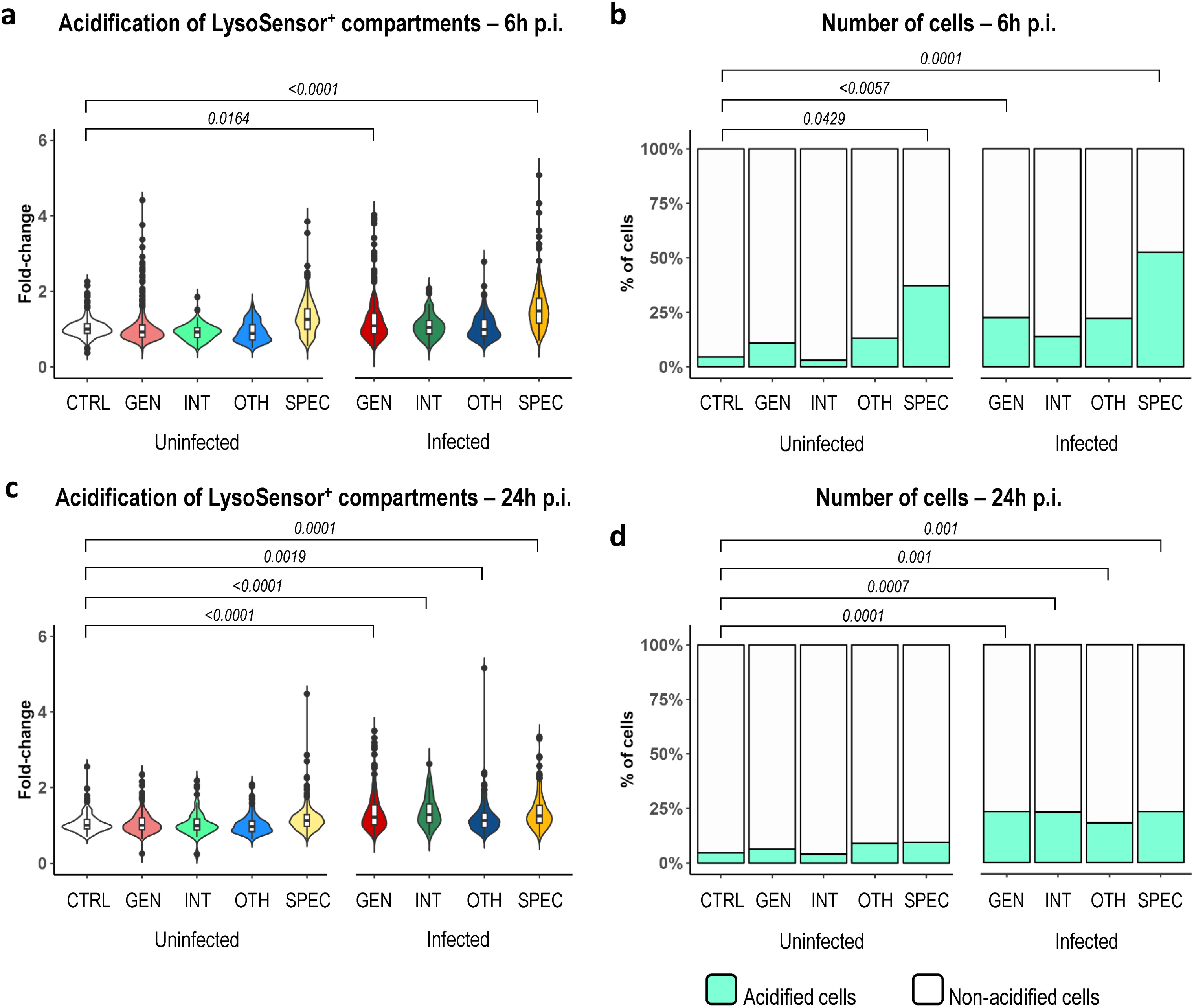
Phagosomal acidification in M1 cells. Phagosomal acidification at 6h p.i. (a) and at 24h p.i. (c). Fold-changes are visualized as a violin plot to show the distribution, with a superimposed whisker plot (boxplot) representing the median, interquartile range, and potential outliers. Data are normalized on M1 THP-1 macrophages uninfected control and strains are grouped as previously reported in Table 1. Linear mixed-effects models followed by *post hoc* analysis. Percentage of cells showing phagosomal acidification at 6h p.i. (b) and at 24h p.i. (d) is also reported. Logistic mixed-effects models followed by *post hoc* analysis. GEN: Generalist strains *M. tuberculosis* 12594/02 (L2)(12), *M. tuberculosis* 1934/03 (L2)(12), *M. tuberculosis* H37Rv NCTC 7416 (L4); INT: Intermediate strain *M. tuberculosis* CDC1551(12); OTH: Other strains *M. tuberculosis* H37Ra NCTC 7417, *M. bovis* BCG; SPEC: Specialist strains *M. africanum* 10514/01 (L6)(12), *M. africanum* 5468/02 (L6)(12), *M. tuberculosis* 1797/03 (L1)(12), *M. tuberculosis* 4850/03 (L1)(12). p-values: comparison between uninfected controls and each category. Uninfected CTRL = no exposure to mycobacteria; Uninfected GEN/INT/OTH/SPEC = exposure to mycobacteria but no uptake observed; Infected = intracellular mycobacteria observed. Secondary analysis comparing M1 vs M2 is available in Supplementary 3.

**Figure 3:**
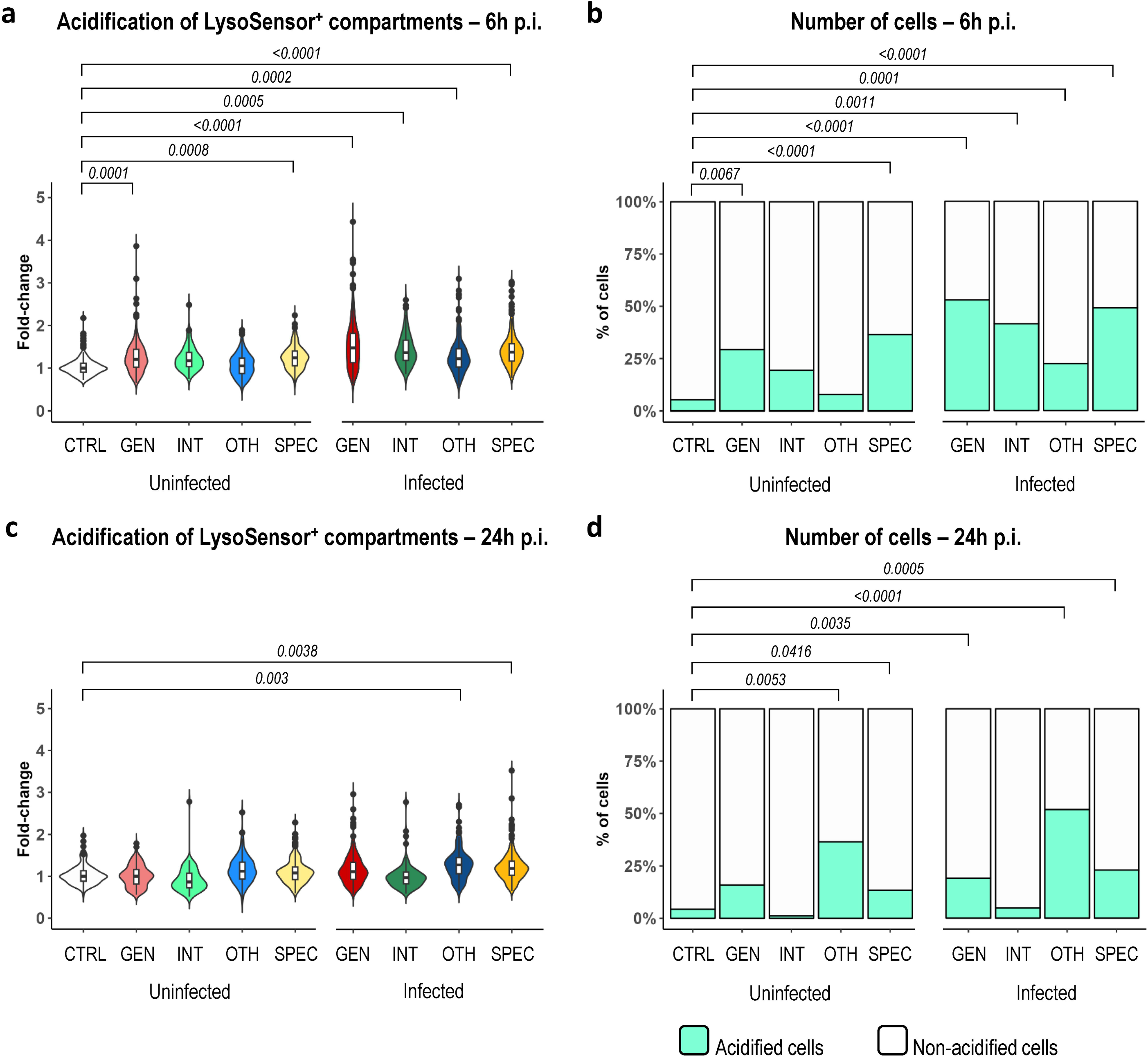
Phagosomal acidification in M2 cells. Phagosomal acidification at 6h p.i. (a) and at 24h p.i. (c). Fold-changes are visualized as a violin plot to show the distribution, with a superimposed whisker plot (boxplot) representing the median, interquartile range, and potential outliers. Data are normalized on M2 THP-1 macrophages uninfected control (CTRL) and strains are grouped as previously reported in Table 1. Linear mixed-effects models followed by *post hoc* analysis. Percentage of cells showing phagosomal acidification at 6h p.i. (b) and at 24h p.i. (d) is also reported. Logistic mixed-effects models followed by *post hoc* analysis. GEN: Generalist strains *M. tuberculosis* 12594/02 (L2)(12), *M. tuberculosis* 1934/03 (L2)(12), *M. tuberculosis* H37Rv NCTC 7416 (L4); INT: Intermediate strain *M. tuberculosis* CDC1551(12); OTH: Other strains *M. tuberculosis* H37Ra NCTC 7417, *M. bovis* BCG; SPEC: Specialist strains *M. africanum* 10514/01 (L6)(12), *M. africanum* 5468/02 (L6)(12), *M. tuberculosis* 1797/03 (L1)(12), *M. tuberculosis* 4850/03 (L1)(12). p-values: comparison between uninfected controls and each category. Uninfected CTRL = no exposure to mycobacteria; Uninfected GEN/INT/OTH/SPEC = exposure to mycobacteria but no uptake observed; Infected = intracellular mycobacteria observed. Secondary analysis comparing M1 vs M2 is available in Supplementary 3.

We then evaluated the replicative potential of the MTB strains attenuated by the killing in M1 and M2 macrophages expressed as number of mycobacteria internalized per cell (M/C).

All the strains infect M1 macrophages with comparable efficiency as M/C values at 6h p.i. are not significantly different (Figure 4a). MTB colocalization inside acidified compartments is similar among all the groups (Figure 4b). Comparison of M/C values at 6h p.i. *vs* 24h p.i. within the same group highlights some differences in proliferation: GEN, INT and OTH show increased M/C at 24h p.i. (p-values 0.0139, 0.0011, and <0.0001 respectively), whereas SPEC strains display no significant differences between the timepoints considered (p-value 0.7748) (Supplementary 3). SPEC strains show the lowest M/C values compared to the other groups, suggesting an impaired proliferation or a higher killing (Figure 4a). The MTB-phagosome colocalization index within acidified compartments is lower for SPEC and INT strains compared to the GEN strains (Figure 4b).

**Figure 4:**
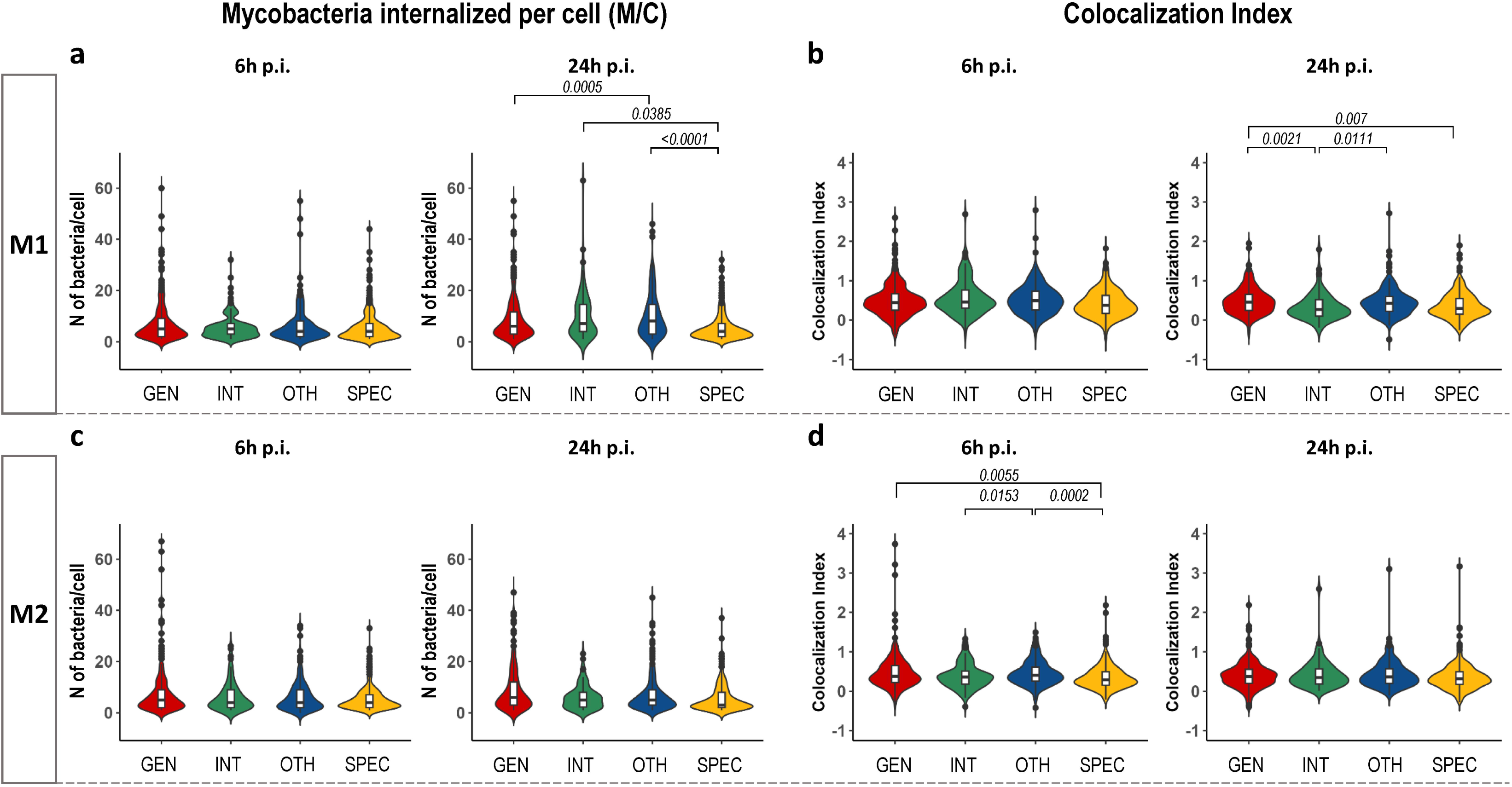
a) Effective number of mycobacteria internalized per cell (M/C) at 6h p.i and at 24h p.i. in M1 cells. b) MTB colocalization with acidified compartments at 6h p.i. and 24h p.i. in M1 cells.(c) Effective number of mycobacteria internalized per cell (M/C) at 6h p.i and at 24h p.i. in M2 cells.(d) MTB colocalization with acidified compartments at 6h p.i. and 24h p.i. in M2 cells. Linear mixed-effects models followed by *post hoc* analysis. GEN: Generalist strains *M. tuberculosis* 12594/02 (L2)(12), *M. tuberculosis* 1934/03 (L2)(12), *M. tuberculosis* H37Rv NCTC 7416 (L4); INT: Intermediate strain *M. tuberculosis* CDC1551(12); OTH: Other strains *M. tuberculosis* H37Ra NCTC 7417, *M. bovis* BCG; SPEC: Specialist strains *M. africanum* 10514/01 (L6)(12), *M. africanum* 5468/02 (L6)(12), *M. tuberculosis* 1797/03 (L1)(12), *M. tuberculosis* 4850/03 (L1)(12). Secondary analysis comparing M1 vs M2 is available in Supplementary 3.

In M2 macrophages, all the strains infect and survive with similar rates even though GEN strains display a heavy-tail distribution due to cells with very high M/C values (Figure 4c). INT and SPEC strains show a reduced colocalization with acidified compartments at 6h p.i as compared to GEN strains which is abrogated at 24h p.i. (Figure 4d). Comparison of M/C values at 6h p.i. *vs* 24h p.i. within the same group highlights differences in proliferation only for GEN OTH (p-values 0.001, and 0.0442 respectively), whereas SPEC and INT strains display no significant differences between the timepoints considered (p-values 0.1501, and 0.6585 respectively) (Supplementary 3). No correlation between M/C and the colocalization index is found for the described time points (data not shown).

Together these findings indicate that the ability of phagosome acidification to halt MTB growth is triggered by all the strains in M2 macrophages soon after infection and is unpaired in GEN and INT-infected cells upon infection progression. Conversely, in M1 macrophages, complete phagosome acidification occurs later but with greater efficiency. This enhanced efficiency is observed across all the isolates examined. Nonetheless, the single-cell approach revealed the heterogeneity of the response in the macrophage population indicating that only 25% of infected cells is still capable to elicit phagosomal acidification and, possibly, protect the cells. Furthermore, SPEC isolates induce a bystander effect in adjacent, uninfected macrophages. As suggested by the analysis of the number of LysoSensor-positive compartments (Supplementary 2), the bystander increase in acidification in uninfected macrophages appears to reflect a true enhancement of phagosomal acidification per compartment rather than an increase in compartment number. It is worth mentioning that M1 and M2 macrophages show a different acidification response timing to MTB infections suggesting an intrinsic flexibility and adaptability of the protective mechanisms.

To further investigate the heterogeneity of macrophage responses, we applied linear quantile mixed models (LQMM) and logistic mixed-effects models to perform comparison between M1 and M2 (Supplementary 3). LQMM showed no significant differences in median acidification values between M1 and M2, indicating similar overall levels. In contrast, logistic mixed-effects models identified three significant differences: (i) the percentage of cells showing acidification and (ii) the colocalization index at 6h p.i., and (iii) the number of mycobacteria per cell at 24h p.i. These findings suggest that, while median acidification is comparable, the responses of individual cells differ in terms of the proportion of responders, timing, and infection outcome. Group-specific analyses revealed that these differences are primarily driven by the GEN (non-infected and infected at 6h p.i.) and the INT (infected at 24h p.i.) groups, whereas other macrophage subgroups showed comparable responses (Supplementary 3).

### Autophagy and apoptosis

Although our primary focus was on acidification, we also explored additional host responses, including autophagy, apoptosis and necrosis are provided as Supplementary 4 and 5, respectively. Briefly, none of the strain groups induced effective autophagy: although autophagosome number increased, their size did not show a significant enlargement, suggesting impaired maturation and limited progression of the autophagic flux (Supplementary 4). This pattern was consistent in both M1 and M2 macrophages. Apoptosis was more pronounced in M1 macrophages, particularly in bystander non-infected cells exposed to GEN and OTH strains (∼40%). A similar, though not statistically significant, trend was observed in infected cells. In M2 macrophages, only bystander cells exposed to OTH strains showed a modest, non-significant increase in apoptosis (∼30%) (Supplementary 5).

### Macrophage survival

To quantify the impact of MTB infections on macrophage survival, we compared cell viability after infection to viability on uninfected cells. Figure 5a shows viability data at 48h p.i. indicating that some strains decrease the viability by 20%, whereas others by 50% in M1 cells. At the same time point, all the strains are less aggressive on M2 cells (Figure 5b). As the infection progresses for 5 days, M1 cell viability for all the infected populations is 50% as compared to the uninfected populations (Figure 5c). Conversely, M2 macrophages show a dramatic viability decrease (80%) for all the infecting MTB categories (Figure 5d).

**Figure 5:**
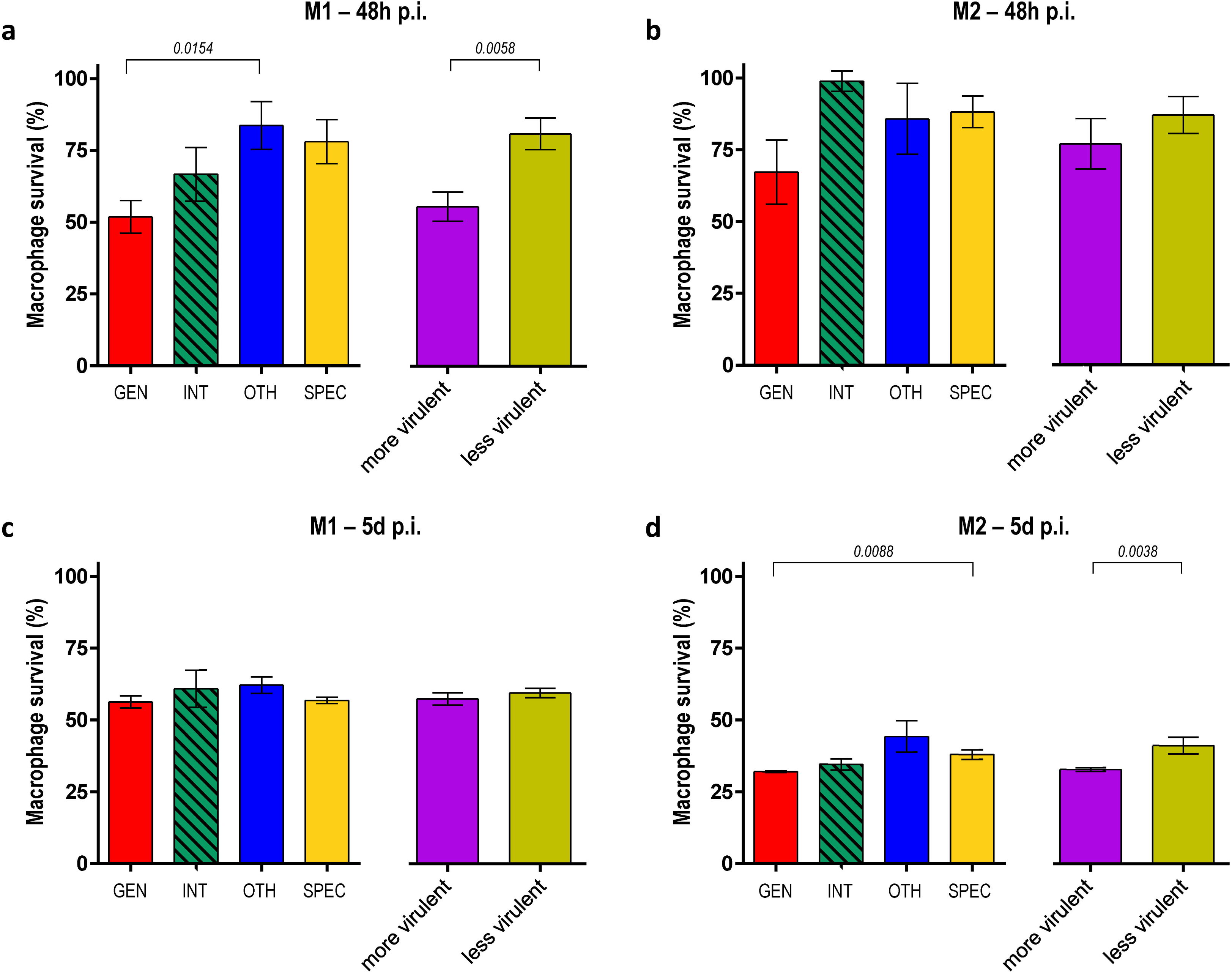
Cell survival at 48h and 5 days post-infection. Percentage of M1 survival at 48h (a) and at 5 days (c) post-infection. Percentage of M2 survival at 48h (b) and at 5 days (d) post-infection. Data (mean ± standard error of the mean) are normalized on uninfected controls and strains are grouped as previously reported in Table 1. Kruskal-Wallis and Mann–Whitney tests. INT group is reported for comparison but excluded from the statistical model (n < 5, see Materials and Methods). GEN: Generalist strains *M. tuberculosis* 12594/02 (L2)(12), *M. tuberculosis* 1934/03 (L2)(12), *M. tuberculosis* H37Rv NCTC 7416 (L4); INT: Intermediate strain *M. tuberculosis* CDC1551(12); OTH: Other strains *M. tuberculosis* H37Ra NCTC 7417, *M. bovis* BCG; SPEC: Specialist strains *M. africanum* 10514/01 (L6)(12), *M. africanum* 5468/02 (L6)(12), *M. tuberculosis* 1797/03 (L1)(12), *M. tuberculosis* 4850/03 (L1)(12).

By clustering the tested strains according to their virulence (see Table 1), it is noticeable that highly virulent strains reduce M1 cell viability by 50% within 48 hours p.i., while low virulent strains are less effective at killing M1 macrophages even at 5 days p.i. However, this difference in virulence appears mitigated in M2 cells. Highly virulent strains reduce M2 cell viability by 80% between 3 and 5 days, while low virulent strains manage to kill nearly 70% of M2 cells by 5 days p.i.

### Intramacrophage MTB survival

Colony Forming Unit (CFU) assays are performed to quantify both the efficiency of the different strains to enter M1 and M2 macrophage cells (CFU at 4h p.i., Supplementary 6) and the MTB survival potential (CFU at 48h p.i.). Briefly, macrophages are exposed to a constant number of MTB cells. After 4h or 48h, macrophages are washed to remove non phagocytosed bacteria, lysed and contained-MTB cells plated on agar plates. To calculate the MTB replicative potential, CFU values at 48h p.i. are normalized on those at 4h p.i. thus expressing the MTB ability to replicate intracellularly. H37Rv and H37Ra show similar low replicative potential, while clinical strains grow faster in both M1 and M2 macrophages. L2 strains show an initial lower uptake in M1 cells, with fast growth rate (Fig. 6). These findings suggest that M2 macrophages are more easily infected by MTB, and that clinically-adapted isolates can replicate more efficiently than laboratory-adapted strains in the macrophages.

**Figure 6:**
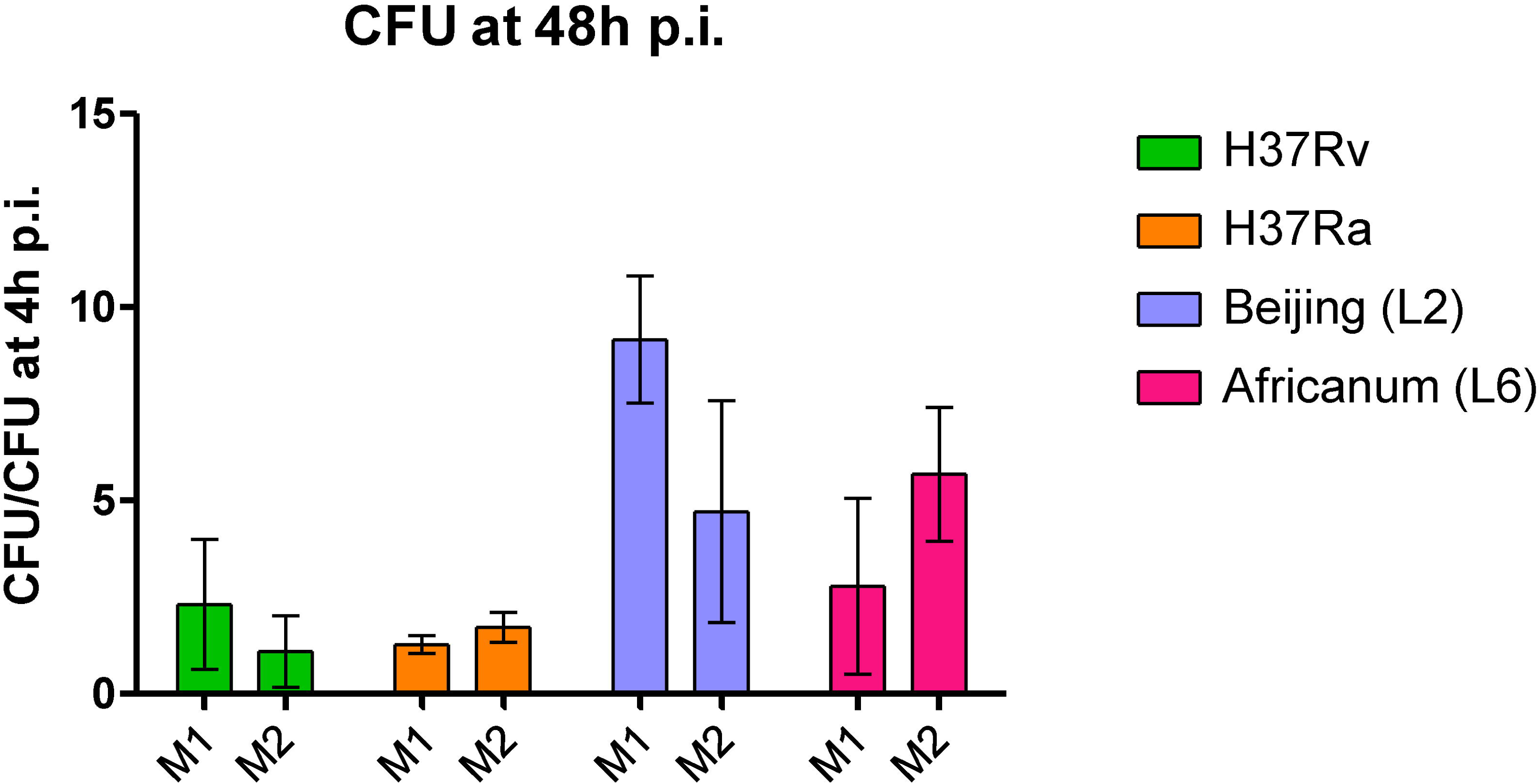
MTB intramacrophage survival at 48h post-infection. CFU results at at 48h p.i. Data (mean ± standard error of the mean) are normalized on CFU at 4h p.i.. Beijing: GEN/L2 (*M. tuberculosis* 12594/02(12), *M. tuberculosis* 1934/03(12)); Africanum: SPEC/L6 (*M. africanum* 10514/01(12), *M. africanum* 5468/02(12)). Statistical model not applied (n < 5, see Materials and Methods).

## DISCUSSION

Our single-cell based study shows that both MTB lineages and macrophage differentiation status critically steer the cell-autonomous defensive mechanisms deployed upon MTB infection. To our knowledge, our study is among the first to integrate both host immunophenotype heterogeneity and *M. tuberculosis* genetic diversity within a single experimental setting. The use of single-cell based analyses represents an additional novelty relevant to comprehend the heterogeneity of the host-pathogen interaction.

On a mechanistic level, our results contribute to a better understanding of MTB survival within macrophages, pathogenesis and virulence. On a clinical level, our results suggest a way to optimize therapeutic protocols already in use for mycobacterial infections (25). While the magnitude of differences in our single-cell data appears small, such shifts can still have significant biological implications. In single-cell studies, even subtle changes, such as early cellular transitions, rare subpopulation dynamics, may reflect downstream amplification effects. These results provide insights into cellular heterogeneity and the underlying biology, warranting further investigation.

### MTB lineages calibrate phagosome acidification response unique to macrophage phenotypes

The central role of phagosomal acidification is highlighted by the fact that most permissive monocytes display less lysosome content, acidification, and proteolytic activity and enable MTB in vivo persistence (26, 27). Recently, Bedard et al. contributed to the field describing how lysosomal failure is critical for MTB (H37Rv) survival, and how drug restoring lysosomal function can counteract MTB survival (28).

Our study provides evidence that different MTB lineages in different macrophage phenotypes cope with phagosomal acidification in macrophages in lineage- and phenotype-specific ways (illustrated in Figure S2). In M1 macrophages, the higher level of acidification together with the concomitant higher colocalization of modern, virulent GEN/INT isolates, suggests better tolerance to the acidic environment. This interpretation is further supported by our observation that, in M1 macrophages, GEN strains transiently limit early phagosomal acidification but ultimately sustain a larger pool of acidified compartments per cell, providing an expanded interface for host-pathogen interactions. In contrast, SPEC strains retain more acidified compartments early on but show no further increase, suggesting a distinct intracellular survival strategy. Consistent with this view, L2 strains evolved specific strategies to cope with acidification during intramacrophage infection (29–32). In contrast, L1, L5 and L6 (*M. africanum*) isolates harbor mutations negatively affecting the adaptive response to acidic pH and lysosomal delivery(33, 34). Despite recent findings showing no significant differences in mycobacterial survival of modern and L1 isolates in acidic pH *in vitro*, the transcriptional regulatory network of ancient and modern lineages is obviously distinct, with crucial implications for stress response *in vivo* (30).

Differently, the weak M2 macrophage antimicrobial defenses are quickly hijacked by modern/virulent GEN strains which block phagosomal acidification. Whereas ancient/less virulent SPEC strains seem not to affect phagosomal acidification itself, colocalization analysis at the time points considered suggests they escape from acidified compartments(35). Consistent with this view, we observed no significant differences in the number of LysoSensor-positive compartments across MTB groups, indicating that phagosomal acidification per cell is comparable in this macrophage phenotype. It is well known that less virulent strains trigger an apoptotic bystander effect on uninfected M2 macrophages (36, 37). For the first time, we report here a similar bystander effect with increased acidification in M2 macrophages, underscoring the critical role of macrophage polarization in conditioning the microenvironment (38). Our observation that uninfected bystander macrophages show increased acidification suggests a functional amplification of host defense extending beyond directly infected cells. This response may be mediated by local signalling from infected macrophages, priming neighbouring cells for more efficient antigen processing and antimicrobial activity. By enhancing the processing of secreted mycobacterial antigens and facilitating antigen presentation, such a mechanism could strengthen the overall capacity of the macrophage population to control infection. Furthermore, even if the infection prompts the phagosomal acidification, the number of M1 or M2 macrophages triggering the signal remains relatively low.

### Macrophage phenotype forebodes cell survival and MTB intracellular replication

Finally, we assessed strains’ ability to enter, replicate in host cells, and kill macrophages as an extreme measure of virulence (considered as “ability to cause damage”). Compared to SPEC strains, GEN strains showed higher replicative potential in both M1 and M2 macrophages. Furthermore, M/C data trends also suggest an increased uptake of GEN strains in M2 macrophages. Survival rate strongly depends on the strain-specific virulence in M1 cells, while M2 are inherently more sensitive to the infection *per se* with no contribution of strain identity.

These findings are aligned with the microscopy observations for apoptosis, where GENs/more virulent strains induced higher cell death mainly in M1 macrophages. Of note, M2 macrophages result more permissive to MTB growth compared to M1, an outcome partially due to the lower MTB intake by M1 macrophages in comparison with M2. Based on these results we may imagine that in an *in vivo* scenario, the interplay between the specific MTB lineage and the immune environment where all the other immune cells are included (i.e. T cells, Natural Killer cells, MAIT cells) and potential coinfection or comorbidities may have a tremendous impact on the immune response, containment of the mycobacteria and the clinical outcome.

Suitable models of alveolar macrophages to dissect host-pathogen interactions in TB, are currently missing being the Max Planck Institute mouse cells or the human peripheral blood mononuclear cells poorly validated (39, 40). We used PMA-differentiated THP-1 cells to reduce donor-to-donor variability inherent to primary cell models (41–43). In addition, favouring a “pre-programming” mode as opposed to a “reprogramming” mode, we avoided to recapitulate features of tissue niche and cell origin contributions which might lead to a basic understanding of the whole processes (44–47). Increasing the number of MTB isolates considered might be also needed. However, THP-1 derived macrophages do not fully recapitulate human alveolar macrophage physiology. Therefore, while THP-1 cells provide an internally consistent platform for comparing MTB lineages, extrapolation to alveolar macrophages in vivo should be made with caution and will require confirmation in primary human alveolar macrophages or *in vivo* models. Acidified compartments were identified using LysoSensor; future studies including organelle-specific markers (e.g. LAMP1) could provide additional resolution on the identity of MTB-containing compartments and to directly assess lysosome biogenesis or distinguish whether observed changes reflect altered lysosomal number versus acidification levels. Future work incorporating lysosomal biogenesis markers and functional assays would help clarify these mechanisms. While CYTO-ID staining provides an overview of autophagic vesicle accumulation, it does not allow mechanistic dissection of the autophagic process. As such, our data indicates no evidence of completed autophagic flux based on CYTO-ID, but further analyses (e.g. LC3/p62 turnover assays, with or without bafilomycin) would be required to define the underlying mechanisms. Finally, we did not test the effect of drugs such as metformin, cysteamine imatinib or pravastatin (24).

In conclusion, our study underlines the role of the genetic diversity of MTBC strains in the pathogenesis of TB and in relationship with defined host conditions. While acknowledging that *in vivo* infection dynamics evolve over time, our *in vitro* findings reinforce the role of macrophage polarization in shaping intracellular stress conditions encountered by MTB and remains an underestimated component of *in vitro/ex vivo* infection models. Alveolar macrophages (resembled by M2 polarization in this study based on previous evidence (48, 49)) are more permissive to infection, and MTB is subjected to less stress and fitness costs; on the other side, in interstitial recruited macrophages (mirrored by M1 polarization in our study) MTB is challenged with less permissive conditions (50, 51). These results align with previous studies conducted in human models. We recognize that murine studies have recently shown a shift in MTB cellular reservoirs during chronic infection stages, with monocyte-derived macrophages becoming dominant (27, 52).

Whereas previous studies showed a slower mycobacterial replication rate in interstitial recruited macrophages compared to alveolar macrophages, our data provide evidence that this feature is further influenced by the mycobacterial lineage, which affects both the cell entry rate and intra-macrophage replication. Collectively, our data supports the view that reliance solely on reference laboratory strains or static macrophage models may not fully recapitulate the dynamic complexity of TB pathogenesis.

The use of a single-cell analytical approach highlighted the heterogeneity of the macrophage response to *M. tuberculosis* infection. Although linear quantile mixed models (LQMM) showed no significant differences in median acidification values between M1 and M2, logistic mixed-effects models identified three significant differences: (i) the percentage of cells showing acidification and (ii) the colocalization index, and (iii) the number of mycobacteria per cell. These results indicate that, while the average acidification level is similar between M1 and M2, the responses of individual cells differ in terms of responder proportion, timing, and infection outcome. Group-specific analyses further revealed that these differences are primarily driven by the INT and GEN groups. These findings underscore the heterogeneity of macrophage responses and highlight the importance of considering both host cell type and infection status in evaluating MTB-macrophage interactions, further emphasizing the value of single-cell approaches in detecting subtle but biologically meaningful heterogeneity. While a general increase in phagosomal acidification may be observed, our data suggest that this increment is mainly due to a relatively small proportion of cells rather than being a population-average response and are aligned with previous similar findings (27, 53). Furthermore, the differences in acidification time could be due to fluctuations in the levels of MTB damaging (virulence) factors, as already hypothesized (54). This highlights the need for single-cell methods capable of discriminating signaling responses driven by infection. In recent years, host-directed therapy targeting the pathogen-induced immune modulation represents a promising alternative (or at least adjunct) therapy, with several benefits over antibiotic treatment alone, including the reduction of emergence of drug resistance and increased host compatibility (55). Our findings might have implications for host-directed therapies, particularly those targeting phagolysosomal maturation (56). Therefore, it is critical to consider MTB and macrophage heterogeneity in exploring host-directed therapeutics. In addition, our findings might even impact the effects of standard anti-tubercular drugs on phagosome maturation and autophagy (24, 25). Future studies are needed to identify individual factors responsible for lineage-specific behaviors.

## FUNDING

The research leading to these results has received funding from the Italian Ministry of Health “Ricerca Finalizzata 2016” under grant agreement GR-2016-02364014 to PM. The funders had no role in the design of this study and will not have any role during its execution, analyses, interpretation of the data, or decision to submit results.

## CONFLICT OF INTERESTS

None to declare.

## AUTHOR CONTRIBUTIONS

Matteo Chiacchiaretta: Methodology, Formal analysis, Investigation, Visualization, Writing – Original draft, Writing – Review & Editing

Rita Sorrentino: Methodology, Formal analysis, Investigation, Visualization, Writing – Original draft, Writing – Review & Editing

Nadia Bresciani: Investigation, Writing – Review & Editing

Alessandra Aiello: Investigation, Writing – Review & Editing

Valentina Vanini: Investigation, Writing – Review & Editing

Samuel Zambrano: Methodology, Formal analysis, Data curation, Writing – Review & Editing

Davide Mazza: Formal analysis, Data curation, Writing – Review & Editing

Federica Cugnata: Formal analysis, Data curation, Visualization, Writing – Review & Editing

Daniela M. Cirillo: Methodology, Writing – Review & Editing

Alessandra Agresti: Methodology, Visualization, Writing – Review & Editing

Delia Goletti: Methodology, Writing – Review & Editing

Paolo Miotto: Conceptualization, Methodology, Formal analysis, Investigation, Visualization, Supervision, Project administration, Writing – Original draft, Writing – Review & Editing, Funding acquisition

## DATA AVAILABILITY STATEMENT

Data is provided within the manuscript or supplementary information files.

## FIGURE LEGENDS

**Figure S1**: Expression profiling of specific markers for M1 (IL-1β, IL-6, CD80, TNF-α) and M2 (CD206) after polarization stimuli. Gene expression is provided as fold-change compared to M0 THP-1-derived macrophages (endogenous control: GAPDH).

**Figure S2:** Top panel. Temporal dynamics of phagosomal acidification in M1 macrophages infected with MTB strains. (Left) Bubble plot summarizing the most relevant variables at 6h and 24h post-infection (p.i.) across the most diverse strain categories (GEN, SPEC). Variables include the number of LysoSensor-positive compartments, acidification levels, colocalization of MTB with acidified compartments, percentage of responding cells, and bacillary load (M/C, mycobacteria per cell). Bubble size reflects relative magnitude. (Right) Schematic illustration of infected macrophages showing LysoSensor-positive compartments at 6h and 24h p.i., corresponding to the trends depicted in the bubble plot. The drawings highlight differences in the number and intensity of acidified compartments, bacterial colocalization with acidified vesicles, and overall bacillary burden between GEN and SPEC strains over time. This figure provides a representative visual summary of the key trends, rather than an exhaustive display of all measured variables. GEN: Generalist strains *M. tuberculosis* 12594/02 (L2), *M. tuberculosis* 1934/03 (L2) *M. tuberculosis* H37Rv NCTC 7416 (L4); SPEC: Specialist strains *M. africanum* 10514/01 (L6), *M. africanum* 5468/02 (L6), *M. tuberculosis* 1797/03 (L1), *M. tuberculosis* 4850/03 (L1). Bottom panel. Temporal dynamics of phagosomal acidification in M2 macrophages infected with MTB strains. (Left) Bubble plot summarizing the most relevant variables at 6h and 24h post-infection (p.i.) across the most diverse strain categories (GEN, SPEC). Variables include the number of LysoSensor-positive compartments, acidification levels, colocalization of MTB with acidified compartments, percentage of responding cells, and bacillary load (M/C, mycobacteria per cell). Bubble size reflects relative magnitude. (Right) Schematic illustration of infected macrophages showing LysoSensor-positive compartments at 6h and 24h p.i., corresponding to the trends depicted in the bubble plot. The drawings highlight differences in the number and intensity of acidified compartments, bacterial colocalization with acidified vesicles, and overall bacillary burden between GEN and SPEC strains over time. This figure provides a representative visual summary of the key trends, rather than an exhaustive display of all measured variables. GEN: Generalist strains *M. tuberculosis* 12594/02 (L2), *M. tuberculosis* 1934/03 (L2) *M. tuberculosis* H37Rv NCTC 7416 (L4); SPEC: Specialist strains *M. africanum* 10514/01 (L6), *M. africanum* 5468/02 (L6), *M. tuberculosis* 1797/03 (L1), *M. tuberculosis* 4850/03 (L1).

Created in BioRender.

## Notes

### Competing Interest Statement

The authors have declared no competing interest.

## REFERENCES

1. Huang L, Nazarova EV, Russell DG. 2019. Bacterial Fitness within the Host Macrophage. Microbiol Spectr 7.

2. Barbier M, Wirth T. 2016. The Evolutionary History, Demography, and Spread of the *Mycobacterium tuberculosis* Complex. Microbiol Spectr 4.

3. Romagnoli A, Petruccioli E, Palucci I, Camassa S, Carata E, Petrone L, Mariano S, Sali M, Dini L, Girardi E, Delogu G, Goletti D, Fimia GM. 2018. Clinical isolates of the modern *Mycobacterium tuberculosis* lineage 4 evade host defense in human macrophages through eluding IL-1β-induced autophagy. Cell Death Dis 9:624.

4. Gagneux S. 2018. Ecology and evolution of *Mycobacterium tuberculosis*. Nat Rev Microbiol 16:202–213.

5. Bottai D, Frigui W, Sayes F, Di Luca M, Spadoni D, Pawlik A, Zoppo M, Orgeur M, Khanna V, Hardy D, Mangenot S, Barbe V, Medigue C, Ma L, Bouchier C, Tavanti A, Larrouy-Maumus G, Brosch R. 2020. TbD1 deletion as a driver of the evolutionary success of modern epidemic *Mycobacterium tuberculosis* lineages. Nat Commun 11:684.

6. Casadevall A. 2017. The Pathogenic Potential of a Microbe. mSphere 2.

7. Möller M, Kinnear CJ, Orlova M, Kroon EE, van Helden PD, Schurr E, Hoal EG. 2018. Genetic Resistance to *Mycobacterium tuberculosis* Infection and Disease. Front Immunol 9:2219.

8. Rosain J, Kong XF, Martinez-Barricarte R, Oleaga-Quintas C, Ramirez-Alejo N, Markle J, Okada S, Boisson-Dupuis S, Casanova JL, Bustamante J. 2019. Mendelian susceptibility to mycobacterial disease: 2014-2018 update. Immunol Cell Biol 97:360–367.

9. Khan A, Singh VK, Hunter RL, Jagannath C. 2019. Macrophage heterogeneity and plasticity in tuberculosis. J Leukoc Biol 106:275–282.

10. Capuano SV, Croix DA, Pawar S, Zinovik A, Myers A, Lin PL, Bissel S, Fuhrman C, Klein E, Flynn JL. 2003. Experimental Mycobacterium tuberculosis infection of cynomolgus macaques closely resembles the various manifestations of human M. tuberculosis infection. Infect Immun 71:5831–44.

11. Dinkele R, Gessner S, McKerry A, Leonard B, Seldon R, Koch AS, Morrow C, Gqada M, Kamariza M, Bertozzi CR, Smith B, McLoud C, Kamholz A, Bryden W, Call C, Kaplan G, Mizrahi V, Wood R, Warner DF. 2021. Capture and visualization of live Mycobacterium tuberculosis bacilli from tuberculosis patient bioaerosols. PLoS Pathog 17:e1009262.

12. Homolka S, Niemann S, Russell DG, Rohde KH. 2010. Functional genetic diversity among *Mycobacterium tuberculosis* complex clinical isolates: delineation of conserved core and lineage-specific transcriptomes during intracellular survival. PLoS Pathog 6:e1000988.

13. Stucki D, Brites D, Jeljeli L, Coscolla M, Liu Q, Trauner A, Fenner L, Rutaihwa L, Borrell S, Luo T, Gao Q, Kato-Maeda M, Ballif M, Egger M, Macedo R, Mardassi H, Moreno M, Tudo Vilanova G, Fyfe J, Globan M, Thomas J, Jamieson F, Guthrie JL, Asante-Poku A, Yeboah-Manu D, Wampande E, Ssengooba W, Joloba M, Henry Boom W, Basu I, Bower J, Saraiva M, Vaconcellos SEG, Suffys P, Koch A, Wilkinson R, Gail-Bekker L, Malla B, Ley SD, Beck HP, de Jong BC, Toit K, Sanchez-Padilla E, Bonnet M, Gil-Brusola A, Frank M, Penlap Beng VN, Eisenach K, Alani I, Wangui Ndung’u P, et al. 2016. *Mycobacterium tuberculosis* lineage 4 comprises globally distributed and geographically restricted sublineages. Nat Genet 48:1535–1543.

14. Genin M, Clement F, Fattaccioli A, Raes M, Michiels C. 2015. M1 and M2 macrophages derived from THP-1 cells differentially modulate the response of cancer cells to etoposide. BMC Cancer 15:577.

15. Mantovani A, Sica A, Sozzani S, Allavena P, Vecchi A, Locati M. 2004. The chemokine system in diverse forms of macrophage activation and polarization. Trends Immunol 25:677–86.

16. Murray PJ, Allen JE, Biswas SK, Fisher EA, Gilroy DW, Goerdt S, Gordon S, Hamilton JA, Ivashkiv LB, Lawrence T, Locati M, Mantovani A, Martinez FO, Mege JL, Mosser DM, Natoli G, Saeij JP, Schultze JL, Shirey KA, Sica A, Suttles J, Udalova I, van Ginderachter JA, Vogel SN, Wynn TA. 2014. Macrophage activation and polarization: nomenclature and experimental guidelines. Immunity 41:14–20.

17. Schindelin J, Arganda-Carreras I, Frise E, Kaynig V, Longair M, Pietzsch T, Preibisch S, Rueden C, Saalfeld S, Schmid B, Tinevez JY, White DJ, Hartenstein V, Eliceiri K, Tomancak P, Cardona A. 2012. Fiji: an open-source platform for biological-image analysis. Nat Methods 9:676–82.

18. Pinheiro JC, Bates DM. 2000. Mixed-Effects Models in S and S-PLUS, 1 ed. Springer New York, NY.

19. Laird NM, Ware JH. 1982. Random-effects models for longitudinal data. Biometrics 38:963–74.

20. Geraci M. 2014. Linear Quantile Mixed Models: The lqmm Package for Laplace Quantile Regression. Journal of Statistical Software 57:1–29.

21. Brooks ME, Kristensen K, van Benthem KJ, Magnusson A, Berg CW, Nielsen A, Skaug HJ, Maechler M, Bolker BM. 2017. glmmTMB Balances Speed and Flexibility Among Packages for Zero-inflated Generalized Linear Mixed Modeling. The R Journal 9:378–400.

22. Team RC. 2013. R: A language and environment for statistical computing. R Foundation for Statistical Computing Vienna, Austria.

23. Feoktistova M, Geserick P, Leverkus M. 2016. Crystal Violet Assay for Determining Viability of Cultured Cells. Cold Spring Harb Protoc 2016:pdb.prot087379.

24. Palucci I, Maulucci G, De Maio F, Sali M, Romagnoli A, Petrone L, Fimia GM, Sanguinetti M, Goletti D, De Spirito M, Piacentini M, Delogu G. 2019. Inhibition of Transglutaminase 2 as a Potential Host-Directed Therapy Against. Front Immunol 10:3042.

25. Genestet C, Bernard-Barret F, Hodille E, Ginevra C, Ader F, Goutelle S, Lina G, Dumitrescu O, group LTs. 2018. Antituberculous drugs modulate bacterial phagolysosome avoidance and autophagy in Mycobacterium tuberculosis-infected macrophages. Tuberculosis (Edinb) 111:67–70.

26. Buter J, Cheng TY, Ghanem M, Grootemaat AE, Raman S, Feng X, Plantijn AR, Ennis T, Wang J, Cotton RN, Layre E, Ramnarine AK, Mayfield JA, Young DC, Jezek Martinot A, Siddiqi N, Wakabayashi S, Botella H, Calderon R, Murray M, Ehrt S, Snider BB, Reed MB, Oldfield E, Tan S, Rubin EJ, Behr MA, van der Wel NN, Minnaard AJ, Moody DB. 2019. *Mycobacterium tuberculosis* releases an antacid that remodels phagosomes. Nat Chem Biol 15:889–899.

27. Zheng W, Chang IC, Limberis J, Budzik JM, Zha BS, Howard Z, Chen L, Ernst JD. 2024. Mycobacterium tuberculosis resides in lysosome-poor monocyte-derived lung cells during chronic infection. PLoS Pathog 20:e1012205.

28. Bedard M, van der Niet S, Bernard EM, Babunovic G, Cheng TY, Aylan B, Grootemaat AE, Raman S, Botella L, Ishikawa E, O’Sullivan MP, O’Leary S, Mayfield JA, Buter J, Minnaard AJ, Fortune SM, Murphy LO, Ory DS, Keane J, Yamasaki S, Gutierrez MG, van der Wel N, Moody DB. 2023. A terpene nucleoside from M. tuberculosis induces lysosomal lipid storage in foamy macrophages. J Clin Invest 133.

29. Domenech P, Zou J, Averback A, Syed N, Curtis D, Donato S, Reed MB. 2017. Unique Regulation of the DosR Regulon in the Beijing Lineage of *Mycobacterium tuberculosis*. J Bacteriol 199.

30. Banaei-Esfahani A, Trauner A, Borrell S, Gygli SM, Rustad TR, Feldmann J, Gillet LC, Schubert OT, Sherman DR, Beisel C, Gagneux S, Aebersold R, Collins BC. 2020. Network analysis identifies regulators of lineage-specific phenotypes in *Mycobacterium tuberculosis*. bioRxiv:2020.02.14.943365.

31. Reichlen MJ, Leistikow RL, Scobey MS, Born SEM, Voskuil MI. 2017. Anaerobic *Mycobacterium tuberculosis* Cell Death Stems from Intracellular Acidification Mitigated by the DosR Regulon. J Bacteriol 199.

32. Butler RE, Cihlarova V, Stewart GR. 2010. Effective generation of reactive oxygen species in the mycobacterial phagosome requires K+ efflux from the bacterium. Cell Microbiol 12:1186–93.

33. Malone KM, Gordon SV. 2017. *Mycobacterium tuberculosi*s Complex Members Adapted to Wild and Domestic Animals. In: Gagneux, S. (eds) Strain Variation in the Mycobacterium tuberculosis Complex: Its Role in Biology, Epidemiology and Control, vol 1019. Springer International Publishing, Cham.

34. Panchal V, Jatana N, Malik A, Taneja B, Pal R, Bhatt A, Besra GS, Thukral L, Chaudhary S, Rao V. 2019. A novel mutation alters the stability of PapA2 resulting in the complete abrogation of sulfolipids in clinical mycobacterial strains. FASEB Bioadv 1:306–319.

35. Canton J. 2014. Phagosome maturation in polarized macrophages. J Leukoc Biol 96:729–38.

36. Wojtas B, Fijalkowska B, Wlodarczyk A, Schollenberger A, Niemialtowski M, Hamasur B, Pawlowski A, Krzyzowska M. 2011. Mannosylated lipoarabinomannan balances apoptosis and inflammatory state in mycobacteria-infected and uninfected bystander macrophages. Microb Pathog 51:9–21.

37. Gupta S, Rodriguez GM. 2018. Mycobacterial extracellular vesicles and host pathogen interactions. Pathog Dis 76.

38. Singh PP, LeMaire C, Tan JC, Zeng E, Schorey JS. 2011. Exosomes released from *M. tuberculosis* infected cells can suppress IFN-γ mediated activation of naïve macrophages. PLoS One 6:e18564.

39. Wiens KE, Woyczynski LP, Ledesma JR, Ross JM, Zenteno-Cuevas R, Goodridge A, Ullah I, Mathema B, Djoba Siawaya JF, Biehl MH, Ray SE, Bhattacharjee NV, Henry NJ, Reiner RC, Kyu HH, Murray CJL, Hay SI. 2018. Global variation in bacterial strains that cause tuberculosis disease: a systematic review and meta-analysis. BMC Med 16:196.

40. Reiling N, Homolka S, Kohl TA, Steinhäuser C, Kolbe K, Schütze S, Brandenburg J. 2018. Shaping the niche in macrophages: Genetic diversity of the *M. tuberculosis* complex and its consequences for the infected host. Int J Med Microbiol 308:118–128.

41. Fejer G, Wegner MD, Györy I, Cohen I, Engelhard P, Voronov E, Manke T, Ruzsics Z, Dölken L, Prazeres da Costa O, Branzk N, Huber M, Prasse A, Schneider R, Apte RN, Galanos C, Freudenberg MA. 2013. Nontransformed, GM-CSF-dependent macrophage lines are a unique model to study tissue macrophage functions. Proc Natl Acad Sci U S A 110:E2191–8.

42. Pahari S, Arnett E, Simper J, Azad A, Guerrero-Arguero I, Ye C, Zhang H, Cai H, Wang Y, Lai Z, Jarvis N, Lumbreras M, Maselli DJ, Peters J, Torrelles JB, Martinez-Sobrido L, Schlesinger LS. 2023. A new tractable method for generating human alveolar macrophage-like cells. mBio:e0083423.

43. Madhvi A, Mishra H, Leisching GR, Mahlobo PZ, Baker B. 2019. Comparison of human monocyte derived macrophages and THP1-like macrophages as in vitro models for *M. tuberculosis* infection. Comp Immunol Microbiol Infect Dis 67:101355.

44. Kurynina AV, Erokhina MV, Makarevich OA, Sysoeva VY, Lepekha LN, Kuznetsov SA, Onishchenko GE. 2018. Plasticity of Human THP-1 Cell Phagocytic Activity during Macrophagic Differentiation. Biochemistry (Mosc) 83:200–214.

45. Abuawad A, Mbadugha C, Ghaemmaghami AM, Kim DH. 2020. Metabolic characterisation of THP-1 macrophage polarisation using LC-MS-based metabolite profiling. Metabolomics 16:33.

46. Li P, Hao Z, Wu J, Ma C, Xu Y, Li J, Lan R, Zhu B, Ren P, Fan D, Sun S. 2021. Comparative Proteomic Analysis of Polarized Human THP-1 and Mouse RAW264.7 Macrophages. Front Immunol 12:700009.

47. Shah PT, Tufail M, Wu C, Xing L. 2022. THP-1 cell line model for tuberculosis: A platform for in vitro macrophage manipulation. Tuberculosis (Edinb) 136:102243.

48. Lim JJ, Grinstein S, Roth Z. 2017. Diversity and Versatility of Phagocytosis: Roles in Innate Immunity, Tissue Remodeling, and Homeostasis. Front Cell Infect Microbiol 7:191.

49. Rothchild AC, Olson GS, Nemeth J, Amon LM, Mai D, Gold ES, Diercks AH, Aderem A. 2019. Alveolar macrophages generate a noncanonical NRF2-driven transcriptional response. Sci Immunol 4.

50. Pisu D, Huang L, Narang V, Theriault M, Lê-Bury G, Lee B, Lakudzala AE, Mzinza DT, Mhango DV, Mitini-Nkhoma SC, Jambo KC, Singhal A, Mwandumba HC, Russell DG. 2021. Single cell analysis of *M. tuberculosis* phenotype and macrophage lineages in the infected lung. J Exp Med 218.

51. Huang L, Nazarova EV, Tan S, Liu Y, Russell DG. 2018. Growth of *Mycobacterium tuberculosis in vivo* segregates with host macrophage metabolism and ontogeny. J Exp Med 215:1135–1152.

52. Zheng W, Borja M, Dorman LC, Liu J, Zhou A, Seng A, Arjyal R, Sunshine S, Nalyvayko A, Pisco AO, Rosenberg OS, Neff N, Zha BS. 2025. Single-cell analysis reveals. Sci Adv 11:eadq8158.

53. Solomon SL, Bryson BD. 2023. Single-cell analysis reveals a weak macrophage subpopulation response to Mycobacterium tuberculosis infection. Cell Rep 42:113418.

54. Bussi C, Gutierrez MG. 2019. *Mycobacterium tuberculosis* infection of host cells in space and time. FEMS Microbiol Rev 43:341–361.

55. Kilinç G, Saris A, Ottenhoff THM, Haks MC. 2021. Host-directed therapy to combat mycobacterial infections. Immunol Rev 301:62–83.

56. Taya T, Teruyama F, Gojo S. 2023. Host-directed therapy for bacterial infections -Modulation of the phagolysosome pathway. Front Immunol 14:1227467.

