## Supplementary figures and images for "*Mycobacterium tuberculosis* diversity and macrophage heterogeneity dictate phagosomal acidification"

### Figure S1

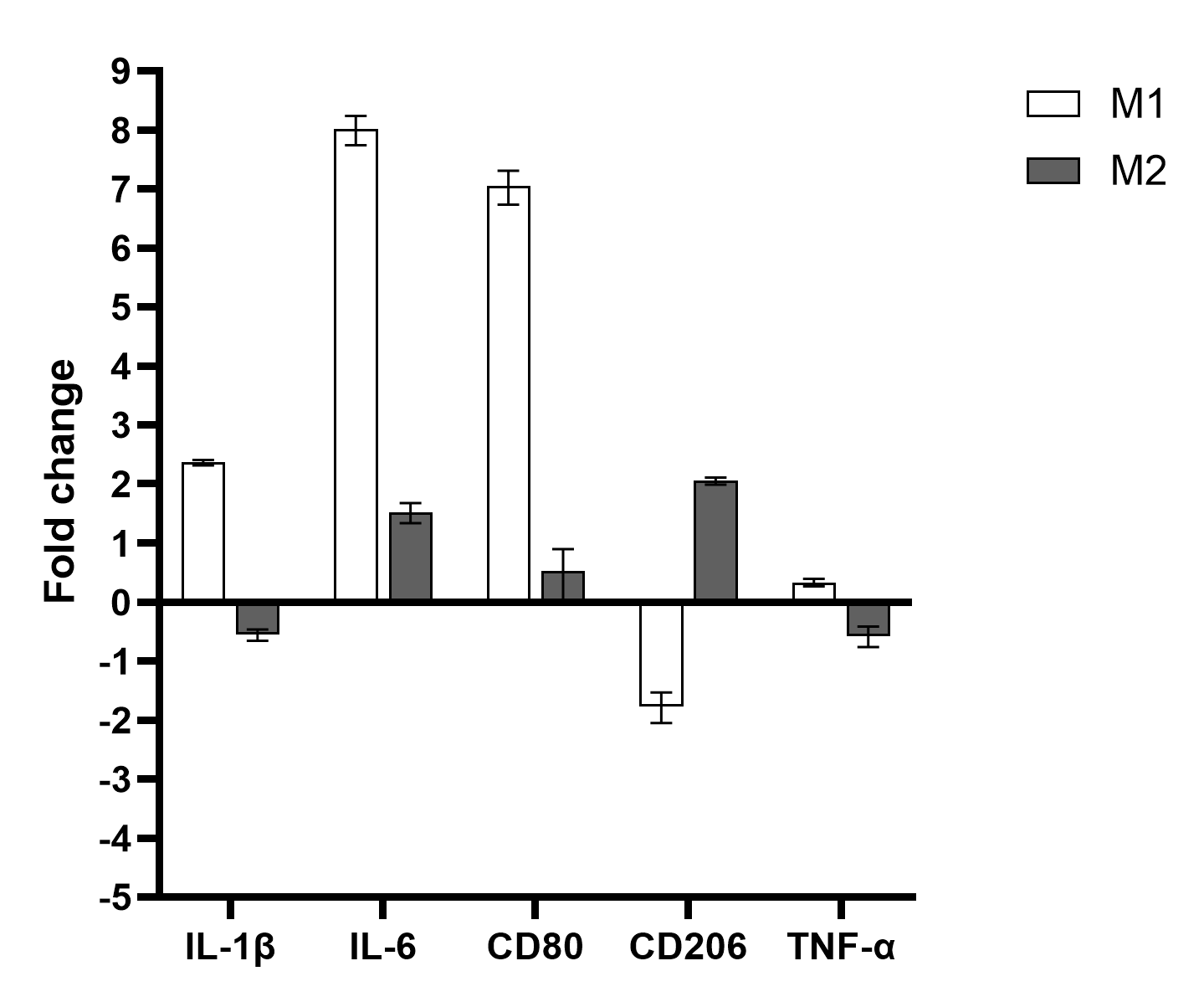

### Figure S2

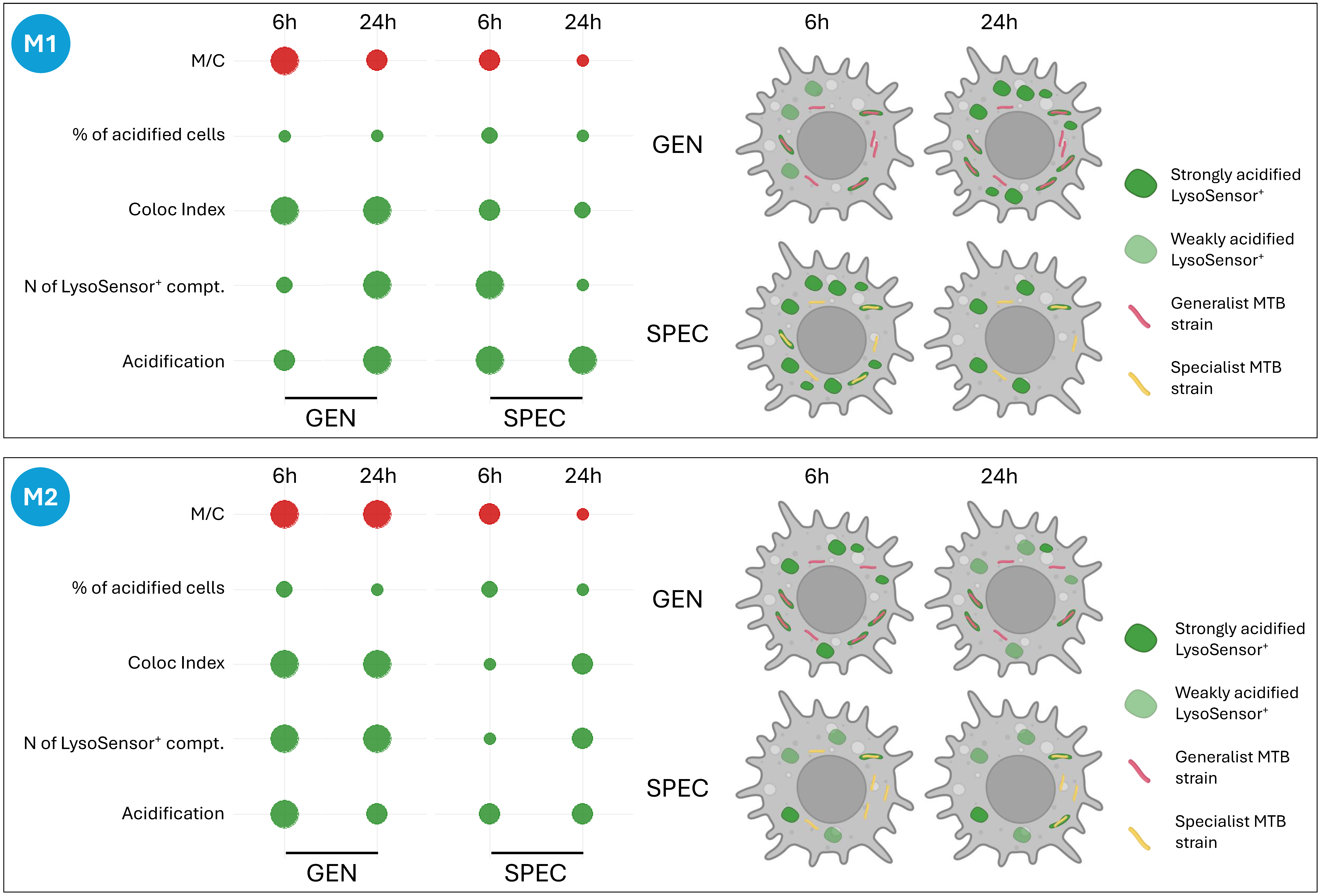
