## Supplementary 1 for "*Mycobacterium tuberculosis* diversity and macrophage heterogeneity dictate phagosomal acidification"

**MATERIALS AND METHODS**

***M. tuberculosis* strains engineering**

All mycobacterial strains were engineered to express mCherry (Ex/Em 587-610 nm) from a low copy number plasmid. Briefly, bacteria were cultured in liquid 7H9 medium + Albumin Dextrose Catalase Oleic Acid (OADC) supplement (Becton Dickinson, New Jersey, USA), 0.05% Tween® 80 (Sigma Aldrich, St. Luis, MO, USA) until $OD_{600}$ 0.4-0.6. Bacteria were centrifuged and washed twice with 10% glycerol, to eliminate salt and contaminants, to achieve the electro-competence, and resuspended in 10% glycerol for frozen stockage. Competent cells were electroporated (2500V, 25μF, 200 ohms) with 2μg of salt-free plasmid and selected on 50μg/mL Hygromycin (Roche, Basel, Switzerland) 7H10 agar plates (Becton Dickinson, New Jersey, USA) [1].

For macrophage infection, the strains were thawed on selective Middlebrook 7H10 agar medium supplemented with 10% OADC supplement, 0.05% Tween® 80, and 50μg/mL hygromycin. A single colony was picked up and used for inoculating Middlebrook 7H9 liquid medium supplemented with 10% Middlebrook OADC supplement, 0.05% Tween® 80, and 50μg/mL hygromycin and incubated at 37 °C until $OD_{600}$~ 0.3. Bacterial suspensions were centrifuged at 17000 g for 5 minutes, washed with sterile PBS, and resuspended in complete RPMI (cRPMI) medium w/o antibiotics, de-clumped by 10 passages through a 25-gauge needle, and diluted in cRPMI immediately before the infection.

**Cells preparation and infection for confocal microscopy**

THP-1 were used for infection experiments at passages 7 to 14 to ensure consistent differentiation and functional responses. Briefly, THP-1 cells (ATCC® TIB-202™) were differentiated in cRPMI (with L-glutamine 2mM, Hepes 10mM, Non-essential amino acids 100nM, Sodium Pyruvate 1mM, Sigma Aldrich, Missouri, USA) supplemented with 10% FBS (Fetal Bovin Serum, Euroclone, Italy), antibiotics (penicillin G 100 U/mL and streptomycin sulfate 100 U/mL, Sigma Aldrich, Missouri, USA), and 100 nM of phorbol 12-myristate 13-acetate (PMA, Sigma Aldrich, Missouri, USA) for 3 days in Eppendorf Cell Imaging 96-wells plates (55,000 cells/well) bottom glass 170μm thick (Eppendorf, Germany). Type I collagen

(Sigma Aldrich, Missouri, USA) was used for improving cell adhesion on bottom glass surface, mimic more physiological conditions and improve both morphology and experimental consistency.

Infection was carried out by centrifugation (300 g x 5 min, room temperature) followed by two hours of incubation at 37 °C and 5% $CO_2$. Macrophage-like cells were then washed twice and supplemented with cRPMI without antibiotics. cRPMI without phenol red was used in case the infection was set up for imaging readouts.

**Images and data analysis**

At least 5 infected (e.g. internalized mycobacteria) plus 5 uninfected cells (e.g. exposed to mycobacteria in the same well of infected cells, but no uptake observed) (unless not available) were considered for each acquisition. Uninfected controls (CTRL) are considered macrophages not exposed to mycobacteria (in a different well).

<u>Phagosomal acidification</u>: To assess acidification dynamics in infected macrophages, we employed LysoSensor staining combined with confocal microscopy and z-stack acquisition. The LysoSensor dye is an acidotropic probe exhibiting a dynamic pH-dependent increase in fluorescence intensity upon acidification only, with very low background in the cytosol. The use of LysoSensor is therefore more appropriate compared to LysoTracker dyes, as it allows to monitor pH changes rather than track acidic organelles. LysoSensor Green was used to quantify the abundance and intensity of acidified compartments in THP-1-derived M1 and M2 macrophages. To specifically assess phagosomal acidification, LysoSensor fluorescence was analyzed in conjunction with intracellular bacterial localization. Acidified compartments and mycobacteria were selected by thresholding the 458 nm and the 561 nm channels, respectively. Intracellular acidification was quantified by applying a size-filtered region of interest (ROI) mask to identify LysoSensor-positive structures, excluding background signal and capturing a defined range of acidic compartments — including but not limited to lysosomes and phagosomes. Briefly, LysoSensor-positive compartments were segmented using Fiji by applying an intensity threshold (Huang method) and size filtering (0.5–1.8 $\mu m^2$) to focus on medium- to large-sized acidified vesicles. This approach excludes background and smaller lysosomes (<0.5 $\mu m^2$) while capturing structures consistent with mature lysosomes, phagolysosomes, and other larger acidic compartments. For clarity, these

LysoSensor-positive structures are hereafter referred to as acidified compartments, encompassing large lysosomes as well as phagolysosomes. Acidified compartments and mycobacteria counts were also acquired for each single cell ROI.

To specifically assess phagosomal acidification, we performed colocalization analysis between LysoSensor fluorescence and internalized bacteria, allowing us to distinguish pathogen-associated acidified compartments from the overall lysosomal response. To analyze the spatial relationship between bacteria and lysosomes (colocalization), we wrote a custom pipeline in Matlab [2]. Briefly, the confocal stacks are maximal projected and the individual cells are segmented manually. Next, the cell nuclei and the bacteria are automatically segmented, and the average LysoSensor intensity around the segmented bacteria $L_{bac}$ is calculated (in a region of 1 pixel in width). Such intensity is compared to the average LysoSensor intensity in the cell $\langle L \rangle$, to provide an inclusion index $I_i=( L_{bac}-\langle L \rangle)/\langle L \rangle$ (Figure 1a). Such index would be higher than zero when bacteria are located inside the LysoSensor-positive acidified compartments and lower than zero when bacteria are excluded from the acidified compartments. Random positioning of bacteria would provide $I_i=0$. Cells touching the edge of the image have been excluded from the analysis.

Statistical analysis: These procedures were employed to properly account for the dependence structure among observations, by including nested random-effect terms, thus considering in the model unobservable sources of heterogeneity among experimental units. For numeric variables, uninfected control THP-1 macrophages (M1 or M2, no exposure to mycobacteria) were used for normalization. Evaluation of bystander effect, defined as an increase of acidification in adjacent, uninfected macrophages (exposed to mycobacteria, but no uptake of mycobacteria observed) we considered the percentage of acidified cells. A cell is considered acidified if its phagosomal acidification value exceeds the upper control limit (mean+1.96SD) based on the THP-1 macrophage controls. In the linear mixed-effects models, to meet the assumption of normality of the residuals of the models, an appropriate transformation was eventually applied to the response variable; and, when necessary, few outliers identified by the specific model were not included in that analysis. Linear quantile mixed models on the median were applied to test differences between M1 and M2. Zero-inflated Mixed-effects Poisson models were applied to properly analyze count data with

excess zeros. For testing difference between pairs of groups, post-hoc analysis was performed applying the FDR procedure to account for multiple comparisons.
Spearman's correlation coefficient was calculated to evaluate the presence of a monotonic relationship between variables.

**Macrophage survival**

Briefly, cells were fixed with cold (-20 °C) methanol (Sigma Aldrich, Missouri, USA) and stained with Crystal violet. This experiment relies on the detachment of adherent cells from cell culture plates during cell death where only live, adherent cells are fixed and stained with Crystal violet. After a wash step, the Crystal violet dye is solubilized with lysis buffer (0.1M sodium citrate, 50% Ethanol, pH 4.2) and measured by absorbance at 570 nm using the BioTek™ Gen5™ software (Agilent).

**References**

1. Maciag A, Dainese E, Rodriguez GM, et al. Global analysis of the *Mycobacterium tuberculosis* Zur (FurB) regulon. J Bacteriol **2007**; 189:730-40.
2. The MathWorks I. MATLAB  version: 9.4.0.813654 (R2018a).  **2018**.
