## Supplementary 2 for "*Mycobacterium tuberculosis* diversity and macrophage heterogeneity dictate phagosomal acidification"

### *PHAGOSOME ACIDIFICATION*

Number of M1 and M2 macrophages considered for each MTBC strain, at each timepoint:

### *6 h p.i.*

| *6 h p.i.* | | | | | | | |
|---|---|---|---|---|---|---|---|
| | | NOT infected | | Infected | | *TOT* | |
| | | N of cells | Replicates | N of cells | Replicates | Not inf | Inf |
| M1 | CTRL | 197 | 8 | *na* | *na* | *1141* | *909* |
| | H37Rv | 130 | 3 | 116 | 3 | | |
| | H37Ra | 119 | 3 | 105 | 3 | | |
| | BCG | 78 | 2 | 84 | 2 | | |
| | CDC1551 | 129 | 3 | 122 | 3 | | |
| | Beijing A | 122 | 3 | 129 | 3 | | |
| | Beijing B | 60 | 2 | 70 | 2 | | |
| | EAI A | 67 | 2 | 66 | 2 | | |
| | EAI B | 79 | 2 | 71 | 2 | | |
| | Afri A | 93 | 2 | 80 | 2 | | |
| | Afri B | 67 | 2 | 66 | 2 | | |
| M2 | CTRL | 190 | 8 | *na* | *na* | *1116* | *932* |
| | H37Rv | 111 | 3 | 113 | 3 | | |
| | BCG | 73 | 2 | 84 | 2 | | |
| | H37Ra | 131 | 3 | 121 | 3 | | |
| | CDC1551 | 134 | 3 | 123 | 3 | | |
| | Beijing A | 126 | 3 | 148 | 3 | | |
| | Beijing B | 60 | 2 | 70 | 2 | | |
| | EAI A | 61 | 2 | 61 | 2 | | |
| | EAI B | 81 | 2 | 73 | 2 | | |
| | Afri A | 83 | 2 | 77 | 2 | | |
| | Afri B | 66 | 2 | 62 | 2 | | |
| | *TOT* | *2257* | | *1841* | | | |
| | | | **4098** | | | | |

### *24 h p.i.*

| *24 h p.i.* | | | | | | | |
|---|---|---|---|---|---|---|---|
| | | NOT infected | | Infected | | *TOT* | |
| | | N of cells | Replicates | N of cells | Replicates | Not inf | Inf |
| M1 | CTRL | 152 | 8 | *na* | *na* | *813* | *806* |
| | H37Rv | 77 | 3 | 110 | 3 | | |
| | H37Ra | 73 | 3 | 111 | 3 | | |
| | BCG | 51 | 2 | 76 | 2 | | |
| | CDC1551 | 75 | 3 | 91 | 3 | | |
| | Beijing A | 81 | 3 | 102 | 3 | | |
| | Beijing B | 48 | 2 | 67 | 2 | | |
| | EAI A | 60 | 2 | 60 | 2 | | |
| | EAI B | 76 | 2 | 67 | 2 | | |
| | Afri A | 53 | 2 | 59 | 2 | | |
| | Afri B | 67 | 2 | 63 | 2 | | |
| M2 | CTRL | 138 | 8 | *na* | *na* | *859* | *808* |
| | H37Rv | 87 | 3 | 105 | 3 | | |
| | BCG | 70 | 2 | 80 | 2 | | |
| | H37Ra | 94 | 3 | 109 | 3 | | |
| | CDC1551 | 84 | 3 | 82 | 3 | | |
| | Beijing A | 83 | 3 | 98 | 3 | | |
| | Beijing B | 43 | 2 | 64 | 2 | | |
| | EAI A | 61 | 2 | 61 | 2 | | |
| | EAI B | 80 | 2 | 80 | 2 | | |
| | Afri A | 60 | 2 | 64 | 2 | | |
| | Afri B | 59 | 2 | 65 | 2 | | |
| | *TOT* | *1672* | | *1614* | | | |
| | | | **3286** | | | | |

### *Total*

| | | *Overall* | | | | | |
|---|---|---|---|---|---|---|---|
| | | NOT infected | | Infected | | *TOT* | |
| | | N of cells | Replicates | N of cells | Replicates | Not inf | Inf |
| M1 | CTRL | 349 | 8 | *na* | *na* | *1954* | *1715* |
| | H37Rv | 207 | 3 | 226 | 3 | | |
| | H37Ra | 192 | 3 | 216 | 3 | | |
| | BCG | 129 | 2 | 160 | 2 | | |
| | CDC1551 | 204 | 3 | 213 | 3 | | |
| | Beijing A | 203 | 3 | 231 | 3 | | |
| | Beijing B | 108 | 2 | 137 | 2 | | |
| | EAI A | 127 | 2 | 126 | 2 | | |
| | EAI B | 155 | 2 | 138 | 2 | | |
| | Afri A | 146 | 2 | 139 | 2 | | |
| | Afri B | 134 | 2 | 129 | 2 | | |
| M2 | CTRL | 328 | 8 | *na* | *na* | *1975* | *1740* |
| | H37Rv | 198 | 3 | 218 | 3 | | |
| | BCG | 143 | 2 | 164 | 2 | | |
| | H37Ra | 225 | 3 | 230 | 3 | | |
| | CDC1551 | 218 | 3 | 205 | 3 | | |
| | Beijing A | 209 | 3 | 246 | 3 | | |
| | Beijing B | 103 | 2 | 134 | 2 | | |
| | EAI A | 122 | 2 | 122 | 2 | | |
| | EAI B | 161 | 2 | 153 | 2 | | |
| | Afri A | 143 | 2 | 141 | 2 | | |
| | Afri B | 125 | 2 | 127 | 2 | | |
| | *TOT* | *3929* | | *3455* | | | |
| | | | 7384 | | | | |

### *Imaging acquisition and analysis details*

| | |
|---|---|
| *Acquisition:* | obj. **63x, oil NA 1.40**<br>**z-stacks: 0.89 µm x 7 stacks**<br>Res. **16-bit**, Format: **1024x1024**<br>Laser 405 (for Hoechst) = **6%** gain: 898.9; offset: 0%<br>Laser 458 (for LysoSensor) = **10% (Argon power: 30%); gain:724.8**, offset: 0%<br>Laser 561 (for mCherry) = **8%; gain: 800**, offset 0%<br>well/sample: 3<br>acquistion postions/well: 2 |
| *Analysis:* | 6 h p.i. and 24 h p.i.; (N of phagolysosomes), phagolysosomes intensity and % of active cells, MOI, colocaluzation myc/acidic compartments |
| **Hoechst (Ex/Em 350/461)** | threshold (Huang, Red; no flags on Dark background and Stack histogram) = approx. 215<br>size = 40-inf $\mu m^2$, circularity = 0-1 (show: overlay mask; flag only Add to Manager) |
| **LysoSensor (Ex/Em 443/505)** | **threshold (Huang, Red; no flags on Dark background and Stack histogram) = 155**<br>**size = 0.5-1.8 $\mu m^2$, circularity = 0-1 (show: overlay mask; NO flag on Add to Manager)** |
| **mCherry (Ex/Em 587-610)** | **threshold (Huang, Red; no flags on Dark background and Stack histogram) = 215**<br>**size = 2-100 $\mu m^2$, circularity = 0-0.9 (show: overlay mask; NO flag on Add to Manager)** |

# M1 – 6 h p.i.

### a) Acifdification of LysoSensor⁺ compartments

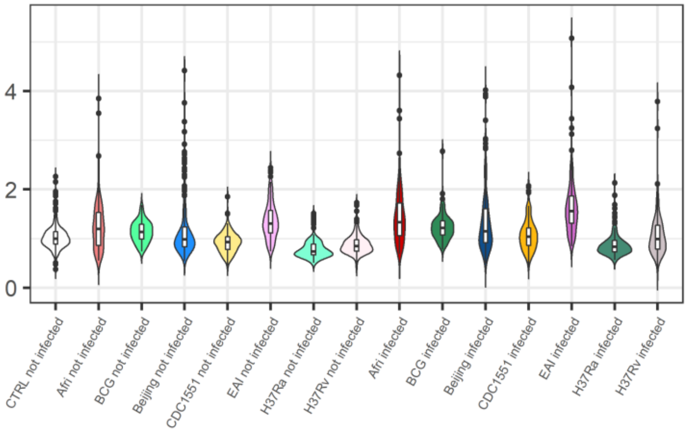


| Par | Value | Std.Error | p.value |
|---|---|---|---|
| (Intercept) | 0.0058 | 0.0668 | 0.9308 |
| groupAfri not infected | 0.1237 | 0.0695 | 0.0754 |
| groupBCG not infected | 0.0317 | 0.1052 | 0.7631 |
| groupBeijing not infected | 0.0952 | 0.0641 | 0.1379 |
| groupCDC1551 not infected | -0.0414 | 0.0774 | 0.5923 |
| groupEAI not infected | 0.1613 | 0.075 | 0.0315 |
| groupH37Ra not infected | -0.1934 | 0.0796 | 0.0152 |
| groupH37Rv not infected | -0.0723 | 0.0801 | 0.3668 |
| groupAfri infected | 0.2767 | 0.0697 | 1e-04 |
| groupBCG infected | 0.097 | 0.1051 | 0.3562 |
| groupBeijing infected | 0.2043 | 0.064 | 0.0014 |
| groupCDC1551 infected | 0.1113 | 0.0776 | 0.1516 |
| groupEAI infected | 0.3083 | 0.0752 | 0 |
| groupH37Ra infected | -0.112 | 0.08 | 0.1614 |
| groupH37Rv infected | 0.0413 | 0.0806 | 0.6083 |

| Contrast | estimate | SE | p.value |
|---|---|---|---|
| none not infected-Afri not infected | -0.1295 | 0.079 | 0.2473 |
| none not infected-BCG not infected | -0.0375 | 0.1119 | 0.7622 |
| none not infected-Beijing not infected | -0.101 | 0.0743 | 0.3244 |
| none not infected-CDC1551 not infected | 0.0356 | 0.086 | 0.7407 |
| none not infected-EAI not infected | -0.1671 | 0.084 | 0.1734 |
| none not infected-H37Ra not infected | 0.1876 | 0.0873 | 0.1516 |
| none not infected-H37Rv not infected | 0.0665 | 0.0877 | 0.5734 |
| none not infected-Afri infected | -0.2825 | 0.0791 | 0.0403 |
| none not infected-BCG infected | -0.1028 | 0.1118 | 0.5209 |
| none not infected-Beijing infected | -0.2101 | 0.0742 | 0.0799 |
| none not infected-CDC1551 infected | -0.1171 | 0.0862 | 0.3244 |
| none not infected-EAI infected | -0.3141 | 0.0842 | 0.0356 |
| none not infected-H37Ra infected | 0.1062 | 0.0877 | 0.3793 |
| none not infected-H37Rv infected | -0.0471 | 0.0882 | 0.6986 |
| Afri not infected-BCG not infected | 0.092 | 0.1167 | 0.5427 |
| Afri not infected-Beijing not infected | 0.0285 | 0.0724 | 0.7407 |
| Afri not infected-CDC1551 not infected | 0.1651 | 0.0858 | 0.1334 |
| Afri not infected-EAI not infected | -0.0376 | 0.0853 | 0.7315 |
| Afri not infected-H37Ra not infected | 0.317 | 0.0932 | 0.0039 |
| Afri not infected-H37Rv not infected | 0.196 | 0.096 | 0.1135 |
| BCG not infected-Beijing not infected | -0.0635 | 0.1143 | 0.6752 |
| BCG not infected-CDC1551 not infected | 0.0732 | 0.1243 | 0.6612 |
| BCG not infected-EAI not infected | -0.1296 | 0.0989 | 0.3074 |
| BCG not infected-H37Ra not infected | 0.2251 | 0.1265 | 0.1585 |
| BCG not infected-H37Rv not infected | 0.104 | 0.127 | 0.5422 |
| Beijing not infected-CDC1551 not infected | 0.1366 | 0.0802 | 0.1734 |
| Beijing not infected-EAI not infected | -0.0662 | 0.0827 | 0.5427 |
| Beijing not infected-H37Ra not infected | 0.2885 | 0.0891 | 0.0064 |
| Beijing not infected-H37Rv not infected | 0.1674 | 0.0895 | 0.1436 |
| CDC1551 not infected-EAI not infected | -0.2028 | 0.098 | 0.1106 |
| CDC1551 not infected-H37Ra not infected | 0.1519 | 0.0968 | 0.2102 |
| CDC1551 not infected-H37Rv not infected | 0.0308 | 0.0968 | 0.7622 |
| EAI not infected-H37Ra not infected | 0.3547 | 0.1017 | 0.0035 |
| EAI not infected-H37Rv not infected | 0.2336 | 0.1025 | 0.0754 |
| H37Ra not infected-H37Rv not infected | -0.1211 | 0.0898 | 0.2947 |
| Afri infected-BCG infected | 0.1798 | 0.1167 | 0.2162 |
| Afri infected-Beijing infected | 0.0725 | 0.0725 | 0.4445 |
| Afri infected-CDC1551 infected | 0.1654 | 0.0862 | 0.1334 |
| Afri infected-EAI infected | -0.0316 | 0.0856 | 0.7478 |
| Afri infected-H37Ra infected | 0.3888 | 0.0936 | 4e-04 |
| Afri infected-H37Rv infected | 0.2354 | 0.0966 | 0.0551 |
| BCG infected-Beijing infected | -0.1073 | 0.1141 | 0.4756 |
| BCG infected-CDC1551 infected | -0.0143 | 0.1244 | 0.9083 |
| BCG infected-EAI infected | -0.2113 | 0.099 | 0.0987 |
| BCG infected-H37Ra infected | 0.209 | 0.1267 | 0.1837 |
| BCG infected-H37Rv infected | 0.0557 | 0.1273 | 0.7315 |
| Beijing infected-CDC1551 infected | 0.093 | 0.0803 | 0.3621 |
| Beijing infected-EAI infected | -0.1041 | 0.0827 | 0.3244 |

### b) Percentage of cells showing acidified compartments

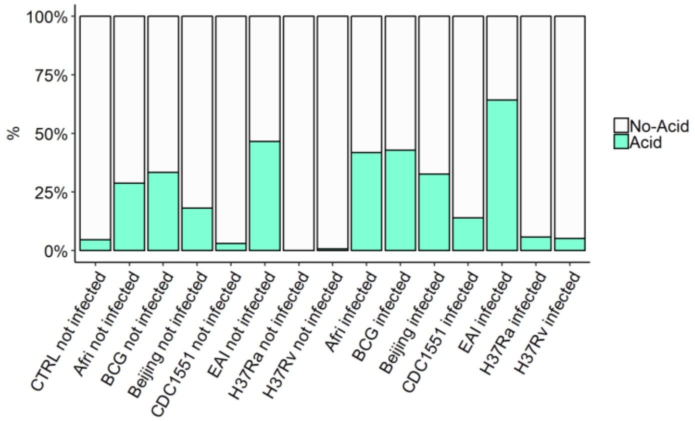


| Par | Estimate | se | OR | pvalue |
|---|---|---|---|---|
| (Intercept) | -3.505 | 0.7631 | 0.03 | 0 |
| groupAfri not infected | 1.8572 | 0.7306 | 6.4057 | 0.011 |
| groupBCG not infected | 0.8416 | 0.9631 | 2.32 | 0.3822 |
| groupBeijing not infected | 1.1595 | 0.7357 | 3.1882 | 0.115 |
| groupCDC1551 not infected | 0.1986 | 0.9662 | 1.2197 | 0.8371 |
| groupEAI not infected | 1.7238 | 0.7466 | 5.6058 | 0.021 |
| groupH37Ra not infected | -14.8469 | 583.2223 | 0 | 0.9797 |
| groupH37Rv not infected | -1.2751 | 1.3749 | 0.2794 | 0.3537 |
| groupAfri infected | 2.998 | 0.7254 | 20.046 | 0 |
| groupBCG infected | 1.25 | 0.9578 | 3.4903 | 0.1919 |
| groupBeijing infected | 2.6757 | 0.7166 | 14.5219 | 2e-04 |
| groupCDC1551 infected | 1.9659 | 0.8611 | 7.1411 | 0.0224 |
| groupEAI infected | 2.8677 | 0.7499 | 17.5963 | 1e-04 |
| groupH37Ra infected | 0.8593 | 0.9051 | 2.3614 | 0.3424 |
| groupH37Rv infected | 0.723 | 1.0227 | 2.0606 | 0.4796 |

| Contrast | estimate | SE | p.value |
|---|---|---|---|
| none not infected-Afri not infected | 1.6478 | 0.7801 | 0.1456 |
| none not infected-BCG not infected | 2.6634 | 1.0411 | 0.0663 |
| none not infected-Beijing not infected | 2.3455 | 0.777 | 0.0239 |
| none not infected-CDC1551 not infected | 3.3063 | 0.9888 | 0.0104 |
| none not infected-EAI not infected | 1.7812 | 0.8266 | 0.1403 |
| none not infected-H37Ra not infected | 18.3519 | 583.2224 | 0.9814 |
| none not infected-H37Rv not infected | 4.7801 | 1.3959 | 0.0097 |
| none not infected-Afri infected | 0.5069 | 0.7747 | 0.718 |
| none not infected-BCG infected | 2.255 | 1.0362 | 0.1403 |
| none not infected-Beijing infected | 0.8293 | 0.7559 | 0.5029 |
| none not infected-CDC1551 infected | 1.5391 | 0.8854 | 0.2071 |
| none not infected-EAI infected | 0.6373 | 0.828 | 0.6468 |
| none not infected-H37Ra infected | 2.6457 | 0.9386 | 0.038 |
| none not infected-H37Rv infected | 2.782 | 1.0503 | 0.0566 |
| Afri not infected-BCG not infected | 1.0156 | 1.035 | 0.5274 |
| Afri not infected-Beijing not infected | 0.6977 | 0.6778 | 0.5029 |
| Afri not infected-CDC1551 not infected | 1.6586 | 0.9211 | 0.1917 |
| Afri not infected-EAI not infected | 0.1334 | 0.7717 | 0.9814 |
| Afri not infected-H37Ra not infected | 16.7041 | 583.2224 | 0.9814 |
| Afri not infected-H37Rv not infected | 3.1323 | 1.4441 | 0.1403 |
| BCG not infected-Beijing not infected | -0.3179 | 1.0413 | 0.939 |
| BCG not infected-CDC1551 not infected | 0.6429 | 1.2377 | 0.8089 |
| BCG not infected-EAI not infected | -0.8822 | 0.837 | 0.5029 |
| BCG not infected-H37Ra not infected | 15.6885 | 583.2229 | 0.9814 |
| BCG not infected-H37Rv not infected | 2.1167 | 1.6078 | 0.3948 |
| Beijing not infected-CDC1551 not infected | 0.9608 | 0.9201 | 0.5029 |
| Beijing not infected-EAI not infected | -0.5643 | 0.7826 | 0.6742 |
| Beijing not infected-H37Ra not infected | 16.0064 | 583.2224 | 0.9814 |
| Beijing not infected-H37Rv not infected | 2.4346 | 1.4383 | 0.2112 |
| CDC1551 not infected-EAI not infected | -1.5252 | 1.043 | 0.3232 |
| CDC1551 not infected-H37Ra not infected | 15.0455 | 583.2227 | 0.9814 |
| CDC1551 not infected-H37Rv not infected | 1.4737 | 1.5552 | 0.5275 |
| EAI not infected-H37Ra not infected | 16.5707 | 583.2225 | 0.9814 |
| EAI not infected-H37Rv not infected | 2.9989 | 1.4773 | 0.157 |
| H37Ra not infected-H37Rv not infected | -13.5718 | 583.2232 | 0.9814 |
| Afri infected-BCG infected | 1.748 | 1.0262 | 0.2112 |
| Afri infected-Beijing infected | 0.3224 | 0.6516 | 0.8147 |
| Afri infected-CDC1551 infected | 1.0322 | 0.8074 | 0.4087 |
| Afri infected-EAI infected | 0.1303 | 0.7644 | 0.9814 |
| Afri infected-H37Ra infected | 2.1388 | 0.9561 | 0.1403 |
| Afri infected-H37Rv infected | 2.275 | 1.109 | 0.157 |
| BCG infected-Beijing infected | -1.4257 | 1.0247 | 0.3566 |
| BCG infected-CDC1551 infected | -0.7159 | 1.1531 | 0.7323 |
| BCG infected-EAI infected | -1.6177 | 0.8452 | 0.1752 |
| BCG infected-H37Ra infected | 0.3907 | 1.2151 | 0.939 |
| BCG infected-H37Rv infected | 0.527 | 1.3151 | 0.8854 |
| Beijing infected-CDC1551 infected | 0.7098 | 0.7852 | 0.549 |
| Beijing infected-EAI infected | -0.192 | 0.767 | 0.972 |

**M1 – 6 h p.i.**

c) Effective number of mycobacteria per cell (M/C)

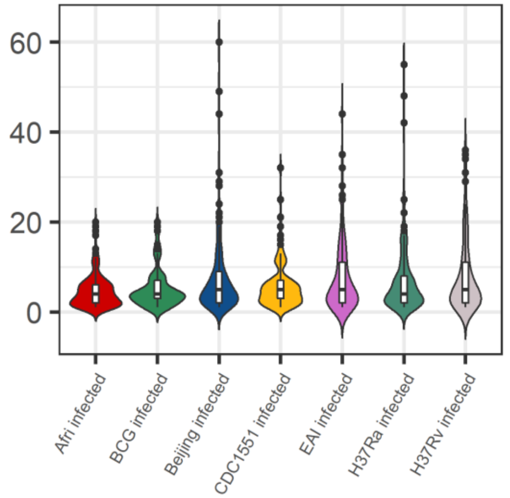


| Par | Value | Std.Error | p.value |
|---|---|---|---|
| (Intercept) | 1.1618 | 0.1532 | 0 |
| groupBCG infected | 0.5123 | 0.2551 | 0.0493 |
| groupBeijing infected | 0.2666 | 0.146 | 0.0729 |
| groupCDC1551 infected | 0.0771 | 0.1751 | 0.6612 |
| groupEAI infected | 0.6082 | 0.183 | 0.0015 |
| groupH37Ra infected | 0.3031 | 0.1976 | 0.1306 |
| groupH37Rv infected | 0.2976 | 0.2059 | 0.1539 |

| Contrast | estimate | SE | p.value |
|---|---|---|---|
| Afri infected - BCG infected | -0.5123 | 0.2551 | 0.3061 |
| Afri infected - Beijing infected | -0.2666 | 0.146 | 0.3061 |
| Afri infected - CDC1551 infected | -0.0771 | 0.1751 | 0.7713 |
| Afri infected - EAI infected | -0.6082 | 0.183 | 0.0324 |
| Afri infected - H37Ra infected | -0.3031 | 0.1976 | 0.3867 |
| Afri infected - H37Rv infected | -0.2976 | 0.2059 | 0.3867 |
| BCG infected - Beijing infected | 0.2457 | 0.2511 | 0.498 |
| BCG infected - CDC1551 infected | 0.4352 | 0.2743 | 0.3867 |
| BCG infected - EAI infected | -0.0959 | 0.2033 | 0.7713 |
| BCG infected - H37Ra infected | 0.2092 | 0.2835 | 0.6083 |
| BCG infected - H37Rv infected | 0.2148 | 0.2855 | 0.6083 |
| Beijing infected - CDC1551 infected | 0.1895 | 0.1614 | 0.4502 |
| Beijing infected - EAI infected | -0.3416 | 0.1787 | 0.3061 |
| Beijing infected - H37Ra infected | -0.0364 | 0.1881 | 0.9146 |
| Beijing infected - H37Rv infected | -0.0309 | 0.1897 | 0.9146 |
| CDC1551 infected - EAI infected | -0.5311 | 0.2133 | 0.1645 |
| CDC1551 infected - H37Ra infected | -0.226 | 0.2009 | 0.4502 |
| CDC1551 infected - H37Rv infected | -0.2204 | 0.2016 | 0.4502 |
| EAI infected - H37Ra infected | 0.3051 | 0.227 | 0.3867 |
| EAI infected - H37Rv infected | 0.3107 | 0.2299 | 0.3867 |
| H37Ra infected - H37Rv infected | 0.0055 | 0.1847 | 0.9763 |

d) Colocalization Index

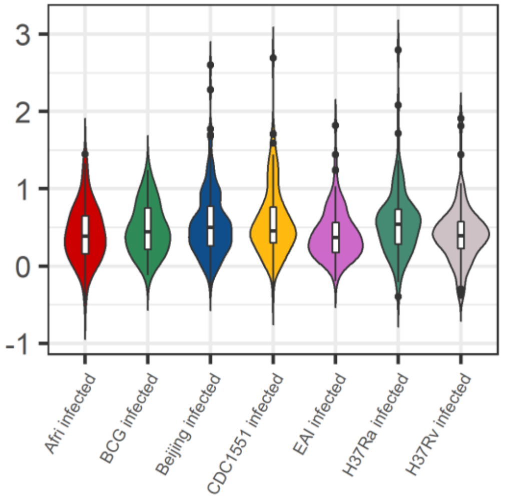


| Par | Value | Std.Error | p.value |
|---|---|---|---|
| (Intercept) | -0.009 | 0.1222 | 0.9412 |
| groupBCG infected | 0.0397 | 0.2109 | 0.8513 |
| groupBeijing infected | 0.2322 | 0.1282 | 0.0752 |
| groupCDC1551 infected | 0.1699 | 0.1498 | 0.2613 |
| groupEAI infected | -0.0855 | 0.1555 | 0.5845 |
| groupH37Ra infected | 0.2281 | 0.1663 | 0.1756 |
| groupH37Rv infected | -0.2072 | 0.1721 | 0.2335 |

| Contrast | estimate | SE | p.value |
|---|---|---|---|
| Afri infected - BCG infected | -0.0397 | 0.2109 | 0.8939 |
| Afri infected - Beijing infected | -0.2322 | 0.1282 | 0.3157 |
| Afri infected - CDC1551 infected | -0.1699 | 0.1498 | 0.5486 |
| Afri infected - EAI infected | 0.0855 | 0.1555 | 0.722 |
| Afri infected - H37Ra infected | -0.2281 | 0.1663 | 0.4609 |
| Afri infected - H37Rv infected | 0.2072 | 0.1721 | 0.5449 |
| BCG infected - Beijing infected | -0.1925 | 0.2068 | 0.6223 |
| BCG infected - CDC1551 infected | -0.1302 | 0.2242 | 0.722 |
| BCG infected - EAI infected | 0.1252 | 0.173 | 0.7081 |
| BCG infected - H37Ra infected | -0.1883 | 0.231 | 0.6756 |
| BCG infected - H37Rv infected | 0.2469 | 0.2313 | 0.5539 |
| Beijing infected - CDC1551 infected | 0.0623 | 0.1381 | 0.7623 |
| Beijing infected - EAI infected | 0.3177 | 0.151 | 0.2084 |
| Beijing infected - H37Ra infected | 0.0042 | 0.1592 | 0.9791 |
| Beijing infected - H37Rv infected | 0.4395 | 0.1583 | 0.0962 |
| CDC1551 infected - EAI infected | 0.2554 | 0.1773 | 0.4609 |
| CDC1551 infected - H37Ra infected | -0.0581 | 0.1698 | 0.8106 |
| CDC1551 infected - H37Rv infected | 0.3771 | 0.1691 | 0.2076 |
| EAI infected - H37Ra infected | -0.3135 | 0.1874 | 0.349 |
| EAI infected - H37Rv infected | 0.1217 | 0.1882 | 0.722 |
| H37Ra infected - H37Rv infected | 0.4353 | 0.1614 | 0.0962 |

# M1 – 24 h p.i.

### a) Acifdification of LysoSensor⁺ compartments

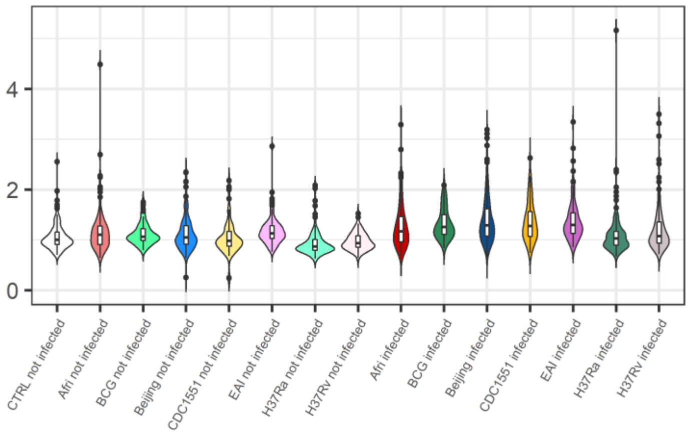


### b) Percentage of cells showing acidified compartments

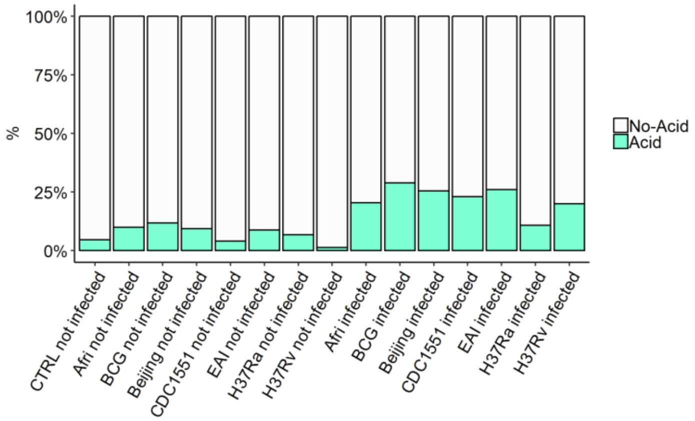


| Par | Value | Std.Error | p.value |
|---|---|---|---|
| (Intercept) | 0.0184 | 0.0353 | 0.6021 |
| groupAfri not infected | 0.0221 | 0.0389 | 0.5709 |
| groupBCG not infected | 0.0757 | 0.0572 | 0.1858 |
| groupBeijing not infected | 0.0485 | 0.0368 | 0.1871 |
| groupCDC1551 not infected | 0.0033 | 0.0444 | 0.9402 |
| groupEAI not infected | 0.0742 | 0.0406 | 0.0675 |
| groupH37Ra not infected | -0.0957 | 0.0466 | 0.0403 |
| groupH37Rv not infected | -0.0249 | 0.0462 | 0.5902 |
| groupAfri infected | 0.1127 | 0.0389 | 0.0038 |
| groupBCG infected | 0.2329 | 0.0548 | 0 |
| groupBeijing infected | 0.2026 | 0.0358 | 0 |
| groupCDC1551 infected | 0.2551 | 0.0435 | 0 |
| groupEAI infected | 0.1992 | 0.0408 | 0 |
| groupH37Ra infected | 0.0394 | 0.0448 | 0.3793 |
| groupH37Rv infected | 0.1119 | 0.045 | 0.0129 |

| Par | Estimate | se | OR | pvalue |
|---|---|---|---|---|
| (Intercept) | -3.1573 | 0.4472 | 0.0425 | 0 |
| groupAfri not infected | 0.6976 | 0.5452 | 2.0088 | 0.2007 |
| groupBCG not infected | 0.8584 | 0.683 | 2.3594 | 0.2088 |
| groupBeijing not infected | 0.6945 | 0.5348 | 2.0028 | 0.1941 |
| groupCDC1551 not infected | 0.0581 | 0.7516 | 1.0598 | 0.9384 |
| groupEAI not infected | 0.4412 | 0.5449 | 1.5546 | 0.4181 |
| groupH37Ra not infected | 0.9405 | 0.6736 | 2.5612 | 0.1626 |
| groupH37Rv not infected | -1.0996 | 1.115 | 0.333 | 0.324 |
| groupAfri infected | 1.6261 | 0.5024 | 5.0838 | 0.0012 |
| groupBCG infected | 1.9978 | 0.5825 | 7.3727 | 6e-04 |
| groupBeijing infected | 1.9321 | 0.4719 | 6.9042 | 0 |
| groupCDC1551 infected | 1.9848 | 0.5336 | 7.2773 | 2e-04 |
| groupEAI infected | 1.7757 | 0.496 | 5.9046 | 3e-04 |
| groupH37Ra infected | 1.422 | 0.5799 | 4.1452 | 0.0142 |
| groupH37Rv infected | 1.855 | 0.5356 | 6.3916 | 5e-04 |

| Contrast | estimate | SE | p.value |
|---|---|---|---|
| none not infected-Afri not infected | -0.0405 | 0.0408 | 0.4642 |
| none not infected-BCG not infected | -0.0941 | 0.0591 | 0.2506 |
| none not infected-Beijing not infected | -0.067 | 0.0387 | 0.2413 |
| none not infected-CDC1551 not infected | -0.0218 | 0.0459 | 0.7302 |
| none not infected-EAI not infected | -0.0927 | 0.0429 | 0.1486 |
| none not infected-H37Ra not infected | 0.0773 | 0.047 | 0.2436 |
| none not infected-H37Rv not infected | 0.0065 | 0.0467 | 0.9384 |
| none not infected-Afri infected | -0.1312 | 0.0407 | 0.0438 |
| none not infected-BCG infected | -0.2514 | 0.0568 | 0.0128 |
| none not infected-Beijing infected | -0.2211 | 0.0378 | 0.0039 |
| none not infected-CDC1551 infected | -0.2735 | 0.045 | 0.0039 |
| none not infected-EAI infected | -0.2176 | 0.0431 | 0.0077 |
| none not infected-H37Ra infected | -0.0578 | 0.0452 | 0.354 |
| none not infected-H37Rv infected | -0.1303 | 0.0454 | 0.0631 |
| Afri not infected-BCG not infected | -0.0536 | 0.0626 | 0.4942 |
| Afri not infected-Beijing not infected | -0.0265 | 0.0399 | 0.6188 |
| Afri not infected-CDC1551 not infected | 0.0187 | 0.0479 | 0.7557 |
| Afri not infected-EAI not infected | -0.0522 | 0.0453 | 0.3578 |
| Afri not infected-H37Ra not infected | 0.1178 | 0.052 | 0.0631 |
| Afri not infected-H37Rv not infected | 0.047 | 0.0529 | 0.4823 |
| BCG not infected-Beijing not infected | 0.0271 | 0.0618 | 0.7302 |
| BCG not infected-CDC1551 not infected | 0.0723 | 0.0677 | 0.3947 |
| BCG not infected-EAI not infected | 0.0014 | 0.0535 | 0.9868 |
| BCG not infected-H37Ra not infected | 0.1714 | 0.0694 | 0.0431 |
| BCG not infected-H37Rv not infected | 0.1006 | 0.0693 | 0.2436 |
| Beijing not infected-CDC1551 not infected | 0.0452 | 0.0453 | 0.4275 |
| Beijing not infected-EAI not infected | -0.0257 | 0.0445 | 0.6703 |
| Beijing not infected-H37Ra not infected | 0.1443 | 0.0502 | 0.0144 |
| Beijing not infected-H37Rv not infected | 0.0734 | 0.0501 | 0.2436 |
| CDC1551 not infected-EAI not infected | -0.0709 | 0.0532 | 0.2814 |
| CDC1551 not infected-H37Ra not infected | 0.099 | 0.0548 | 0.1486 |
| CDC1551 not infected-H37Rv not infected | 0.0282 | 0.0544 | 0.6959 |
| EAI not infected-H37Ra not infected | 0.17 | 0.0558 | 0.0114 |
| EAI not infected-H37Rv not infected | 0.0991 | 0.0557 | 0.1535 |
| H37Ra not infected-H37Rv not infected | -0.0708 | 0.0509 | 0.2588 |
| Afri infected-BCG infected | -0.1202 | 0.0605 | 0.1186 |
| Afri infected-Beijing infected | -0.0899 | 0.0392 | 0.0625 |
| Afri infected-CDC1551 infected | -0.1424 | 0.0471 | 0.0115 |
| Afri infected-EAI infected | -0.0864 | 0.0455 | 0.1375 |
| Afri infected-H37Ra infected | 0.0733 | 0.0504 | 0.2436 |
| Afri infected-H37Rv infected | 9e-04 | 0.0518 | 0.9868 |
| BCG infected-Beijing infected | 0.0303 | 0.059 | 0.6959 |
| BCG infected-CDC1551 infected | -0.0222 | 0.0651 | 0.7835 |
| BCG infected-EAI infected | 0.0338 | 0.0513 | 0.6188 |
| BCG infected-H37Ra infected | 0.1935 | 0.0663 | 0.0132 |
| BCG infected-H37Rv infected | 0.1211 | 0.0665 | 0.1486 |
| Beijing infected-CDC1551 infected | -0.0525 | 0.0438 | 0.3465 |
| Beijing infected-EAI infected | 0.0035 | 0.0438 | 0.9676 |

| Contrast | estimate | SE | p.value |
|---|---|---|---|
| none not infected-Afri not infected | 2.4597 | 0.4095 | 0 |
| none not infected-BCG not infected | 2.2988 | 0.5951 | 6e-04 |
| none not infected-Beijing not infected | 2.4627 | 0.3939 | 0 |
| none not infected-CDC1551 not infected | 3.0991 | 0.6548 | 0 |
| none not infected-EAI not infected | 2.716 | 0.4226 | 0 |
| none not infected-H37Ra not infected | 2.2168 | 0.5547 | 5e-04 |
| none not infected-H37Rv not infected | 4.2569 | 1.0504 | 5e-04 |
| none not infected-Afri infected | 1.5312 | 0.349 | 1e-04 |
| none not infected-BCG infected | 1.1595 | 0.4765 | 0.0524 |
| none not infected-Beijing infected | 1.2251 | 0.3016 | 5e-04 |
| none not infected-CDC1551 infected | 1.1725 | 0.3853 | 0.0098 |
| none not infected-EAI infected | 1.3815 | 0.3567 | 6e-04 |
| none not infected-H37Ra infected | 1.7353 | 0.4357 | 5e-04 |
| none not infected-H37Rv infected | 1.3023 | 0.3821 | 0.003 |
| Afri not infected-BCG not infected | -0.1609 | 0.6796 | 0.9145 |
| Afri not infected-Beijing not infected | 0.003 | 0.4857 | 0.995 |
| Afri not infected-CDC1551 not infected | 0.6394 | 0.7259 | 0.6934 |
| Afri not infected-EAI not infected | 0.2563 | 0.5149 | 0.8655 |
| Afri not infected-H37Ra not infected | -0.2429 | 0.6584 | 0.8697 |
| Afri not infected-H37Rv not infected | 1.7972 | 1.1098 | 0.2766 |
| BCG not infected-Beijing not infected | 0.1639 | 0.6755 | 0.9145 |
| BCG not infected-CDC1551 not infected | 0.8003 | 0.8647 | 0.6934 |
| BCG not infected-EAI not infected | 0.4172 | 0.6115 | 0.7797 |
| BCG not infected-H37Ra not infected | -0.082 | 0.7989 | 0.9483 |
| BCG not infected-H37Rv not infected | 1.958 | 1.1967 | 0.2766 |
| Beijing not infected-CDC1551 not infected | 0.6364 | 0.71 | 0.6934 |
| Beijing not infected-EAI not infected | 0.2533 | 0.5127 | 0.8655 |
| Beijing not infected-H37Ra not infected | -0.2459 | 0.6453 | 0.8697 |
| Beijing not infected-H37Rv not infected | 1.7942 | 1.0982 | 0.2766 |
| CDC1551 not infected-EAI not infected | -0.3831 | 0.7501 | 0.8655 |
| CDC1551 not infected-H37Ra not infected | -0.8823 | 0.8231 | 0.6585 |
| CDC1551 not infected-H37Rv not infected | 1.1577 | 1.2072 | 0.6934 |
| EAI not infected-H37Ra not infected | -0.4992 | 0.6765 | 0.7635 |
| EAI not infected-H37Rv not infected | 1.5408 | 1.1186 | 0.4243 |
| H37Ra not infected-H37Rv not infected | 2.0401 | 1.1464 | 0.2255 |
| Afri infected-BCG infected | -0.3717 | 0.5393 | 0.7797 |
| Afri infected-Beijing infected | -0.3061 | 0.3608 | 0.6934 |
| Afri infected-CDC1551 infected | -0.3587 | 0.4494 | 0.7232 |
| Afri infected-EAI infected | -0.1497 | 0.4142 | 0.8697 |
| Afri infected-H37Ra infected | 0.2041 | 0.5203 | 0.8697 |
| Afri infected-H37Rv infected | -0.2289 | 0.478 | 0.8655 |
| BCG infected-Beijing infected | 0.0657 | 0.5166 | 0.9438 |
| BCG infected-CDC1551 infected | 0.013 | 0.5839 | 0.995 |
| BCG infected-EAI infected | 0.2221 | 0.4403 | 0.8655 |
| BCG infected-H37Ra infected | 0.5758 | 0.6279 | 0.6934 |
| BCG infected-H37Rv infected | 0.1428 | 0.59 | 0.9145 |
| Beijing infected-CDC1551 infected | -0.0526 | 0.397 | 0.9438 |
| Beijing infected-EAI infected | 0.1564 | 0.3882 | 0.8697 |

# M1 – 24 h p.i.

c) Effective number of mycobacteria per cell (M/C)

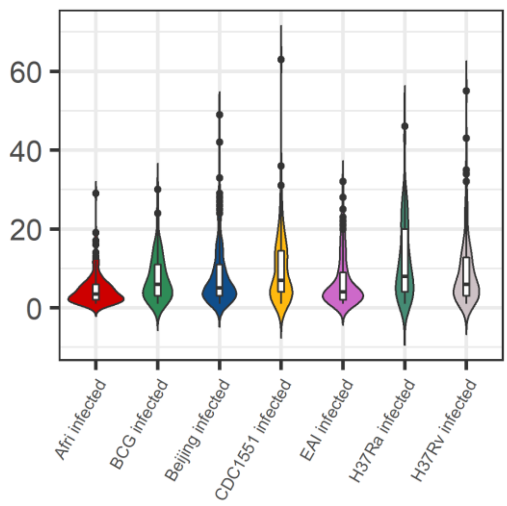


| Par | Value | Std.Error | p.value |
|---|---|---|---|
| (Intercept) | 1.3459 | 0.1565 | 0 |
| groupBCG infected | 0.8519 | 0.2039 | 1e-04 |
| groupBeijing infected | 0.3098 | 0.1109 | 0.0071 |
| groupCDC1551 infected | 0.367 | 0.1412 | 0.0119 |
| groupEAI infected | 0.3906 | 0.1448 | 0.0091 |
| groupH37Ra infected | 0.7005 | 0.162 | 1e-04 |
| groupH37Rv infected | 0.1793 | 0.1664 | 0.2859 |

| Contrast | estimate | SE | p.value |
|---|---|---|---|
| Afri infected - BCG infected | -0.8519 | 0.2039 | 0.0011 |
| Afri infected - Beijing infected | -0.3098 | 0.1109 | 0.0247 |
| Afri infected - CDC1551 infected | -0.367 | 0.1412 | 0.0256 |
| Afri infected - EAI infected | -0.3906 | 0.1448 | 0.0249 |
| Afri infected - H37Ra infected | -0.7005 | 0.162 | 0.0011 |
| Afri infected - H37Rv infected | -0.1793 | 0.1664 | 0.3753 |
| BCG infected - Beijing infected | 0.5421 | 0.202 | 0.0249 |
| BCG infected - CDC1551 infected | 0.4849 | 0.2259 | 0.0688 |
| BCG infected - EAI infected | 0.4613 | 0.1516 | 0.0185 |
| BCG infected - H37Ra infected | 0.1514 | 0.2369 | 0.6128 |
| BCG infected - H37Rv infected | 0.6726 | 0.2381 | 0.0247 |
| Beijing infected - CDC1551 infected | -0.0572 | 0.1262 | 0.6847 |
| Beijing infected - EAI infected | -0.0808 | 0.1427 | 0.6337 |
| Beijing infected - H37Ra infected | -0.3907 | 0.151 | 0.0256 |
| Beijing infected - H37Rv infected | 0.1305 | 0.1506 | 0.4812 |
| CDC1551 infected - EAI infected | -0.0236 | 0.1761 | 0.8938 |
| CDC1551 infected - H37Ra infected | -0.3335 | 0.1598 | 0.0723 |
| CDC1551 infected - H37Rv infected | 0.1877 | 0.1579 | 0.3591 |
| EAI infected - H37Ra infected | -0.3099 | 0.1911 | 0.1781 |
| EAI infected - H37Rv infected | 0.2113 | 0.1926 | 0.3753 |
| H37Ra infected - H37Rv infected | 0.5212 | 0.1322 | 0.0015 |

d) Colocalization Index

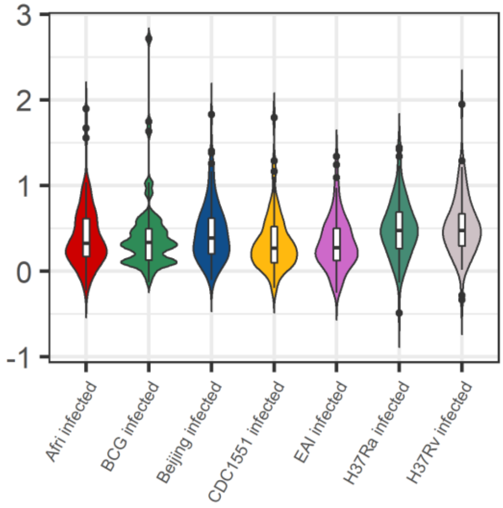


| Par | Value | Std.Error | p.value |
|---|---|---|---|
| (Intercept) | -0.1611 | 0.1235 | 0.1926 |
| groupBCG infected | -0.1594 | 0.1916 | 0.409 |
| groupBeijing infected | 0.131 | 0.1114 | 0.2443 |
| groupCDC1551 infected | -0.1753 | 0.1453 | 0.2326 |
| groupEAI infected | -0.2812 | 0.1399 | 0.0492 |
| groupH37Ra infected | 0.629 | 0.1562 | 2e-04 |
| groupH37Rv infected | 0.6642 | 0.1575 | 1e-04 |

| Contrast | estimate | SE | p.value |
|---|---|---|---|
| Afri infected - BCG infected | 0.1594 | 0.1916 | 0.482 |
| Afri infected - Beijing infected | -0.131 | 0.1114 | 0.3206 |
| Afri infected - CDC1551 infected | 0.1753 | 0.1453 | 0.3206 |
| Afri infected - EAI infected | 0.2812 | 0.1399 | 0.0795 |
| Afri infected - H37Ra infected | -0.629 | 0.1562 | 6e-04 |
| Afri infected - H37Rv infected | -0.6642 | 0.1575 | 4e-04 |
| BCG infected - Beijing infected | -0.2904 | 0.1889 | 0.1944 |
| BCG infected - CDC1551 infected | 0.0159 | 0.2133 | 0.9408 |
| BCG infected - EAI infected | 0.1218 | 0.1478 | 0.482 |
| BCG infected - H37Ra infected | -0.7884 | 0.2173 | 0.0014 |
| BCG infected - H37Rv infected | -0.8236 | 0.2164 | 0.001 |
| Beijing infected - CDC1551 infected | 0.3063 | 0.1345 | 0.0463 |
| Beijing infected - EAI infected | 0.4123 | 0.1368 | 0.0073 |
| Beijing infected - H37Ra infected | -0.498 | 0.1486 | 0.003 |
| Beijing infected - H37Rv infected | -0.5332 | 0.1463 | 0.0014 |
| CDC1551 infected - EAI infected | 0.1059 | 0.1717 | 0.5967 |
| CDC1551 infected - H37Ra infected | -0.8043 | 0.1586 | 0 |
| CDC1551 infected - H37Rv infected | -0.8395 | 0.1494 | 0 |
| EAI infected - H37Ra infected | -0.9102 | 0.1782 | 0 |
| EAI infected - H37Rv infected | -0.9454 | 0.1772 | 0 |
| H37Ra infected - H37Rv infected | -0.0352 | 0.1294 | 0.8259 |

## M2 – 6 h p.i.

a) Acifdification of LysoSensor⁺ compartments

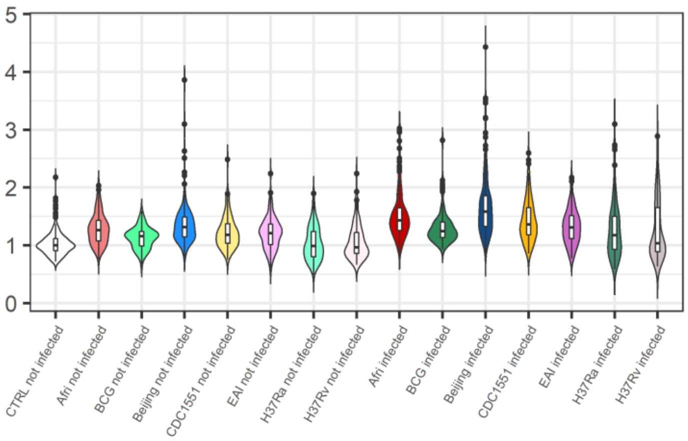


b) Percentage of cells showing acidified compartments

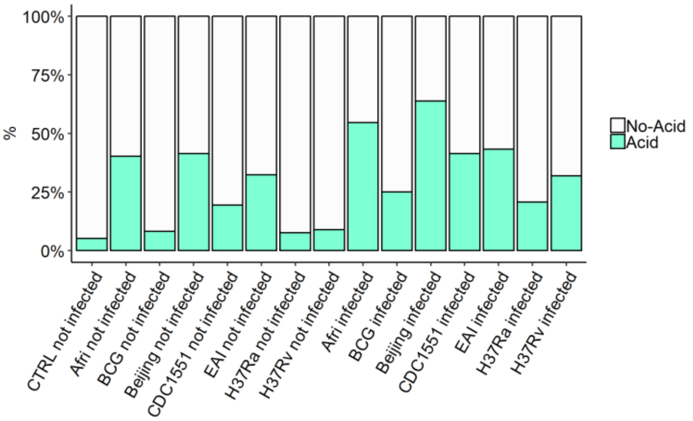


| Par | Value | Std.Error | p.value |
|---|---|---|---|
| (Intercept) | -0.0102 | 0.0427 | 0.8114 |
| groupAfri not infected | 0.2373 | 0.0532 | 0 |
| groupBCG not infected | 0.0963 | 0.0789 | 0.2224 |
| groupBeijing not infected | 0.2672 | 0.0491 | 0 |
| groupCDC1551 not infected | 0.1257 | 0.0587 | 0.0325 |
| groupEAI not infected | 0.1354 | 0.0569 | 0.0174 |
| groupH37Ra not infected | 0.0167 | 0.0593 | 0.7779 |
| groupH37Rv not infected | 0.0451 | 0.0599 | 0.4519 |
| groupAfri infected | 0.3544 | 0.0535 | 0 |
| groupBCG infected | 0.1968 | 0.0788 | 0.0125 |
| groupBeijing infected | 0.4182 | 0.0489 | 0 |
| groupCDC1551 infected | 0.2551 | 0.059 | 0 |
| groupEAI infected | 0.242 | 0.0569 | 0 |
| groupH37Ra infected | 0.178 | 0.0597 | 0.0029 |
| groupH37Rv infected | 0.1709 | 0.06 | 0.0045 |

| Contrast | estimate | SE | p.value |
|---|---|---|---|
| none not infected-Afri not infected | -0.2271 | 0.0533 | 0.0147 |
| none not infected-BCG not infected | -0.0861 | 0.079 | 0.4273 |
| none not infected-Beijing not infected | -0.257 | 0.0493 | 0.0055 |
| none not infected-CDC1551 not infected | -0.1155 | 0.059 | 0.1639 |
| none not infected-EAI not infected | -0.1252 | 0.0569 | 0.1297 |
| none not infected-H37Ra not infected | -0.0065 | 0.0597 | 0.9175 |
| none not infected-H37Rv not infected | -0.0349 | 0.0603 | 0.6881 |
| none not infected-Afri infected | -0.3442 | 0.0536 | 0.0019 |
| none not infected-BCG infected | -0.1866 | 0.0789 | 0.1048 |
| none not infected-Beijing infected | -0.408 | 0.049 | 6e-04 |
| none not infected-CDC1551 infected | -0.245 | 0.0592 | 0.0158 |
| none not infected-EAI infected | -0.2319 | 0.057 | 0.0158 |
| none not infected-H37Ra infected | -0.1679 | 0.0601 | 0.0651 |
| none not infected-H37Rv infected | -0.1607 | 0.0605 | 0.076 |
| Afri not infected-BCG not infected | 0.141 | 0.0873 | 0.1867 |
| Afri not infected-Beijing not infected | -0.0299 | 0.0555 | 0.6881 |
| Afri not infected-CDC1551 not infected | 0.1116 | 0.0653 | 0.1621 |
| Afri not infected-EAI not infected | 0.1019 | 0.0646 | 0.1906 |
| Afri not infected-H37Ra not infected | 0.2205 | 0.0696 | 0.0066 |
| Afri not infected-H37Rv not infected | 0.1922 | 0.0719 | 0.0232 |
| BCG not infected-Beijing not infected | -0.1709 | 0.0854 | 0.0988 |
| BCG not infected-CDC1551 not infected | -0.0294 | 0.0926 | 0.8301 |
| BCG not infected-EAI not infected | -0.0391 | 0.0754 | 0.6919 |
| BCG not infected-H37Ra not infected | 0.0796 | 0.0936 | 0.5107 |
| BCG not infected-H37Rv not infected | 0.0512 | 0.0941 | 0.6881 |
| Beijing not infected-CDC1551 not infected | 0.1415 | 0.0611 | 0.0522 |
| Beijing not infected-EAI not infected | 0.1318 | 0.0624 | 0.0784 |
| Beijing not infected-H37Ra not infected | 0.2504 | 0.0666 | 0.0012 |
| Beijing not infected-H37Rv not infected | 0.2221 | 0.0672 | 0.0047 |
| CDC1551 not infected-EAI not infected | -0.0097 | 0.0734 | 0.9175 |
| CDC1551 not infected-H37Ra not infected | 0.109 | 0.0726 | 0.2106 |
| CDC1551 not infected-H37Rv not infected | 0.0806 | 0.0729 | 0.3846 |
| EAI not infected-H37Ra not infected | 0.1187 | 0.0752 | 0.1906 |
| EAI not infected-H37Rv not infected | 0.0903 | 0.076 | 0.3606 |
| H37Ra not infected-H37Rv not infected | -0.0283 | 0.0686 | 0.7647 |
| Afri infected-BCG infected | 0.1576 | 0.0874 | 0.1409 |
| Afri infected-Beijing infected | -0.0638 | 0.0556 | 0.369 |
| Afri infected-CDC1551 infected | 0.0993 | 0.0658 | 0.2106 |
| Afri infected-EAI infected | 0.1124 | 0.0649 | 0.1591 |
| Afri infected-H37Ra infected | 0.1764 | 0.0702 | 0.0317 |
| Afri infected-H37Rv infected | 0.1835 | 0.0722 | 0.0305 |
| BCG infected-Beijing infected | -0.2214 | 0.0851 | 0.0268 |
| BCG infected-CDC1551 infected | -0.0583 | 0.0926 | 0.6667 |
| BCG infected-EAI infected | -0.0452 | 0.0753 | 0.6773 |
| BCG infected-H37Ra infected | 0.0188 | 0.0937 | 0.8981 |
| BCG infected-H37Rv infected | 0.0259 | 0.094 | 0.8501 |
| Beijing infected-CDC1551 infected | 0.1631 | 0.0612 | 0.0232 |
| Beijing infected-EAI infected | 0.1762 | 0.0623 | 0.0158 |

| Par | Estimate | se | OR | pvalue |
|---|---|---|---|---|
| (Intercept) | -3.1634 | 0.5967 | 0.0423 | 0 |
| groupAfri not infected | 2.4272 | 0.639 | 11.3268 | 1e-04 |
| groupBCG not infected | 0.4143 | 0.9521 | 1.5133 | 0.6634 |
| groupBeijing not infected | 2.3782 | 0.5865 | 10.7859 | 1e-04 |
| groupCDC1551 not infected | 1.0524 | 0.6893 | 2.8645 | 0.1268 |
| groupEAI not infected | 1.6846 | 0.6812 | 5.3902 | 0.0134 |
| groupH37Ra not infected | 1.351 | 0.7642 | 3.8612 | 0.0771 |
| groupH37Rv not infected | -0.0155 | 0.7984 | 0.9846 | 0.9845 |
| groupAfri infected | 3.4862 | 0.6419 | 32.6619 | 0 |
| groupBCG infected | 1.8422 | 0.8825 | 6.3104 | 0.0369 |
| groupBeijing infected | 3.6119 | 0.5845 | 37.0353 | 0 |
| groupCDC1551 infected | 2.579 | 0.6714 | 13.1845 | 1e-04 |
| groupEAI infected | 2.3989 | 0.6793 | 11.0112 | 4e-04 |
| groupH37Ra infected | 2.6859 | 0.7168 | 14.6711 | 2e-04 |
| groupH37Rv infected | 2.3284 | 0.7232 | 10.2615 | 0.0013 |

| Contrast | estimate | SE | p.value |
|---|---|---|---|
| none not infected-Afri not infected | 0.7362 | 0.6118 | 0.3897 |
| none not infected-BCG not infected | 2.749 | 0.9388 | 0.0239 |
| none not infected-Beijing not infected | 0.7851 | 0.5609 | 0.3384 |
| none not infected-CDC1551 not infected | 2.1109 | 0.6688 | 0.0153 |
| none not infected-EAI not infected | 1.4788 | 0.6612 | 0.114 |
| none not infected-H37Ra not infected | 1.8124 | 0.7267 | 0.0663 |
| none not infected-H37Rv not infected | 3.1789 | 0.7727 | 6e-04 |
| none not infected-Afri infected | -0.3229 | 0.6143 | 0.7402 |
| none not infected-BCG infected | 1.3212 | 0.8681 | 0.2881 |
| none not infected-Beijing infected | -0.4485 | 0.5584 | 0.5844 |
| none not infected-CDC1551 infected | 0.5843 | 0.6499 | 0.553 |
| none not infected-EAI infected | 0.7644 | 0.6589 | 0.4078 |
| none not infected-H37Ra infected | 0.4775 | 0.6767 | 0.6177 |
| none not infected-H37Rv infected | 0.835 | 0.6913 | 0.3897 |
| Afri not infected-BCG not infected | 2.0129 | 0.9882 | 0.1544 |
| Afri not infected-Beijing not infected | 0.0489 | 0.5653 | 0.9334 |
| Afri not infected-CDC1551 not infected | 1.3748 | 0.6868 | 0.1581 |
| Afri not infected-EAI not infected | 0.7426 | 0.6922 | 0.4501 |
| Afri not infected-H37Ra not infected | 1.0762 | 0.7976 | 0.3384 |
| Afri not infected-H37Rv not infected | 2.4427 | 0.8509 | 0.0258 |
| BCG not infected-Beijing not infected | -1.9639 | 0.9638 | 0.1544 |
| BCG not infected-CDC1551 not infected | -0.6381 | 1.0514 | 0.6853 |
| BCG not infected-EAI not infected | -1.2703 | 0.8324 | 0.2881 |
| BCG not infected-H37Ra not infected | -0.9367 | 1.1066 | 0.5799 |
| BCG not infected-H37Rv not infected | 0.4298 | 1.1354 | 0.8059 |
| Beijing not infected-CDC1551 not infected | 1.3258 | 0.631 | 0.1496 |
| Beijing not infected-EAI not infected | 0.6936 | 0.6611 | 0.4519 |
| Beijing not infected-H37Ra not infected | 1.0273 | 0.7566 | 0.3384 |
| Beijing not infected-H37Rv not infected | 2.3937 | 0.7908 | 0.0195 |
| CDC1551 not infected-EAI not infected | -0.6322 | 0.7953 | 0.5844 |
| CDC1551 not infected-H37Ra not infected | -0.2986 | 0.8218 | 0.8059 |
| CDC1551 not infected-H37Rv not infected | 1.0679 | 0.8434 | 0.3698 |
| EAI not infected-H37Ra not infected | 0.3336 | 0.873 | 0.8059 |
| EAI not infected-H37Rv not infected | 1.7001 | 0.91 | 0.1944 |
| H37Ra not infected-H37Rv not infected | 1.3665 | 0.8538 | 0.276 |
| Afri infected-BCG infected | 1.644 | 0.9248 | 0.2067 |
| Afri infected-Beijing infected | -0.1257 | 0.5665 | 0.8657 |
| Afri infected-CDC1551 infected | 0.9072 | 0.6688 | 0.3384 |
| Afri infected-EAI infected | 1.0873 | 0.6953 | 0.2856 |
| Afri infected-H37Ra infected | 0.8003 | 0.7498 | 0.4501 |
| Afri infected-H37Rv infected | 1.1578 | 0.7825 | 0.302 |
| BCG infected-Beijing infected | -1.7697 | 0.8937 | 0.1581 |
| BCG infected-CDC1551 infected | -0.7368 | 0.9763 | 0.6004 |
| BCG infected-EAI infected | -0.5567 | 0.7492 | 0.6004 |
| BCG infected-H37Ra infected | -0.8437 | 1.0131 | 0.5799 |
| BCG infected-H37Rv infected | -0.4862 | 1.0232 | 0.7544 |
| Beijing infected-CDC1551 infected | 1.0328 | 0.6103 | 0.2377 |
| Beijing infected-EAI infected | 1.213 | 0.6569 | 0.1945 |

## M2 – 6 h p.i.

#### c) Effective number of mycobacteria per cell (M/C)

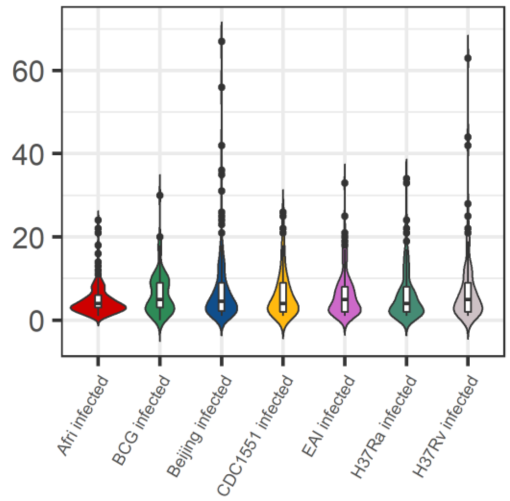


| Par | Value | Std.Error | p.value |
|---|---|---|---|
| (Intercept) | 0.0121 | 0.174 | 0.9446 |
| groupBCG infected | 0.229 | 0.2207 | 0.3039 |
| groupBeijing infected | 0.0098 | 0.1149 | 0.9324 |
| groupCDC1551 infected | -0.0838 | 0.1396 | 0.5507 |
| groupEAI infected | 0.2112 | 0.1578 | 0.186 |
| groupH37Ra infected | -0.3097 | 0.1632 | 0.0628 |
| groupH37Rv infected | -0.2894 | 0.176 | 0.1055 |

| Contrast | estimate | SE | p.value |
|---|---|---|---|
| Afri infected - BCG infected | -0.229 | 0.2207 | 0.4195 |
| Afri infected - Beijing infected | -0.0098 | 0.1149 | 0.9324 |
| Afri infected - CDC1551 infected | 0.0838 | 0.1396 | 0.6425 |
| Afri infected - EAI infected | -0.2112 | 0.1578 | 0.3211 |
| Afri infected - H37Ra infected | 0.3097 | 0.1632 | 0.1977 |
| Afri infected - H37Rv infected | 0.2894 | 0.176 | 0.2694 |
| BCG infected - Beijing infected | 0.2192 | 0.2183 | 0.4195 |
| BCG infected - CDC1551 infected | 0.3127 | 0.2395 | 0.3211 |
| BCG infected - EAI infected | 0.0178 | 0.1622 | 0.9324 |
| BCG infected - H37Ra infected | 0.5387 | 0.2516 | 0.1977 |
| BCG infected - H37Rv infected | 0.5184 | 0.2574 | 0.1977 |
| Beijing infected - CDC1551 infected | 0.0936 | 0.125 | 0.5648 |
| Beijing infected - EAI infected | -0.2014 | 0.1549 | 0.3211 |
| Beijing infected - H37Ra infected | 0.3195 | 0.1521 | 0.1977 |
| Beijing infected - H37Rv infected | 0.2992 | 0.1596 | 0.1977 |
| CDC1551 infected - EAI infected | -0.2949 | 0.1846 | 0.2694 |
| CDC1551 infected - H37Ra infected | 0.226 | 0.1612 | 0.3211 |
| CDC1551 infected - H37Rv infected | 0.2057 | 0.1685 | 0.3406 |
| EAI infected - H37Ra infected | 0.5209 | 0.2009 | 0.1977 |
| EAI infected - H37Rv infected | 0.5006 | 0.2083 | 0.1977 |
| H37Ra infected - H37Rv infected | -0.0203 | 0.1445 | 0.9324 |

#### d) Colocalization Index

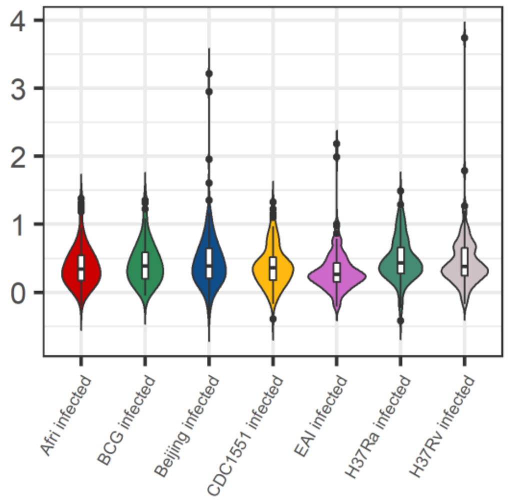


| Par | Value | Std.Error | p.value |
|---|---|---|---|
| (Intercept) | -0.2367 | 0.0657 | 3e-04 |
| groupBCG infected | 0.1292 | 0.1071 | 0.2323 |
| groupBeijing infected | 0.2211 | 0.084 | 0.0108 |
| groupCDC1551 infected | 0.054 | 0.0961 | 0.5762 |
| groupEAI infected | -0.1516 | 0.0943 | 0.1135 |
| groupH37Ra infected | 0.3735 | 0.0968 | 3e-04 |
| groupH37Rv infected | 0.1878 | 0.0981 | 0.0605 |

| Contrast | estimate | SE | p.value |
|---|---|---|---|
| Afri infected - BCG infected | -0.1292 | 0.1071 | 0.3049 |
| Afri infected - Beijing infected | -0.2211 | 0.084 | 0.036 |
| Afri infected - CDC1551 infected | -0.054 | 0.0961 | 0.6318 |
| Afri infected - EAI infected | 0.1516 | 0.0943 | 0.1702 |
| Afri infected - H37Ra infected | -0.3735 | 0.0968 | 0.002 |
| Afri infected - H37Rv infected | -0.1878 | 0.0981 | 0.1163 |
| BCG infected - Beijing infected | -0.0919 | 0.0994 | 0.4433 |
| BCG infected - CDC1551 infected | 0.0752 | 0.1098 | 0.5787 |
| BCG infected - EAI infected | 0.2808 | 0.1083 | 0.036 |
| BCG infected - H37Ra infected | -0.2443 | 0.1104 | 0.0811 |
| BCG infected - H37Rv infected | -0.0585 | 0.1116 | 0.6318 |
| Beijing infected - CDC1551 infected | 0.1671 | 0.0874 | 0.1163 |
| Beijing infected - EAI infected | 0.3727 | 0.0855 | 6e-04 |
| Beijing infected - H37Ra infected | -0.1524 | 0.0882 | 0.1445 |
| Beijing infected - H37Rv infected | 0.0334 | 0.0896 | 0.7111 |
| CDC1551 infected - EAI infected | 0.2056 | 0.0974 | 0.0913 |
| CDC1551 infected - H37Ra infected | -0.3195 | 0.0998 | 0.0093 |
| CDC1551 infected - H37Rv infected | -0.1338 | 0.1011 | 0.2673 |
| EAI infected - H37Ra infected | -0.5251 | 0.0981 | 0 |
| EAI infected - H37Rv infected | -0.3394 | 0.0994 | 0.0062 |
| H37Ra infected - H37Rv infected | 0.1858 | 0.1018 | 0.1278 |

# M2 – 24 h p.i.

a) Acifdification of LysoSensor+ compartments

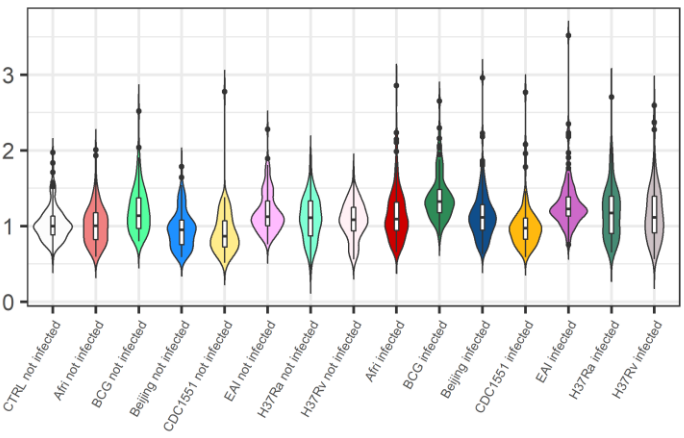


b) Percentage of cells showing acidified compartments

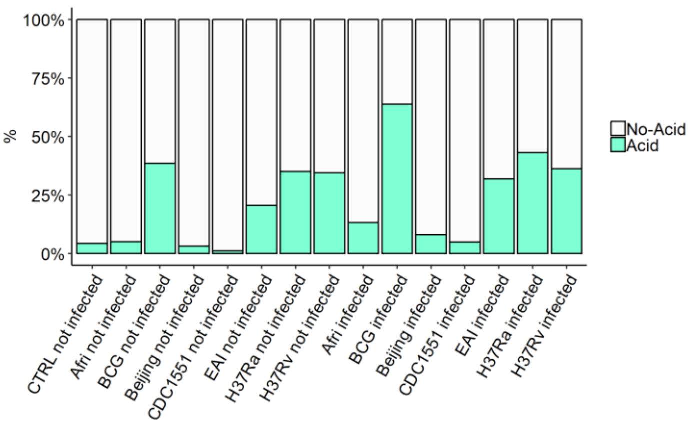


| Par | Value | Std.Error | p.value |
|---|---|---|---|
| (Intercept) | 0.0139 | 0.0526 | 0.7917 |
| groupAfri not infected | -0.0115 | 0.0369 | 0.7548 |
| groupBCG not infected | 0.0972 | 0.054 | 0.0724 |
| groupBeijing not infected | -0.0691 | 0.0351 | 0.0492 |
| groupCDC1551 not infected | -0.0907 | 0.0411 | 0.0275 |
| groupEAI not infected | 0.0767 | 0.0392 | 0.0506 |
| groupH37Ra not infected | -0.0554 | 0.044 | 0.2083 |
| groupH37Rv not infected | -0.1223 | 0.0446 | 0.0061 |
| groupAfri infected | 0.1065 | 0.0368 | 0.0039 |
| groupBCG infected | 0.2481 | 0.0534 | 0 |
| groupBeijing infected | 0.0653 | 0.034 | 0.055 |
| groupCDC1551 infected | -0.023 | 0.0416 | 0.5806 |
| groupEAI infected | 0.1561 | 0.0393 | 1e-04 |
| groupH37Ra infected | 0.0528 | 0.0426 | 0.2149 |
| groupH37Rv infected | -0.0563 | 0.0435 | 0.195 |

| Contrast | estimate | SE | p.value |
|---|---|---|---|
| none not infected-Afri not infected | -0.0024 | 0.056 | 0.9672 |
| none not infected-BCG not infected | -0.1111 | 0.0691 | 0.2339 |
| none not infected-Beijing not infected | 0.0552 | 0.0549 | 0.4292 |
| none not infected-CDC1551 not infected | 0.0768 | 0.0588 | 0.3256 |
| none not infected-EAI not infected | -0.0906 | 0.058 | 0.2376 |
| none not infected-H37Ra not infected | 0.0415 | 0.0602 | 0.5771 |
| none not infected-H37Rv not infected | 0.1084 | 0.0606 | 0.1932 |
| none not infected-Afri infected | -0.1203 | 0.056 | 0.1202 |
| none not infected-BCG infected | -0.262 | 0.0687 | 0.0207 |
| none not infected-Beijing infected | -0.0792 | 0.0541 | 0.2675 |
| none not infected-CDC1551 infected | 0.0091 | 0.0591 | 0.8966 |
| none not infected-EAI infected | -0.17 | 0.0581 | 0.0498 |
| none not infected-H37Ra infected | -0.0667 | 0.0592 | 0.3739 |
| none not infected-H37Rv infected | 0.0424 | 0.0598 | 0.5771 |
| Afri not infected-BCG not infected | -0.1087 | 0.0593 | 0.1202 |
| Afri not infected-Beijing not infected | 0.0576 | 0.0376 | 0.202 |
| Afri not infected-CDC1551 not infected | 0.0792 | 0.0441 | 0.1238 |
| Afri not infected-EAI not infected | -0.0882 | 0.0434 | 0.0835 |
| Afri not infected-H37Ra not infected | 0.0438 | 0.0495 | 0.4553 |
| Afri not infected-H37Rv not infected | 0.1108 | 0.0514 | 0.0664 |
| BCG not infected-Beijing not infected | 0.1663 | 0.0588 | 0.0158 |
| BCG not infected-CDC1551 not infected | 0.1879 | 0.064 | 0.0126 |
| BCG not infected-EAI not infected | 0.0205 | 0.049 | 0.7218 |
| BCG not infected-H37Ra not infected | 0.1525 | 0.0666 | 0.0498 |
| BCG not infected-H37Rv not infected | 0.2195 | 0.0672 | 0.0054 |
| Beijing not infected-CDC1551 not infected | 0.0216 | 0.0418 | 0.6575 |
| Beijing not infected-EAI not infected | -0.1458 | 0.043 | 0.0041 |
| Beijing not infected-H37Ra not infected | -0.0138 | 0.0478 | 0.8113 |
| Beijing not infected-H37Rv not infected | 0.0532 | 0.0488 | 0.3547 |
| CDC1551 not infected-EAI not infected | -0.1674 | 0.0507 | 0.0051 |
| CDC1551 not infected-H37Ra not infected | -0.0354 | 0.0509 | 0.5771 |
| CDC1551 not infected-H37Rv not infected | 0.0316 | 0.0515 | 0.5966 |
| EAI not infected-H37Ra not infected | 0.132 | 0.0542 | 0.0395 |
| EAI not infected-H37Rv not infected | 0.199 | 0.0551 | 0.0022 |
| H37Ra not infected-H37Rv not infected | 0.067 | 0.0474 | 0.2367 |
| Afri infected-BCG infected | -0.1417 | 0.0587 | 0.0395 |
| Afri infected-Beijing infected | 0.0411 | 0.0367 | 0.3514 |
| Afri infected-CDC1551 infected | 0.1294 | 0.0446 | 0.0131 |
| Afri infected-EAI infected | -0.0496 | 0.0434 | 0.3468 |
| Afri infected-H37Ra infected | 0.0536 | 0.0484 | 0.3514 |
| Afri infected-H37Rv infected | 0.1628 | 0.0504 | 0.0057 |
| BCG infected-Beijing infected | 0.1828 | 0.0576 | 0.0064 |
| BCG infected-CDC1551 infected | 0.2711 | 0.0638 | 3e-04 |
| BCG infected-EAI infected | 0.092 | 0.0484 | 0.1061 |
| BCG infected-H37Ra infected | 0.1953 | 0.0652 | 0.011 |
| BCG infected-H37Rv infected | 0.3045 | 0.066 | 1e-04 |
| Beijing infected-CDC1551 infected | 0.0883 | 0.0414 | 0.0674 |
| Beijing infected-EAI infected | -0.0908 | 0.0422 | 0.0664 |

| Par | Estimate | se | OR | pvalue |
|---|---|---|---|---|
| (Intercept) | -3.484 | 0.6133 | 0.0307 | 0 |
| groupAfri not infected | 0.9109 | 0.6428 | 2.4866 | 0.1565 |
| groupBCG not infected | 1.7782 | 0.5663 | 5.9191 | 0.0017 |
| groupBeijing not infected | 0.5988 | 0.7014 | 1.82 | 0.3933 |
| groupCDC1551 not infected | -0.2013 | 1.1277 | 0.8176 | 0.8583 |
| groupEAI not infected | 1.3244 | 0.5316 | 3.7599 | 0.0127 |
| groupH37Ra not infected | 1.4107 | 0.5892 | 4.0989 | 0.0166 |
| groupH37Rv not infected | 1.1886 | 0.5907 | 3.2824 | 0.0442 |
| groupAfri infected | 1.9846 | 0.5506 | 7.2758 | 3e-04 |
| groupBCG infected | 2.805 | 0.5607 | 16.5272 | 0 |
| groupBeijing infected | 1.4598 | 0.5585 | 4.3052 | 0.009 |
| groupCDC1551 infected | 1.3262 | 0.7207 | 3.7668 | 0.0657 |
| groupEAI infected | 1.951 | 0.5188 | 7.0354 | 2e-04 |
| groupH37Ra infected | 2.3266 | 0.5764 | 10.243 | 1e-04 |
| groupH37Rv infected | 1.7108 | 0.5785 | 5.5335 | 0.0031 |

| Contrast | estimate | SE | p.value |
|---|---|---|---|
| none not infected-Afri not infected | 2.5731 | 0.6172 | 0.001 |
| none not infected-BCG not infected | 1.7058 | 0.5856 | 0.0189 |
| none not infected-Beijing not infected | 2.8851 | 0.6739 | 0.001 |
| none not infected-CDC1551 not infected | 3.6853 | 1.1059 | 0.0077 |
| none not infected-EAI not infected | 2.1596 | 0.5412 | 0.0011 |
| none not infected-H37Ra not infected | 2.0732 | 0.5744 | 0.0032 |
| none not infected-H37Rv not infected | 2.2954 | 0.5766 | 0.0011 |
| none not infected-Afri infected | 1.4994 | 0.5203 | 0.0192 |
| none not infected-BCG infected | 0.6789 | 0.5802 | 0.3944 |
| none not infected-Beijing infected | 2.0241 | 0.5265 | 0.0015 |
| none not infected-CDC1551 infected | 2.1577 | 0.6859 | 0.0116 |
| none not infected-EAI infected | 1.533 | 0.5264 | 0.0189 |
| none not infected-H37Ra infected | 1.1574 | 0.5541 | 0.1178 |
| none not infected-H37Rv infected | 1.7731 | 0.56 | 0.0116 |
| Afri not infected-BCG not infected | -0.8673 | 0.5955 | 0.2996 |
| Afri not infected-Beijing not infected | 0.3121 | 0.6701 | 0.7089 |
| Afri not infected-CDC1551 not infected | 1.1122 | 1.1161 | 0.4785 |
| Afri not infected-EAI not infected | -0.4135 | 0.543 | 0.5859 |
| Afri not infected-H37Ra not infected | -0.4998 | 0.6498 | 0.5859 |
| Afri not infected-H37Rv not infected | -0.2777 | 0.6554 | 0.7174 |
| BCG not infected-Beijing not infected | 1.1794 | 0.6683 | 0.2131 |
| BCG not infected-CDC1551 not infected | 1.9795 | 1.1224 | 0.2131 |
| BCG not infected-EAI not infected | 0.4538 | 0.3524 | 0.3666 |
| BCG not infected-H37Ra not infected | 0.3675 | 0.6323 | 0.6653 |
| BCG not infected-H37Rv not infected | 0.5896 | 0.6358 | 0.5065 |
| Beijing not infected-CDC1551 not infected | 0.8001 | 1.1422 | 0.6218 |
| Beijing not infected-EAI not infected | -0.7256 | 0.6229 | 0.3944 |
| Beijing not infected-H37Ra not infected | -0.8119 | 0.6971 | 0.3944 |
| Beijing not infected-H37Rv not infected | -0.5898 | 0.701 | 0.5602 |
| CDC1551 not infected-EAI not infected | -1.5257 | 1.0974 | 0.3237 |
| CDC1551 not infected-H37Ra not infected | -1.6121 | 1.1087 | 0.2996 |
| CDC1551 not infected-H37Rv not infected | -1.3899 | 1.1093 | 0.3679 |
| EAI not infected-H37Ra not infected | -0.0863 | 0.5905 | 0.898 |
| EAI not infected-H37Rv not infected | 0.1358 | 0.5944 | 0.8569 |
| H37Ra not infected-H37Rv not infected | 0.2221 | 0.3913 | 0.6653 |
| Afri infected-BCG infected | -0.8205 | 0.4894 | 0.2459 |
| Afri infected-Beijing infected | 0.5247 | 0.4148 | 0.3679 |
| Afri infected-CDC1551 infected | 0.6583 | 0.6258 | 0.45 |
| Afri infected-EAI infected | 0.0336 | 0.4105 | 0.9348 |
| Afri infected-H37Ra infected | -0.342 | 0.5317 | 0.6424 |
| Afri infected-H37Rv infected | 0.2737 | 0.5451 | 0.7051 |
| BCG infected-Beijing infected | 1.3452 | 0.5093 | 0.0372 |
| BCG infected-CDC1551 infected | 1.4788 | 0.7104 | 0.1178 |
| BCG infected-EAI infected | 0.8541 | 0.3319 | 0.0423 |
| BCG infected-H37Ra infected | 0.4784 | 0.6097 | 0.5859 |
| BCG infected-H37Rv infected | 1.0942 | 0.6158 | 0.2131 |
| Beijing infected-CDC1551 infected | 0.1336 | 0.621 | 0.8569 |
| Beijing infected-EAI infected | -0.4911 | 0.4352 | 0.4082 |

**M2 – 24 h p.i.**

c) Effective number of mycobacteria per cell (M/C)

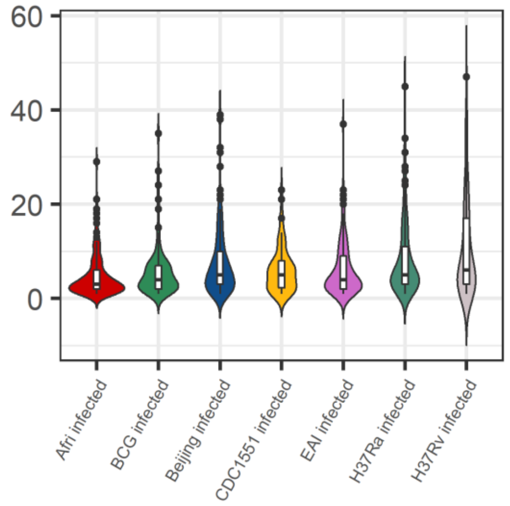


| Par | Value | Std.Error | p.value |
|---|---|---|---|
| (Intercept) | 1.126 | 0.1674 | 0 |
| groupBCG infected | 0.7435 | 0.235 | 0.0025 |
| groupBeijing infected | 0.3785 | 0.1311 | 0.0055 |
| groupCDC1551 infected | 0.1261 | 0.1629 | 0.4418 |
| groupEAI infected | 0.6265 | 0.1691 | 5e-04 |
| groupH37Ra infected | 0.6752 | 0.186 | 6e-04 |
| groupH37Rv infected | 0.7453 | 0.1955 | 3e-04 |

| Contrast | estimate | SE | p.value |
|---|---|---|---|
| Afri infected - BCG infected | -0.7435 | 0.235 | 0.0104 |
| Afri infected - Beijing infected | -0.3785 | 0.1311 | 0.0164 |
| Afri infected - CDC1551 infected | -0.1261 | 0.1629 | 0.6185 |
| Afri infected - EAI infected | -0.6265 | 0.1691 | 0.0042 |
| Afri infected - H37Ra infected | -0.6752 | 0.186 | 0.0042 |
| Afri infected - H37Rv infected | -0.7453 | 0.1955 | 0.0042 |
| BCG infected - Beijing infected | 0.365 | 0.2331 | 0.1984 |
| BCG infected - CDC1551 infected | 0.6174 | 0.2601 | 0.0488 |
| BCG infected - EAI infected | 0.117 | 0.1746 | 0.6634 |
| BCG infected - H37Ra infected | 0.0684 | 0.271 | 0.8669 |
| BCG infected - H37Rv infected | -0.0018 | 0.2748 | 0.9949 |
| Beijing infected - CDC1551 infected | 0.2524 | 0.1509 | 0.1744 |
| Beijing infected - EAI infected | -0.248 | 0.1672 | 0.215 |
| Beijing infected - H37Ra infected | -0.2966 | 0.1765 | 0.1744 |
| Beijing infected - H37Rv infected | -0.3667 | 0.18 | 0.0968 |
| CDC1551 infected - EAI infected | -0.5004 | 0.2047 | 0.0461 |
| CDC1551 infected - H37Ra infected | -0.549 | 0.1879 | 0.0164 |
| CDC1551 infected - H37Rv infected | -0.6192 | 0.1915 | 0.0104 |
| EAI infected - H37Ra infected | -0.0486 | 0.2197 | 0.8669 |
| EAI infected - H37Rv infected | -0.1187 | 0.2246 | 0.7399 |
| H37Ra infected - H37Rv infected | -0.0701 | 0.1606 | 0.7748 |

d) Colocalization Index

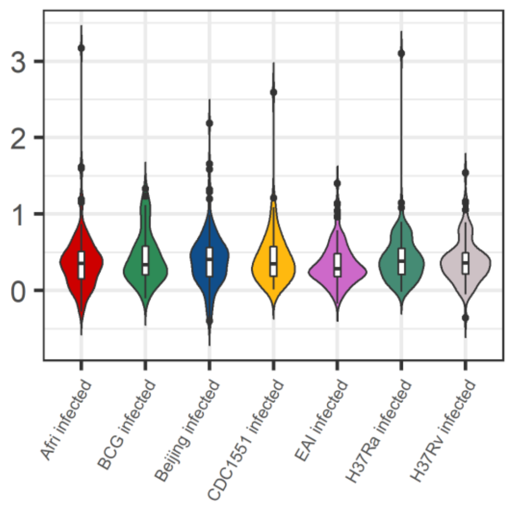


| Par | Value | Std.Error | p.value |
|---|---|---|---|
| (Intercept) | -0.1201 | 0.0896 | 0.1804 |
| groupBCG infected | 0.1305 | 0.1528 | 0.3968 |
| groupBeijing infected | 0.1024 | 0.1093 | 0.3527 |
| groupCDC1551 infected | 0.1207 | 0.1284 | 0.3511 |
| groupEAI infected | -0.1137 | 0.119 | 0.3433 |
| groupH37Ra infected | 0.121 | 0.1261 | 0.3414 |
| groupH37Rv infected | 0.0021 | 0.1336 | 0.9873 |

| Contrast | estimate | SE | p.value |
|---|---|---|---|
| Afri infected - BCG infected | -0.1305 | 0.1528 | 0.6543 |
| Afri infected - Beijing infected | -0.1024 | 0.1093 | 0.6543 |
| Afri infected - CDC1551 infected | -0.1207 | 0.1284 | 0.6543 |
| Afri infected - EAI infected | 0.1137 | 0.119 | 0.6543 |
| Afri infected - H37Ra infected | -0.121 | 0.1261 | 0.6543 |
| Afri infected - H37Rv infected | -0.0021 | 0.1336 | 0.9985 |
| BCG infected - Beijing infected | 0.0281 | 0.1498 | 0.9985 |
| BCG infected - CDC1551 infected | 0.0098 | 0.1651 | 0.9985 |
| BCG infected - EAI infected | 0.2442 | 0.1298 | 0.4907 |
| BCG infected - H37Ra infected | 0.0095 | 0.1599 | 0.9985 |
| BCG infected - H37Rv infected | 0.1283 | 0.1637 | 0.6543 |
| Beijing infected - CDC1551 infected | -0.0183 | 0.1235 | 0.9985 |
| Beijing infected - EAI infected | 0.2161 | 0.1154 | 0.4907 |
| Beijing infected - H37Ra infected | -0.0185 | 0.1226 | 0.9985 |
| Beijing infected - H37Rv infected | 0.1003 | 0.1274 | 0.6543 |
| CDC1551 infected - EAI infected | 0.2344 | 0.1374 | 0.4907 |
| CDC1551 infected - H37Ra infected | -3e-04 | 0.1352 | 0.9985 |
| CDC1551 infected - H37Rv infected | 0.1186 | 0.1404 | 0.6543 |
| EAI infected - H37Ra infected | -0.2346 | 0.1318 | 0.4907 |
| EAI infected - H37Rv infected | -0.1158 | 0.1365 | 0.6543 |
| H37Ra infected - H37Rv infected | 0.1188 | 0.1235 | 0.6543 |

**ANALYSIS ON THE NUMBER OF LYSOSENSOR-POSITIVE COMPARTMENTS**

**GEO_HOST_adaptation, M1-day=1**

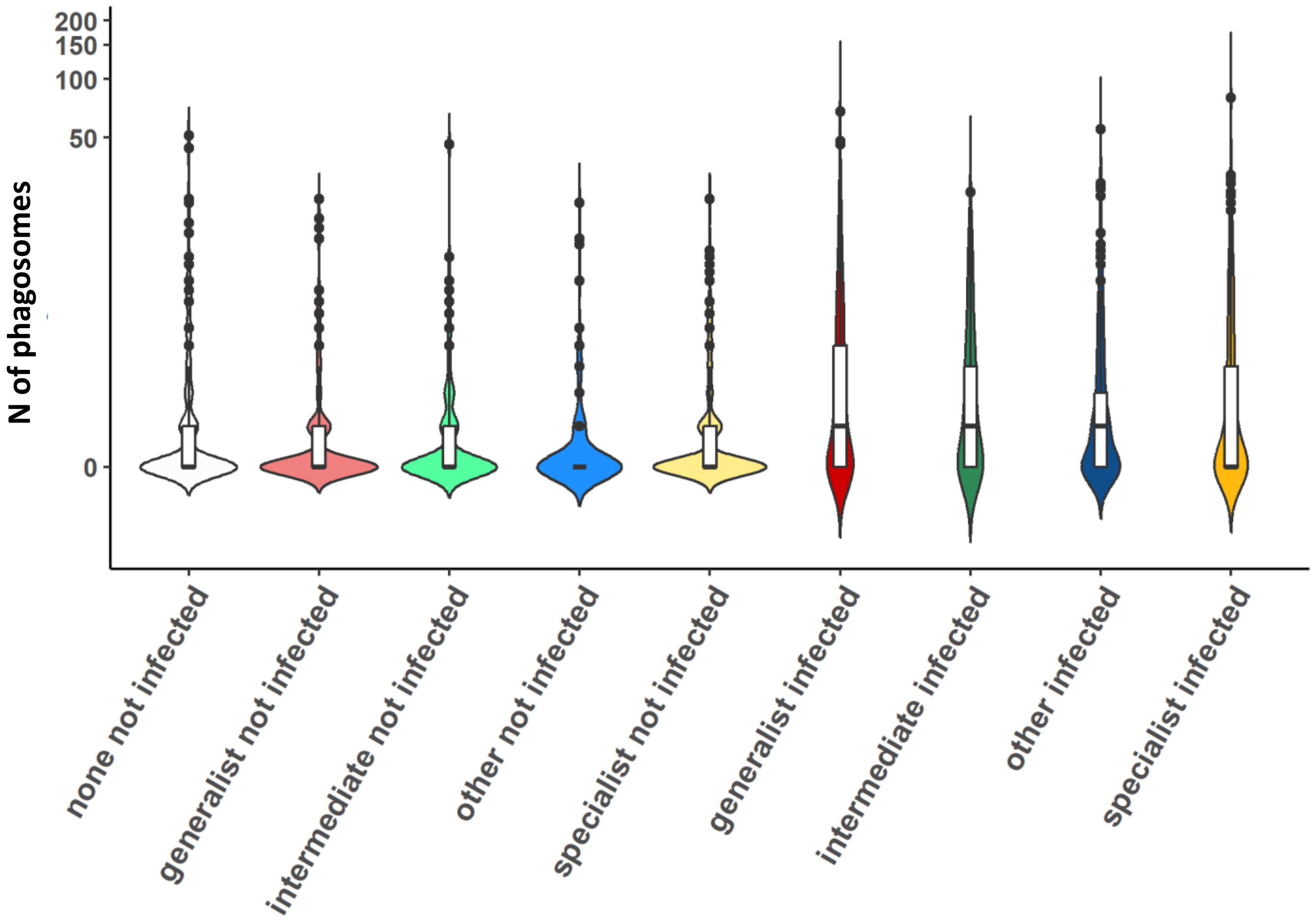

Zero-inflated Mixed-effects Poisson model
Random effects were defined nested with respect to EXP/well

Conditional model

| | **Estimate** | **SE** | **p-value** |
|---|---|---|---|
| Intercept | 0.836 | 0.276 | 0.0025 |
| groupgeneralist not infected | -0.252 | 0.2359 | 0.2855 |
| groupintermediate not infected | -0.1189 | 0.3026 | 0.6944 |
| groupother not infected | -0.1857 | 0.2745 | 0.4988 |
| groupspecialist not infected | 0.1435 | 0.2552 | 0.5739 |
| groupgeneralist infected | 0.3905 | 0.2296 | 0.0889 |
| groupintermediate infected | 0.2791 | 0.293 | 0.3408 |
| groupother infected | 0.1793 | 0.262 | 0.4938 |
| groupspecialist infected | 0.9579 | 0.2492 | 0.0001 |

Zero-inflation model

| | **Estimate** | **SE** | **p-value** |
|---|---|---|---|
| Intercept | 0.623 | 0.3462 | 0.0719 |
| groupgeneralist not infected | 0.388 | 0.3098 | 0.2104 |
| groupintermediate not infected | 0.4105 | 0.3952 | 0.2989 |
| groupother not infected | 0.9999 | 0.3517 | 0.0045 |
| groupspecialist not infected | 0.0701 | 0.3225 | 0.8279 |
| groupgeneralist infected | -1.0992 | 0.3073 | 3e-04 |
| groupintermediate infected | -0.7924 | 0.3911 | 0.0428 |
| groupother infected | -0.5407 | 0.34 | 0.1118 |
| groupspecialist infected | -0.7732 | 0.3181 | 0.0151 |

| **Contrast** | **estimate** | **SE** | **p-value*** |
|---|---|---|---|
| none not infected-generalist not infected | 0.252 | 0.2359 | 0.5709 |
| none not infected-intermediate not infected | 0.1189 | 0.3026 | 0.7936 |
| none not infected-other not infected | 0.1857 | 0.2745 | 0.7041 |
| none not infected-specialist not infected | -0.1435 | 0.2552 | 0.7652 |
| none not infected-generalist infected | -0.3905 | 0.2296 | 0.2135 |
| none not infected-intermediate infected | -0.2791 | 0.293 | 0.6162 |
| none not infected-other infected | -0.1793 | 0.262 | 0.7041 |
| **none not infected-specialist infected** | **-0.9579** | **0.2492** | **0.001** |
| generalist not infected-intermediate not infected | -0.1331 | 0.28 | 0.7936 |
| generalist not infected-other not infected | -0.0663 | 0.2621 | 0.8352 |
| generalist not infected-specialist not infected | -0.3955 | 0.2309 | 0.2135 |
| intermediate not infected-other not infected | 0.0668 | 0.3263 | 0.8377 |
| intermediate not infected-specialist not infected | -0.2624 | 0.3007 | 0.6162 |
| other not infected-specialist not infected | -0.3292 | 0.2611 | 0.4526 |
| generalist infected-intermediate infected | 0.1114 | 0.2636 | 0.7936 |
| generalist infected-other infected | 0.2112 | 0.2432 | 0.6162 |
| **generalist infected-specialist infected** | **-0.5674** | **0.2175** | **0.0312** |
| intermediate infected-other infected | 0.0998 | 0.3064 | 0.8124 |
| intermediate infected-specialist infected | -0.6788 | 0.285 | 0.0517 |
| **other infected-specialist infected** | **-0.7786** | **0.2412** | **0.005** |
| **generalist not infected-generalist infected** | **-0.6425** | **0.07** | **<0.0001** |
| **intermediate not infected-intermediate infected** | **-0.398** | **0.1062** | **0.0011** |
| **other not infected-other infected** | **-0.365** | **0.1108** | **0.0047** |
| **specialist not infected-specialist infected** | **-0.8144** | **0.0745** | **<0.0001** |

* False discovery rate (FDR) correction

GEO_HOST_adaptation, M1-day=2

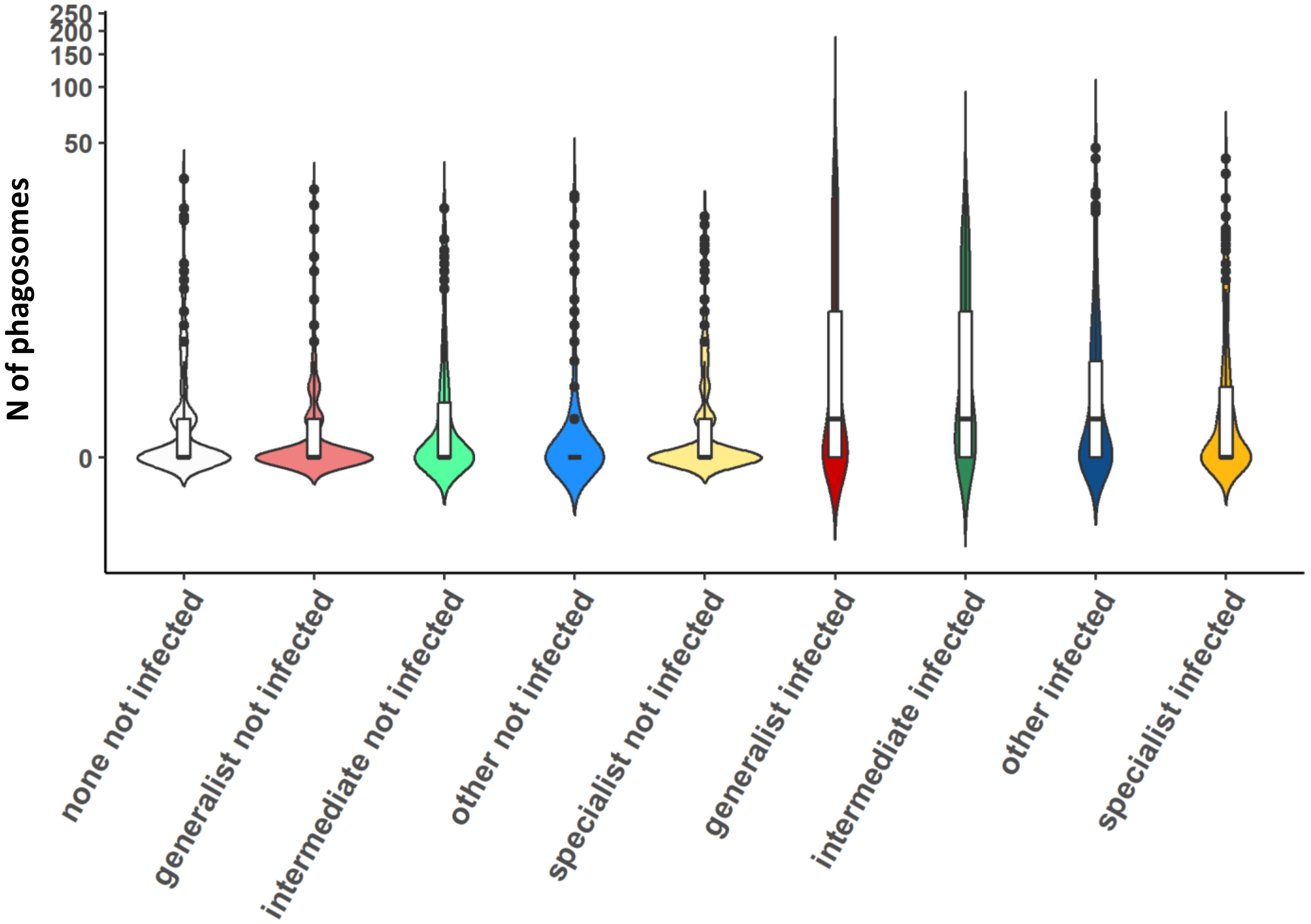

Zero-inflated Mixed-effects Poisson model
Random effects were defined nested with respect to EXP/well

Conditional model

| | Estimate | SE | p-value |
|---|---|---|---|
| Intercept | 0.7967 | 0.2426 | 0.001 |
| groupgeneralist not infected | -0.024 | 0.2663 | 0.928 |
| groupintermediate not infected | 0.5369 | 0.3227 | 0.0961 |
| groupother not infected | 0.4789 | 0.2945 | 0.104 |
| groupspecialist not infected | 0.3259 | 0.2741 | 0.2345 |
| groupgeneralist infected | 1.2219 | 0.2559 | <0.0001 |
| groupintermediate infected | 0.8383 | 0.3151 | 0.0078 |
| groupother infected | 0.8774 | 0.2848 | 0.0021 |
| groupspecialist infected | 0.5818 | 0.2715 | 0.0321 |

Zero-inflation model

| | Estimate | SE | p-value |
|---|---|---|---|
| Intercept | 0.3109 | 0.2958 | 0.2933 |
| groupgeneralist not infected | 0.4307 | 0.307 | 0.1606 |
| groupintermediate not infected | 0.4463 | 0.3747 | 0.2335 |
| groupother not infected | 0.9704 | 0.3463 | 0.0051 |
| groupspecialist not infected | 0.2976 | 0.2986 | 0.3189 |
| groupgeneralist infected | -0.7279 | 0.2837 | 0.0103 |
| groupintermediate infected | -1.0123 | 0.3718 | 0.0065 |
| groupother infected | -0.3559 | 0.3117 | 0.2535 |
| groupspecialist infected | -0.3045 | 0.2951 | 0.3021 |

| **Contrast** | **estimate** | **SE** | **p-value*** |
|---|---|---|---|
| none not infected-generalist not infected | 0.024 | 0.2663 | 0.928 |
| none not infected-intermediate not infected | -0.5369 | 0.3227 | 0.1919 |
| none not infected-other not infected | -0.4789 | 0.2945 | 0.1919 |
| none not infected-specialist not infected | -0.3259 | 0.2741 | 0.3142 |
| **none not infected-generalist infected** | **-1.2219** | **0.2559** | **<0.0001** |
| **none not infected-intermediate infected** | **-0.8383** | **0.3151** | **0.0234** |
| **none not infected-other infected** | **-0.8774** | **0.2848** | **0.0099** |
| none not infected-specialist infected | -0.5818 | 0.2715 | 0.0856 |
| generalist not infected-intermediate not infected | -0.5609 | 0.2985 | 0.133 |
| generalist not infected-other not infected | -0.5029 | 0.2684 | 0.133 |
| generalist not infected-specialist not infected | -0.35 | 0.2356 | 0.2357 |
| intermediate not infected-other not infected | 0.058 | 0.3358 | 0.928 |
| intermediate not infected-specialist not infected | 0.211 | 0.3121 | 0.5989 |
| other not infected-specialist not infected | 0.1529 | 0.2635 | 0.6419 |
| generalist infected-intermediate infected | 0.3836 | 0.2806 | 0.2575 |
| generalist infected-other infected | 0.3445 | 0.2456 | 0.2573 |
| **generalist infected-specialist infected** | **0.6401** | **0.2198** | **0.0144** |
| intermediate infected-other infected | -0.0391 | 0.3197 | 0.928 |
| intermediate infected-specialist infected | 0.2565 | 0.3022 | 0.5002 |
| other infected-specialist infected | 0.2956 | 0.2493 | 0.3142 |
| **generalist not infected-generalist infected** | **-1.2459** | **0.0847** | **<0.0001** |
| **intermediate not infected-intermediate infected** | **-0.3014** | **0.1102** | **0.0213** |
| **other not infected-other infected** | **-0.3986** | **0.0943** | **0.0002** |
| **specialist not infected-specialist infected** | **-0.2559** | **0.0736** | **0.003** |

* False discovery rate (FDR) correction

GEO_HOST_adaptation, M2-day=1

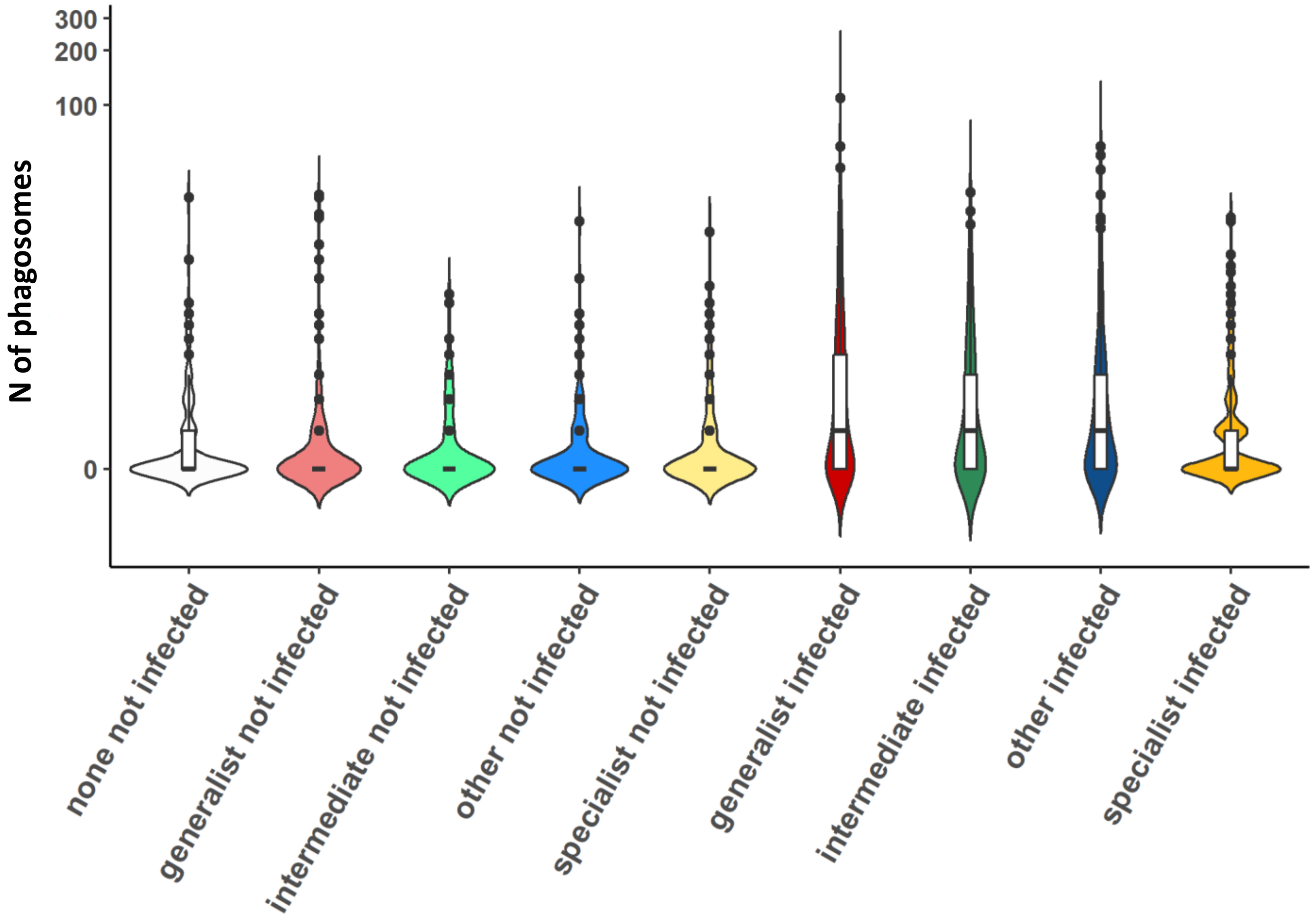

Zero-inflated Mixed-effects Poisson model
Random effects were defined nested with respect to EXP/well

Conditional model

| | Estimate | SE | p-value |
|---|---|---|---|
| Intercept | 0.5071 | 0.2957 | 0.0863 |
| groupgeneralist not infected | 0.2814 | 0.2841 | 0.3219 |
| groupintermediate not infected | -0.3785 | 0.3625 | 0.2963 |
| groupother not infected | -0.1204 | 0.3034 | 0.6914 |
| groupspecialist not infected | -0.0118 | 0.3037 | 0.969 |
| groupgeneralist infected | 0.8031 | 0.2782 | 0.0039 |
| groupintermediate infected | 0.688 | 0.3403 | 0.0432 |
| groupother infected | 0.8053 | 0.2886 | 0.0053 |
| groupspecialist infected | 0.3975 | 0.2975 | 0.1815 |

Zero-inflation model

| | Estimate | SE | p-value |
|---|---|---|---|
| Intercept | 0.88 | 0.3608 | 0.0147 |
| groupgeneralist not infected | 0.3193 | 0.2765 | 0.2482 |
| groupintermediate not infected | 0.4372 | 0.348 | 0.209 |
| groupother not infected | 0.2471 | 0.2982 | 0.4074 |
| groupspecialist not infected | -0.0445 | 0.2904 | 0.8783 |
| groupgeneralist infected | -1.2474 | 0.2695 | <0.0001 |
| groupintermediate infected | -0.7226 | 0.3164 | 0.0224 |
| groupother infected | -1.0517 | 0.2833 | 0.0002 |
| groupspecialist infected | -0.9201 | 0.2875 | 0.0014 |

| Contrast | estimate | SE | p-value* |
|---|---|---|---|
| none not infected-generalist not infected | -0.2814 | 0.2841 | 0.4829 |
| none not infected-intermediate not infected | 0.3785 | 0.3625 | 0.4741 |
| none not infected-other not infected | 0.1204 | 0.3034 | 0.78 |
| none not infected-specialist not infected | 0.0118 | 0.3037 | 0.9932 |
| **none not infected-generalist infected** | **-0.8031** | **0.2782** | **0.0187** |
| none not infected-intermediate infected | -0.688 | 0.3403 | 0.1296 |
| **none not infected-other infected** | **-0.8053** | **0.2886** | **0.0211** |
| none not infected-specialist infected | -0.3975 | 0.2975 | 0.3629 |
| generalist not infected-intermediate not infected | 0.66 | 0.3059 | 0.1062 |
| generalist not infected-other not infected | 0.4018 | 0.2783 | 0.3246 |
| generalist not infected-specialist not infected | 0.2932 | 0.2522 | 0.4524 |
| intermediate not infected-other not infected | -0.2581 | 0.3579 | 0.6278 |
| intermediate not infected-specialist not infected | -0.3667 | 0.3371 | 0.4741 |
| other not infected-specialist not infected | -0.1086 | 0.277 | 0.78 |
| generalist infected-intermediate infected | 0.1151 | 0.273 | 0.78 |
| generalist infected-other infected | -0.0022 | 0.2555 | 0.9932 |
| generalist infected-specialist infected | 0.4056 | 0.2378 | 0.2347 |
| intermediate infected-other infected | -0.1173 | 0.3213 | 0.78 |
| intermediate infected-specialist infected | 0.2905 | 0.306 | 0.4834 |
| other infected-specialist infected | 0.4078 | 0.2529 | 0.2564 |
| **generalist not infected-generalist infected** | **-0.5217** | **0.0701** | **<0.0001** |
| **intermediate not infected-intermediate infected** | **-1.0665** | **0.1467** | **<0.0001** |
| **other not infected-other infected** | **-0.9257** | **0.1112** | **<0.0001** |
| **specialist not infected-specialist infected** | **-0.4093** | **0.0948** | **0.0001** |

* False discovery rate (FDR) correction

**GEO_HOST_adaptation, M2-day=2**

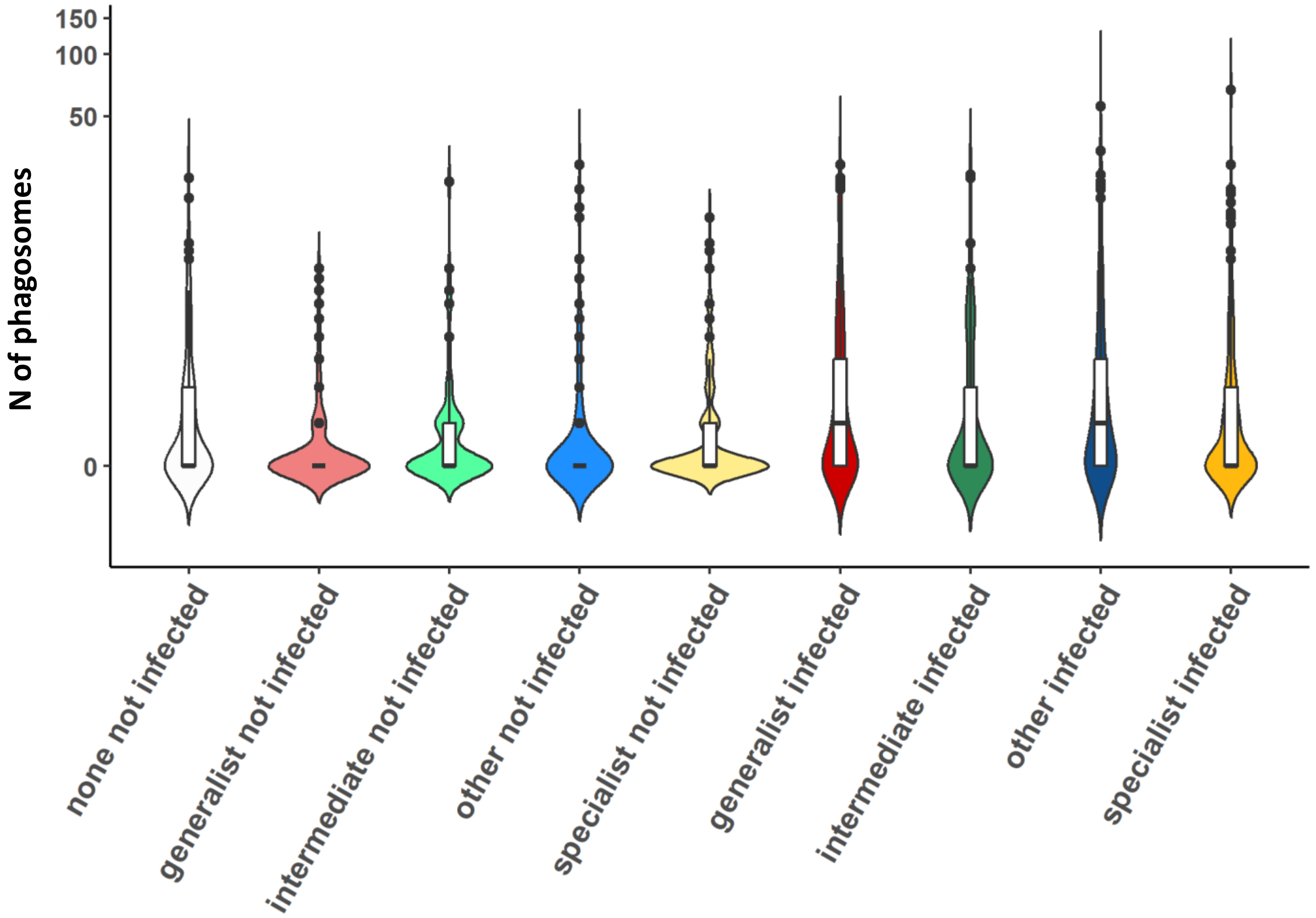

Zero-inflated Mixed-effects Poisson model
Random effects were defined nested with respect to EXP/well

Conditional model

| | **Estimate** | **SE** | **p-value** |
|---|---|---|---|
| Intercept | 1.0011 | 0.2105 | <0.0001 |
| groupgeneralist not infected | -0.4274 | 0.2408 | 0.0759 |
| groupintermediate not infected | -0.3705 | 0.293 | 0.2061 |
| groupother not infected | 0.1869 | 0.2657 | 0.4817 |
| groupspecialist not infected | -0.2388 | 0.2496 | 0.3386 |
| groupgeneralist infected | 0.3822 | 0.2184 | 0.0801 |
| groupintermediate infected | 0.3787 | 0.2784 | 0.1738 |
| groupother infected | 0.358 | 0.256 | 0.162 |
| groupspecialist infected | 0.2613 | 0.2413 | 0.2789 |

Zero-inflation model

| | **Estimate** | **SE** | **p-value** |
|---|---|---|---|
| Intercept | 0.3905 | 0.3332 | 0.2411 |
| groupgeneralist not infected | 0.577 | 0.2816 | 0.0404 |
| groupintermediate not infected | 0.2645 | 0.3391 | 0.4354 |
| groupother not infected | 0.6891 | 0.295 | 0.0195 |
| groupspecialist not infected | 0.2852 | 0.2667 | 0.2849 |
| groupgeneralist infected | -0.6093 | 0.2499 | 0.0148 |
| groupintermediate infected | 0.3193 | 0.3171 | 0.3139 |
| groupother infected | -0.8391 | 0.2757 | 0.0023 |
| groupspecialist infected | -0.2514 | 0.2539 | 0.3222 |

| **Contrast** | **estimate** | **SE** | **p-value** |
|---|---|---|---|
| none not infected-generalist not infected | 0.4274 | 0.2408 | 0.2137 |
| none not infected-intermediate not infected | 0.3705 | 0.293 | 0.4122 |
| none not infected-other not infected | -0.1869 | 0.2657 | 0.7226 |
| none not infected-specialist not infected | 0.2388 | 0.2496 | 0.5805 |
| none not infected-generalist infected | -0.3822 | 0.2184 | 0.2137 |
| none not infected-intermediate infected | -0.3787 | 0.2784 | 0.3791 |
| none not infected-other infected | -0.358 | 0.256 | 0.3791 |
| none not infected-specialist infected | -0.2613 | 0.2413 | 0.5149 |
| generalist not infected-intermediate not infected | -0.0569 | 0.2881 | 0.9639 |
| generalist not infected-other not infected | -0.6143 | 0.2555 | 0.0972 |
| generalist not infected-specialist not infected | -0.1885 | 0.2386 | 0.6871 |
| intermediate not infected-other not infected | -0.5574 | 0.3102 | 0.2137 |
| intermediate not infected-specialist not infected | -0.1316 | 0.2999 | 0.808 |
| other not infected-specialist not infected | 0.4258 | 0.2426 | 0.2137 |
| generalist infected-intermediate infected | 0.0035 | 0.2536 | 0.9891 |
| generalist infected-other infected | 0.0242 | 0.223 | 0.9834 |
| generalist infected-specialist infected | 0.1209 | 0.2061 | 0.787 |
| intermediate infected-other infected | 0.0208 | 0.2875 | 0.9834 |
| intermediate infected-specialist infected | 0.1174 | 0.2786 | 0.808 |
| other infected-specialist infected | 0.0967 | 0.2242 | 0.808 |
| **generalist not infected-generalist infected** | **-0.8096** | **0.1212** | **<0.0001** |
| **intermediate not infected-intermediate infected** | **-0.7492** | **0.1509** | **<0.0001** |
| other not infected-other infected | -0.171 | 0.0921 | 0.2137 |
| **specialist not infected-specialist infected** | **-0.5001** | **0.0905** | **<0.0001** |

* False discovery rate (FDR) correction
