## Supplementary material for "*Mycobacterium tuberculosis* diversity and macrophage heterogeneity dictate phagosomal acidification": Supplmenetary 3

### SUPPLEMENTARY 3

***Number of mycobacteria internalized per cell (M/C) over time***

Comparison of M/C observed between time1 (6h p.i.) and time2 (24h p.i.)

GEO HOST ADAPTATION, M1

| Par | Value | Std.Error | p.value |
|---|---|---|---|
| (Intercept) | -0.1346 | 0.1234 | 0.2754 |
| groupintermediate infected | -0.0708 | 0.1359 | 0.6043 |
| groupother infected | 0.1237 | 0.1239 | 0.3219 |
| groupspecialist infected | -0.0134 | 0.1083 | 0.9021 |
| time2 | 0.1794 | 0.0728 | 0.0139 |
| groupintermediate infected:time2 | 0.2195 | 0.1423 | 0.1232 |
| groupother infected:time2 | 0.3517 | 0.1179 | 0.0029 |
| groupspecialist infected:time2 | -0.201 | 0.1048 | 0.0553 |

| **contrast** | **group** | **estimate** | **SE** | **p.value** |
|---|---|---|---|---|
| time1 - time2 | generalist infected | -0.1794 | 0.0728 | 0.0139 |
| time1 - time2 | intermediate infected | -0.3989 | 0.1222 | 0.0011 |
| time1 - time2 | other infected | -0.5312 | 0.0931 | <0.0001 |
| time1 - time2 | specialist infected | 0.0216 | 0.0753 | 0.7748 |

GEO HOST ADAPTATION, M2

| Par | Value | Std.Error | p.value |
|---|---|---|---|
| (Intercept) | -0.0873 | 0.1289 | 0.4985 |
| groupintermediate infected | -0.0647 | 0.1302 | 0.6209 |
| groupother infected | 0.0188 | 0.117 | 0.8728 |
| groupspecialist infected | 0.0851 | 0.1044 | 0.4181 |
| time2 | 0.2436 | 0.0739 | 0.001 |
| groupintermediate infected:time2 | -0.2991 | 0.1456 | 0.0401 |
| groupother infected:time2 | -0.0636 | 0.1157 | 0.5824 |
| groupspecialist infected:time2 | -0.3512 | 0.1051 | 0.0008 |

| **contrast** | **group** | **estimate** | **SE** | **p.value** |
|---|---|---|---|---|
| time1 - time2 | generalist infected | -0.2436 | 0.0739 | 0.001 |
| time1 - time2 | intermediate infected | 0.0555 | 0.1256 | 0.6585 |
| time1 - time2 | other infected | -0.18 | 0.0894 | 0.0442 |
| time1 - time2 | specialist infected | 0.1076 | 0.0747 | 0.1501 |

#### COMPARISON M1 vs M2

#### Acidification Mean

Statistical method 1: Linear quantile mixed models (LQMM) on the median were applied to test the difference between M1 and M2, accounting for group membership.

Time 1

| | Value | Std. Error | p-value |
|---|---|---|---|
| Intercept | 0.8125 | 0.1582 | <0.0001 |
| groupgeneralist not infected | 0.0737 | 0.0960 | 0.4461 |
| groupintermediate not infected | 0.1759 | 0.0846 | 0.0428 |
| groupother not infected | -0.1229 | 0.1001 | 0.2258 |
| groupspecialist not infected | 0.2224 | 0.0858 | 0.0126 |
| groupgeneralist infected | 0.2114 | 0.1092 | 0.0586 |
| groupintermediate infected | 0.3325 | 0.0956 | 0.0011 |
| groupother infected | -0.0058 | 0.1000 | 0.9539 |
| groupspecialist infected | 0.3908 | 0.0842 | 0.0000 |
| M2 | 0.2244 | 0.1469 | 0.1330 |

M2 values are not significantly different from the M1 values (p=0.133).

Time 2

| | Value | Std. Error | p-value |
|---|---|---|---|
| Intercept | 1.0034 | 0.0266 | <0.0001 |
| groupgeneralist not infected | -0.0037 | 0.0357 | 0.9188 |
| groupintermediate not infected | -0.0771 | 0.0394 | 0.0561 |
| groupother not infected | 0.0646 | 0.0541 | 0.2384 |
| groupspecialist not infected | 0.1043 | 0.0266 | 0.0003 |
| groupgeneralist infected | 0.1422 | 0.0405 | 0.0010 |
| groupintermediate infected | 0.0865 | 0.0551 | 0.1230 |
| groupother infected | 0.1805 | 0.0558 | 0.0022 |
| groupspecialist infected | 0.2150 | 0.0298 | 0.0000 |
| M2 | -0.0109 | 0.0359 | 0.7627 |

M2 values are not significantly different from the M1 values (p=0.7627).

Statistical method 2: Linear quantile mixed models (LQMM) on the median were applied to assess the difference between M1 and M2 within each group.

Time 1

| | Value | Std. Error | p-value |
|---|---|---|---|
| Intercept | 0.9152 | 0.1184 | <0.0001 |
| groupgeneralist not infected | -0.0003 | 0.1209 | 0.9978 |
| groupintermediate not infected | -0.1275 | 0.1129 | 0.2642 |
| groupother not infected | -0.1929 | 0.1375 | 0.1668 |
| groupspecialist not infected | 0.2745 | 0.1447 | 0.0637 |
| groupgeneralist infected | 0.1042 | 0.1349 | 0.4438 |

| | | | |
|---|---|---|---|
| groupintermediate infected | 0.0231 | 0.1188 | 0.8469 |
| groupother infected | -0.1062 | 0.1329 | 0.4278 |
| groupspecialist infected | 0.4701 | 0.1537 | 0.0036 |
| M2 | 0.1071 | 0.1079 | 0.3261 |
| groupgeneralist not infected:M2 | 0.2334 | 0.1656 | 0.1650 |
| groupintermediate not infected:M2 | 0.3555 | 0.1096 | 0.0021 |
| groupother not infected:M2 | 0.1829 | 0.1429 | 0.2066 |
| groupspecialist not infected:M2 | -0.0619 | 0.1295 | 0.6350 |
| groupgeneralist infected:M2 | 0.3084 | 0.1962 | 0.1225 |
| groupintermediate infected:M2 | 0.3697 | 0.1272 | 0.0055 |
| groupother infected:M2 | 0.2560 | 0.1530 | 0.1007 |
| groupspecialist infected:M2 | -0.1236 | 0.1398 | 0.3811 |

M2 values were significantly different from M1 values in groups intermediate not infected and intermediate infected, while no significant differences were observed in the other groups.

Time 2

| | Value | Std. Error | p-value |
|---|---|---|---|
| (Intercept) | 0.9919 | 0.0242 | < 0.0001 |
| groupgeneralist not infected | 0.0119 | 0.0433 | 0.7848 |
| groupintermediate not infected | 0.0221 | 0.0520 | 0.6730 |
| groupother not infected | -0.0653 | 0.0517 | 0.2128 |
| groupspecialist not infected | 0.1401 | 0.0415 | 0.0015 |
| groupgeneralist infected | 0.1766 | 0.0658 | 0.0099 |
| groupintermediate infected | 0.2892 | 0.0729 | 0.0002 |
| groupother infected | 0.0730 | 0.0588 | 0.2207 |
| groupspecialist infected | 0.2641 | 0.0444 | 0.0000 |
| M2 | 0.0106 | 0.0332 | 0.7518 |
| groupgeneralist not infected:M2 | -0.0384 | 0.0639 | 0.5507 |
| groupintermediate not infected:M2 | -0.1111 | 0.0730 | 0.1341 |
| groupother not infected:M2 | 0.1677 | 0.0881 | 0.0629 |
| groupspecialist not infected:M2 | -0.0465 | 0.0663 | 0.4863 |
| groupgeneralist infected:M2 | -0.1028 | 0.0792 | 0.2005 |
| groupintermediate infected:M2 | -0.3102 | 0.0794 | 0.0003 |
| groupother infected:M2 | 0.1416 | 0.1067 | 0.1906 |
| groupspecialist infected:M2 | -0.0552 | 0.0677 | 0.4191 |

M2 values were significantly different from M1 values in groups intermediate infected, while no significant differences were observed in the other groups.

#### Number of cells

Statistical method 1: Logistic mixed-effects models were applied to test the difference between M1 and M2 in the percentage of cells showing acidification, accounting for group membership.

Time 1

| | Estimate | se | OR | p-value |
|---|---|---|---|---|
| Intercept | -3.7842 | 0.5571 | 0.0227 | <0.0001 |
| groupgeneralist not infected | 1.1209 | 0.5041 | 3.0676 | 0.0262 |
| groupintermediate not infected | 0.5415 | 0.6702 | 1.7185 | 0.4192 |
| groupother not infected | 0.7893 | 0.5811 | 2.2019 | 0.1744 |
| groupspecialist not infected | 1.8373 | 0.4994 | 6.2793 | 0.0002 |
| groupgeneralist infected | 2.6184 | 0.4955 | 13.7142 | <0.0001 |
| groupintermediate infected | 2.1332 | 0.6457 | 8.4415 | 0.001 |
| groupother infected | 1.827 | 0.5671 | 6.2155 | 0.0013 |
| groupspecialist infected | 2.8635 | 0.4992 | 17.5222 | <0.0001 |
| M2 | 0.8475 | 0.3018 | 2.3338 | 0.005 |

A significant difference in the proportion of acidifying cells was observed between M1 and M2 (0.005).

Time 2

| | Estimate | se | OR | p-value |
|---|---|---|---|---|
| Intercept | -3.1748 | 0.4154 | 0.0418 | <0.0001 |
| groupgeneralist not infected | 0.6032 | 0.4181 | 1.8279 | 0.1491 |
| groupintermediate not infected | -0.2386 | 0.6771 | 0.7877 | 0.7245 |
| groupother not infected | 1.0186 | 0.4377 | 2.7693 | 0.0200 |
| groupspecialist not infected | 0.6341 | 0.4030 | 1.8853 | 0.1157 |
| groupgeneralist infected | 1.6096 | 0.3937 | 5.0006 | <0.0001 |
| groupintermediate infected | 1.6095 | 0.5028 | 5.0003 | 0.0014 |
| groupother infected | 1.9249 | 0.4217 | 6.8544 | <0.0001 |
| groupspecialist infected | 1.5695 | 0.3910 | 4.8045 | 0.0001 |
| M2 | 0.0117 | 0.1991 | 1.0118 | 0.9532 |

No significant difference in the proportion of acidifying cells was observed between M1 and M2.

Statistical method 2: Logistic mixed-effects models were applied to test the difference between M1 and M2 within each group.

Time 1

| | Estimate | se | OR | p-value |
|---|---|---|---|---|
| Intercept | -3.3699 | 0.6497 | 0.0344 | <0.0001 |
| groupgeneralist not infected | 0.0635 | 0.7231 | 1.0655 | 0.9301 |
| groupintermediate not infected | -0.5919 | 0.9997 | 0.5533 | 0.5538 |
| groupother not infected | 0.8454 | 0.7901 | 2.3289 | 0.2846 |
| groupspecialist not infected | 1.7948 | 0.6766 | 6.0184 | 0.008 |
| groupgeneralist infected | 1.5876 | 0.695 | 4.892 | 0.0224 |
| groupintermediate infected | 1.1566 | 0.8967 | 3.179 | 0.1971 |
| groupother infected | 1.5115 | 0.7781 | 4.5333 | 0.0521 |
| groupspecialist infected | 2.9336 | 0.677 | 18.7945 | <0.0001 |

| | | | | |
|---|---|---|---|---|
| M2 | 0.1024 | 0.7596 | 1.1079 | 0.8927 |
| groupgeneralist not infected:M2 | 1.9428 | 0.9801 | 6.978 | 0.0475 |
| groupintermediate not infected:M2 | 2.0135 | 1.2882 | 7.4897 | 0.118 |
| groupother not infected:M2 | -0.1649 | 1.0789 | 0.848 | 0.8785 |
| groupspecialist not infected:M2 | 0.002 | 0.9402 | 1.002 | 0.9983 |
| groupgeneralist infected:M2 | 1.9113 | 0.9547 | 6.7622 | 0.0453 |
| groupintermediate infected:M2 | 1.8178 | 1.199 | 6.1583 | 0.1295 |
| groupother infected:M2 | 0.5581 | 1.0481 | 1.7473 | 0.5944 |
| groupspecialist infected:M2 | -0.2412 | 0.9401 | 0.7857 | 0.7975 |

The probability of acidification was significantly different between M1 and M2 in the generalist not Infected and generalist Infected groups, while no significant differences were observed in the other groups.

Time 2

| | Estimate | se | OR | p-value |
|---|---|---|---|---|
| Intercept | -3.1815 | 0.5058 | 0.0415 | <0.0001 |
| groupgeneralist not infected | 0.4609 | 0.5746 | 1.5856 | 0.4224 |
| groupintermediate not infected | 0.3962 | 0.815 | 1.4862 | 0.6269 |
| groupother not infected | 0.2905 | 0.6173 | 1.3371 | 0.6379 |
| groupspecialist not infected | 0.5087 | 0.5395 | 1.6631 | 0.3457 |
| groupgeneralist infected | 2.0166 | 0.5151 | 7.5124 | 0.0001 |
| groupintermediate infected | 2.3323 | 0.6226 | 10.3013 | 0.0002 |
| groupother infected | 1.1574 | 0.559 | 3.1816 | 0.0384 |
| groupspecialist infected | 1.663 | 0.5156 | 5.2751 | 0.0013 |
| M2 | 0.0421 | 0.6478 | 1.043 | 0.9482 |
| groupgeneralist not infected:M2 | 0.2377 | 0.7987 | 1.2683 | 0.766 |
| groupintermediate not infected:M2 | -1.3997 | 1.4084 | 0.2467 | 0.3203 |
| groupother not infected:M2 | 1.3791 | 0.8256 | 3.9714 | 0.0949 |
| groupspecialist not infected:M2 | 0.256 | 0.759 | 1.2917 | 0.7359 |
| groupgeneralist infected:M2 | -0.8395 | 0.7401 | 0.4319 | 0.2566 |
| groupintermediate infected:M2 | -1.7714 | 0.9715 | 0.1701 | 0.0683 |
| groupother infected:M2 | 1.4896 | 0.7768 | 4.4352 | 0.0551 |
| groupspecialist infected:M2 | -0.1638 | 0.7323 | 0.8489 | 0.823 |

No significant difference in the proportion of acidifying cells was observed between M1 and M2.

#### Colocalization index

Statistical method 1: Linear mixed-effects models were applied to test the difference between M1 and M2, accounting for group membership.

Time 1 (Ordered Quantile normalization transformation)

| | Value | Std.Error | p-value |
|---|---|---|---|
| Intercept | 0.1125 | 0.0645 | 0.0815 |
| groupintermediate infected | -0.0107 | 0.0931 | 0.9088 |
| groupother infected | 0.1109 | 0.0842 | 0.1902 |
| groupspecialist infected | -0.1970 | 0.0748 | 0.0094 |
| M2 | -0.1651 | 0.0560 | 0.0037 |

M2 values are significantly different from the M1 values (p=0.0037).

Time 2 (Ordered Quantile normalization transformation)

| | Value | Std.Error | p-value |
|---|---|---|---|
| Intercept | 0.1148 | 0.0748 | 0.1251 |
| groupintermediate infected | -0.2001 | 0.1092 | 0.0692 |
| groupother infected | 0.0296 | 0.0904 | 0.7435 |
| groupspecialist infected | -0.2184 | 0.0811 | 0.0080 |
| M2 | -0.0640 | 0.0600 | 0.2878 |

M2 values are not significantly different from the M1 values (p=0.2878).

Statistical method 2: Linear mixed-effects models were applied to test the difference between M1 and M2 within each group.

Time 1 (Ordered Quantile normalization transformation)

| | Value | Std.Error | p-value |
|---|---|---|---|
| (Intercept) | 0.0680 | 0.0754 | 0.3673 |
| groupintermediate infected | 0.1731 | 0.1295 | 0.1835 |
| groupother infected | 0.1407 | 0.1156 | 0.2257 |
| groupspecialist infected | -0.1541 | 0.1022 | 0.1339 |
| M2 | -0.0779 | 0.0949 | 0.4132 |
| groupintermediate infected:M2 | -0.3662 | 0.1800 | 0.0439 |
| groupother infected:M2 | -0.0589 | 0.1536 | 0.7018 |
| groupspecialist infected:M2 | -0.0848 | 0.1381 | 0.5402 |

M2 values were significantly different from M1 values in group intermediate infected, while no significant differences were observed in the other groups.

Time 2 (Ordered Quantile normalization transformation)

| | Value | Std.Error | p-value |
|---|---|---|---|
| Intercept | 0.2052 | 0.0851 | 0.0160 |
| groupintermediate infected | -0.4874 | 0.1471 | 0.0012 |
| groupother infected | -0.0520 | 0.1184 | 0.6610 |
| groupspecialist infected | -0.3424 | 0.1074 | 0.0018 |
| M2 | -0.2564 | 0.1024 | 0.0135 |
| groupintermediate infected:M2 | 0.5763 | 0.2039 | 0.0055 |
| groupother infected:M2 | 0.1825 | 0.1579 | 0.2500 |
| groupspecialist infected:M2 | 0.2632 | 0.1440 | 0.0699 |

M2 values were significantly different from M1 values in groups generalist infected and intermediate infected, while no significant differences were observed in the other groups.

#### N of mycobacteria per cell (M/C)

Statistical method 1: Linear mixed-effects models were applied to test the difference between M1 and M2, accounting for group membership.

Time 1 (Ordered Quantile normalization transformation)

| | Value | Std.Error | p-value |
|---|---|---|---|
| Intercept | 0.0064 | 0.1018 | 0.9498 |
| groupintermediate infected | -0.1199 | 0.1014 | 0.2393 |
| groupother infected | -0.0270 | 0.0945 | 0.7752 |
| groupspecialist infected | -0.0202 | 0.0830 | 0.8086 |
| M2 | 0.0091 | 0.0589 | 0.8775 |

M2 values are not significantly different from the M1 values (p=0.8775).

Time 2 (Ordered Quantile normalization transformation)

| | Value | Std.Error | p-value |
|---|---|---|---|
| (Intercept) | 0.0738 | 0.1102 | 0.5031 |
| groupintermediate infected | -0.0756 | 0.1033 | 0.4656 |
| groupother infected | 0.2127 | 0.0901 | 0.0197 |
| groupspecialist infected | -0.1456 | 0.0801 | 0.0714 |
| M2 | -0.1149 | 0.0566 | 0.0442 |

M2 values are significantly different from the M1 values (p=0.0442).

Statistical method 2: Linear mixed-effects models were applied to assess the difference between M1 and M2 within each group.

Time 1

| | Value | Std.Error | p-value |
|---|---|---|---|
| Intercept | 0.0192 | 0.1103 | 0.8622 |
| groupintermediate infected | -0.1487 | 0.1420 | 0.2970 |
| groupother infected | -0.0180 | 0.1273 | 0.8877 |
| groupspecialist infected | -0.0536 | 0.1115 | 0.6318 |
| M2 | -0.0159 | 0.1019 | 0.8763 |
| groupintermediate infected:M2 | 0.0575 | 0.1946 | 0.7682 |
| groupother infected:M2 | -0.0176 | 0.1647 | 0.9151 |
| groupspecialist infected:M2 | 0.0666 | 0.1468 | 0.6510 |

M2 values are not significantly different from the M1 values.

Time 2

| | Value | Std.Error | p-value |
|---|---|---|---|
| (Intercept) | 0.0224 | 0.1155 | 0.8462 |
| groupintermediate infected | 0.0998 | 0.1380 | 0.4709 |
| groupother infected | 0.3612 | 0.1164 | 0.0024 |
| groupspecialist infected | -0.1468 | 0.1041 | 0.1607 |
| M2 | -0.0088 | 0.0958 | 0.9267 |
| groupintermediate infected:M2 | -0.3575 | 0.1920 | 0.0648 |
| groupother infected:M2 | -0.2963 | 0.1515 | 0.0527 |
| groupspecialist infected:M2 | -0.0044 | 0.1360 | 0.9743 |

M2 values are not significantly different from the M1 values.
