## Supplementary 4 for "*Mycobacterium tuberculosis* diversity and macrophage heterogeneity dictate phagosomal acidification"

***AUTOPHAGY***

***MATERIALS & METHODS***

**Live imaging microscopy**

Induction of autophagosomes, live imaging: M1 and M2 macrophages were stained with Hoechst 33342 for nuclei (Ex/Em 450-461 nm), and with CYTO-ID® Green (Ex/Em 499-548 nm) from the CYTO-ID® autophagy detection kit 2.0 (Enzo Biochem, Farmingdale, NYS, USA) according to manufacturer's instructions for monitoring induction of autophagosomes. The dye exhibits bright fluorescence upon incorporation into pre-autophagosomes, autophagosomes, and auto-phagolysosomes, whereas it shows limited staining of lysosomes [1]. Imaging experiments were performed at 24h p.i. As control of autophagy induction, macrophages were treated with 500nM of rapamycin and 50µM of chloroquine overnight. For each strain, 8 images (2 positions for each well out of 4 wells per strain) were acquired as *z*-stacks (1.48 µm x 4 stacks) in at least two independent experiments.

**Images and data analysis**

Induction of autophagosomes: Single-cells were manually segmented. Autophagosomes and mycobacteria were selected by thresholding the 488 nm and the 561 nm channel, respectively. CYTO-ID signal was acquired as mean signal intensity in cells ROI. mCherry mean signal across the cells ROI has been used as a proxy for mycobacterial burden. The autophagic pathway was analyzed by considering both the number and the size of autophagosomes [2]. The number of autophagosomes per cell was calculated using MatLab.

***RESULTS***

**Autophagic flux**

We evaluated the autophagic flux in THP-1 M1 and M2 macrophages infected with H37Rv, H37Ra, L6, and L2 at 24 h p.i. by the use of the CYTO-ID kit (Figure S1). A total of 1960 single cells were considered for the analysis (on average, for each strain tested, M1: 68±27 not infected cells + 67±14 infected cells; M2: 64±22 not infected cells + 63±9 infected cells).

| *24 h p.i.* | | | | | | | |
|---|---|---|---|---|---|---|---|
| | | **NOT infected** | | **Infected** | | ***TOT*** | |
| | | **N of cells** | **Replicates** | **N of cells** | **Replicates** | **Not inf** | **Inf** |
| **M1** | **CTRL** | 121 | 5 | *na* | *na* | ***609*** | ***403*** |
| | **RapaChlo** | 95 | 5 | *na* | *na* | | |
| | **DMSO** | 67 | 5 | *na* | *na* | | |
| | **H37Rv** | 48 | 3 | 80 | 3 | | |
| | **H37Ra** | 49 | 2 | 51 | 2 | | |
| | **Beijing A** | 77 | 3 | 84 | 3 | | |
| | **Beijing B** | 37 | 2 | 57 | 2 | | |
| | **Afri A** | 65 | 3 | 76 | 3 | | |
| | **Afri B** | 50 | 2 | 55 | 2 | | |
| **M2** | **CTRL** | 108 | 5 | *na* | *na* | ***572*** | ***376*** |
| | **RapaChlo** | 87 | 5 | *na* | *na* | | |
| | **DMSO** | 66 | 5 | *na* | *na* | | |
| | **H37Rv** | 64 | 3 | 70 | 3 | | |
| | **H37Ra** | 46 | 2 | 55 | 2 | | |
| | **Beijing A** | 40 | 3 | 74 | 3 | | |
| | **Beijing B** | 45 | 2 | 59 | 2 | | |
| | **Afri A** | 64 | 3 | 66 | 3 | | |
| | **Afri B** | 52 | 2 | 52 | 2 | | |
| | ***TOT*** | ***1181*** | | ***779*** | | | |
| | | **1960** | | | | | |

| | |
|---|---|
| *Acquisition:* | obj. **63x, oil NA 1.40**<br>**z-stacks: 1.48 µm x 4 stacks**<br>Res. **16-bit**, Format: **1024x1024**<br>Laser 405 (for Hoechst) = **40%** gain: 935; offset: 0%<br>Laser 488 (for CYTO-ID) = 2**0% (Argon power: 30%); gain: 780**, offset: 0%<br>Laser 561 (for mCherry) = 1**8%; gain: 905**, offset 0%<br>well/sample: 2+2<br>acquistion postions/well: 2 |
| *Analysis:* | 24 h p.i.; **N of autophagosomes, size of autophagosomes** |
| **Hoechst (Ex/Em 350/461)** | |
| **CYTO-ID (Ex/Em 499/548)** | **threshold (Default, Red; no flags on Dark background and Stack histogram) = 160**<br>**size = 5-infitity pixel units, circularity = 0-1 (show: nothing; Flags on Add to Manager, Display results)** |
| **mCherry (Ex/Em 587-610)** | |

The size and the number of autophagosomes were considered, and strains were grouped as previously reported in Table 1.

### Autophagy

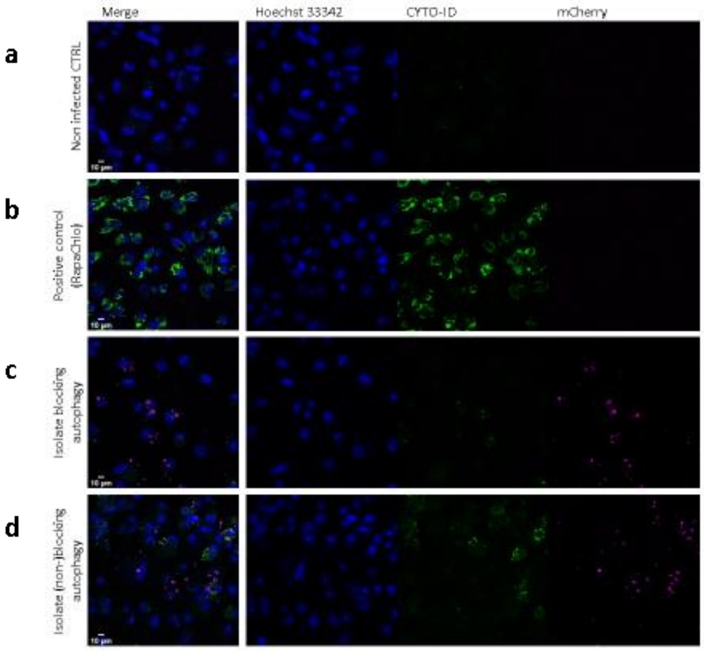


**Fig. S3.1**: THP-1 cells non-infected (a); treated with autophagy inducer (b); infected with autophagy-blocking bacteria (c); infected with autophagy non-blocking bacteria (d). The fluorescence images were acquired after staining with CYTO-ID (Green) and Hoechst (Blue). Bacteria are reported in red. The reported images are illustrative of the pipeline adopted

In M1 macrophages, only the group "other" showed some degree of autophagy; indeed, whereas also generalist and specialist groups show an increase in the number of autophagosomes (Figure S3.2, panel a), their size does not show a statistically increase (Figure S3.2, panel b), thus suggesting poor maturation and progression in the autophagic flux. In M2 macrophages, despite a certain increase in the number of autophagosomes (Figure S3.2, panel c), their size does not show a statistically significant increase (Figure S3.2, panel d), thus suggesting poor maturation and progression in the autophagic flux. None of the group is inducing autophagy. No correlation between number of mycobacteria internalized per cell (M/C) and autophagy was found.

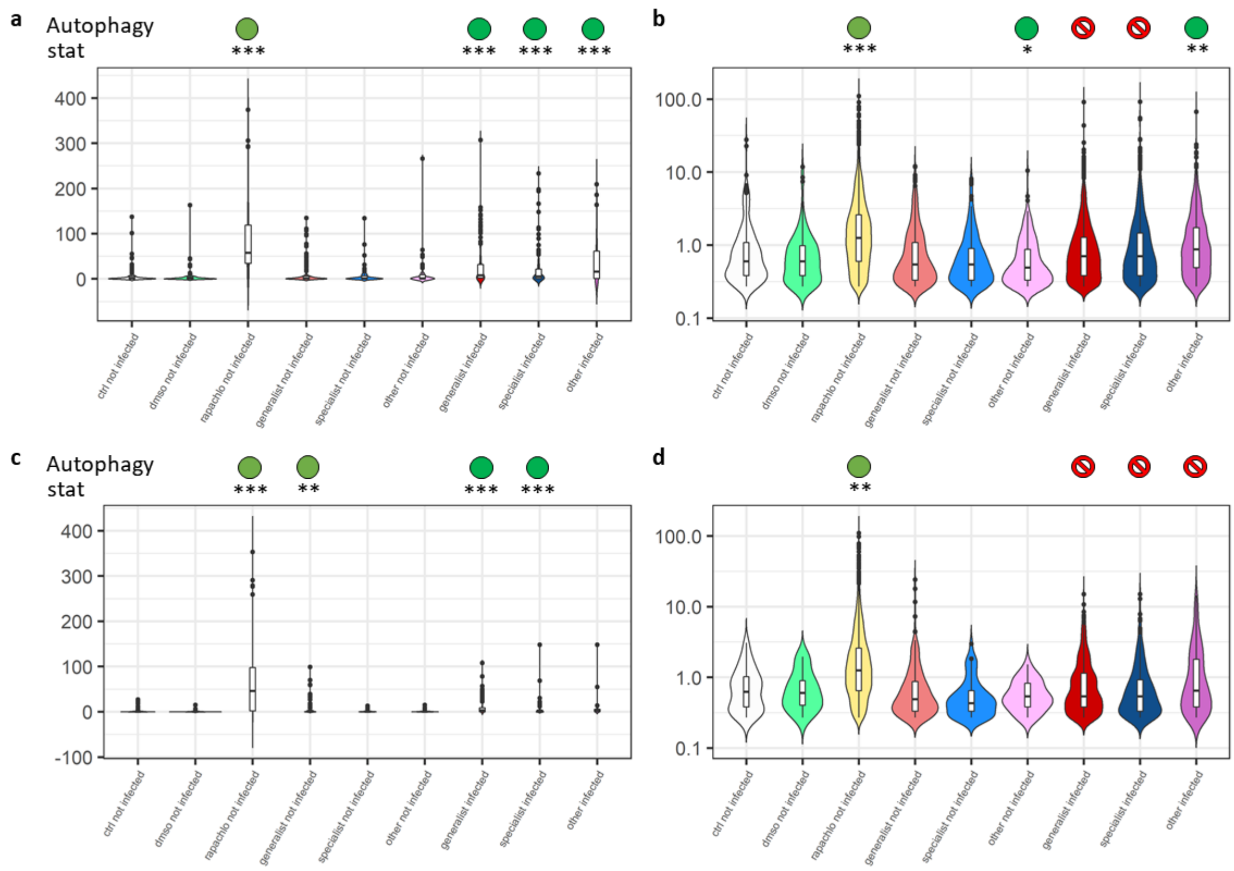


**Figure S3.2**: Autophagic flux at 24h p.i. Autophagosomes number in M1 (a and b panels, respectively) and M2 (c and d panels, respectively) macrophages. Stat = statistical analysis compared to non-infected controls. In the “autophagy” line a visual interpretation is reported as follow: green circle = autophagic flux observed (statistically significant difference compared to non-infected controls); red prohibition sign = autophagic flux not observed (blocked) (no statistically significant difference compared to non-infected controls).

These findings indicate that none of the isolates trigger proper autophagy at the time points considered, irrespectively of the macrophage sub-type. To be noted that in both cases the number of autophagosomes increased, thus suggesting that the pathway was somehow activated but blocked in its full maturation.

*M. tuberculosis* may evade macrophage clearance due to autophagy [3]. Our data did not show relevant differences among less virulent, ancient, specialist, and more virulent, modern, generalist strains as either the categories can block the formation of autophagosomes. Previous findings showed similar results, despite highlighting a difference in the mechanism used by different genotypes to hinder autophagy [3, 4].

### REFERENCES


1. Oeste CL, Seco E, Patton WF, Boya P, Pérez-Sala D. Interactions between autophagic and endo-lysosomal markers in endothelial cells. Histochem Cell Biol **2013**; 139:659-70.

2. Klionsky DJ, Abdelmohsen K, Abe A, et al. Guidelines for the use and interpretation of assays for monitoring autophagy (3rd edition). Autophagy **2016**; 12:1-222.

3. Haque MF, Boonhok R, Prammananan T, et al. Resistance to cellular autophagy by Mycobacterium tuberculosis Beijing strains. Innate Immun **2015**; 21:746-58.

4. Romagnoli A, Petruccioli E, Palucci I, et al. Clinical isolates of the modern Mycobacterium tuberculosis lineage 4 evade host defense in human macrophages through eluding IL-1β-induced autophagy. Cell Death Dis **2018**; 9:624.
