## Supplementary 5 for "*Mycobacterium tuberculosis* diversity and macrophage heterogeneity dictate phagosomal acidification"

***APOPTOSIS AND NECROSIS***

***MATERIALS & METHODS***

**Live imaging microscopy**

Induction of apoptosis and necrosis, live imaging: M1 and M2 macrophages were stained with CytoCalcein Violet for cytosol (Ex/Em 405-450 nm, a marker for living cells cytoplasm), with Apopxin Green (Ex/Em 490-525 nm, apoptotic marker), and 7-AAD (Ex/Em 546/657, necrotic marker) according to the Apoptosis/Necrosis assay kit (Abcam, Cambridge, UK) for monitoring apoptosis/necrosis at 24h p.i. Apopxin targets the hallmark of apoptosis-exposed phosphatidylserine. Loss of plasma membrane integrity represents a straightforward approach to demonstrate late-stage apoptosis and necrosis, and it is detected by the ability of the membrane-impermeable 7-AAD to label the nucleus. As control of apoptosis, macrophages were treated with 5µg/mL cyclohexamide and 2ng/mL TNFα overnight. For each strain, 4 images (2 positions for each well out of 2 wells per strain) were acquired as *z*-stacks (1.48 µm x 4 stacks) in at least two independent experiments.

A 488 nm laser was used for CYTO-ID and Apopxin, a 561 nm laser was used for mCherry and 7-AAD.

**Images and data analysis**

Induction of apoptosis and necrosis: Single cells were manually segmented. CytoCalcein Violet, and Apopxin Green were acquired as mean signal intensity in cells ROI. Signal from 7-AAD in the nuclei was never observed at the timepoints considered, thus analysis on the necrosis was not performed. Apoptotic cells have been identified using FlowClust (ver. 3.9), a Bioconductor R package implementing a robust model-based clustering approach based on multivariate t mixture models with the Box-Cox transformation for gating cell populations (rule of identifying outliers: 83% quantile) [1]. High mycobacterial burden was considered when mCherry signal was found to be equal or above the central value of the distribution of all mCherry signals in infected cells. Signals below this value were defined as low mycobacterial burden.

### *RESULTS*

#### Apoptosis and necrosis

We evaluated the induction of apoptosis and necrosis THP-1 M1 and M2 macrophages infected with H37Rv, H37Ra, L6, and L2 at 24h p.i. (Figure S4.1). A total of 2470 single cells were considered for the analysis (on average, for each strain tested, M1: 84±34 not infected cells + 75±13 infected cells; M2: 87±44 not infected cells + 79±15 infected cells).

| *24 h p.i.* | | | | | | | |
|---|---|---|---|---|---|---|---|
| | | **NOT infected** | | **Infected** | | ***TOT*** | |
| | | **N of cells** | **Replicates** | **N of cells** | **Replicates** | **Not inf** | **Inf** |
| | **CTRL** | 138 | 2 | *na* | *na* | | |
| | **H37Rv** | 88 | 2 | 70 | | | |
| | **H37Ra** | 94 | 2 | 82 | | | |
| **M1** | **Beijing A** | 95 | 2 | 94 | | ***689*** | ***451*** |
| | **Beijing B** | 69 | 2 | 69 | | | |
| | **Afri A** | 117 | 2 | 80 | | | |
| | **Afri B** | 88 | 2 | 56 | | | |
| | **CTRL** | 164 | 2 | *na* | *na* | | |
| | **H37Rv** | 86 | 2 | 82 | 2 | | |
| | **H37Ra** | 107 | 2 | 81 | 2 | | |
| **M2** | **Beijing A** | 128 | 2 | 76 | 2 | ***725*** | ***475*** |
| | **Beijing B** | 60 | 2 | 103 | 2 | | |
| | **Afri A** | 85 | 2 | 76 | 2 | | |
| | **Afri B** | 95 | 2 | 57 | 2 | | |
| | ***TOT*** | ***1414*** | | ***926*** | | | |
| | | | | **2340** | | | |

| | |
|---|---|
| *Acquisition:* | obj. **63x, oil NA 1.40**<br>**z-stacks: 1.48 µm x 4 stacks**<br>Res. **16-bit**, Format: **1024x1024**<br>Laser 405 (for CytoCalcein Violet) = **6% gain: 625**; offset: 0%; **pinhole: 95.60**<br>Laser 488 (for Apopxin) = **10% (Argon power: 20%); gain: 670;** offset: 0%; **pinhole: 330**<br>Laser 561 (for mCherry and 7-AAD) = **8%; gain: 700;** offset 0%; **pinhole: 95.60**<br>well/sample: 2<br>acquistion postions/well: 2 |
| *Analysis:* | 24 h p.i.; **percentages of apoptotic/necrotic/live cells** |
| **Hoechst (Ex/Em 405/450)** | |
| **Apopxin (Ex/Em 490/525)** | |
| **7-AAD (Ex/Em 546/657)** | |
| **mCherry (Ex/Em 587-610)** | |

Automated gating of apopxin-positive (apoptosis) and cytocalcein-positive (live) cells using flowClust [1] (Rule for identifying outliers: 83% quantile; Number of outliers: 232 (9.39%)).

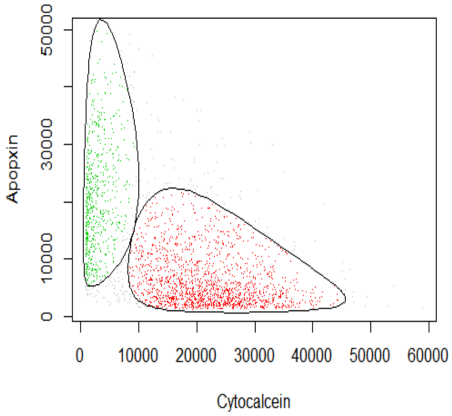


Despite the marker for necrosis 7-AAD had a spectrum overlapping to one of mCherry mycobacteria (7-AAD Ex/Em 546/657; mCherry Ex/Em 587-610 nm) and both have been imaged using the 561 laser, the morphology of the bacteria and necrotic cells was completely different and easily distinguishable (Figure S4.1). However, in our testing conditions (24h p.i.) necrosis was never observed, thus it was not further considered.

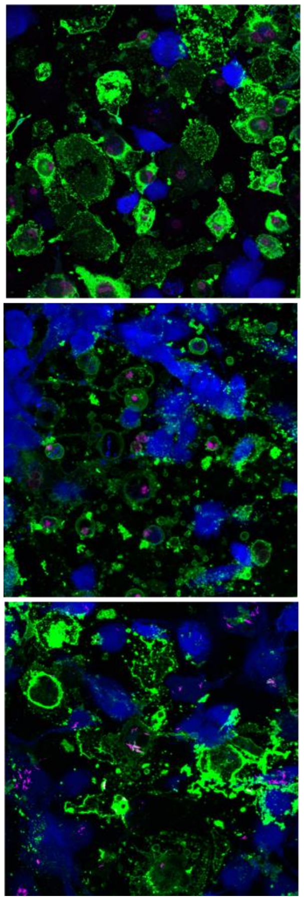

Induced necrosis:
Starvation (uninfected THP-1 derived macropahges)

7-AAD

Induced necrosis:
Cyclohexamide + TNFα (uninfected THP-1 derived macropahges)

7-AAD

Infected THP-1 derived macrophages

mCherry (mycobacteria)

**Figure S4.1**: Morphology of the bacteria and necrotic cells was completely different and easily distinguishable.

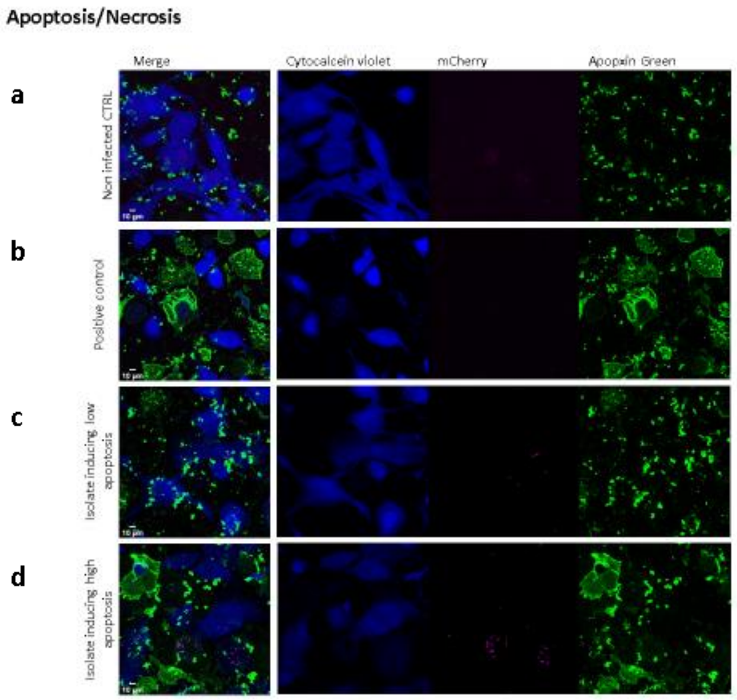


**Figure S4.2**: THP-1 cells non-infected (a); treated with apoptosis inducer (b); infected with apoptosis-blocking bacteria (c); infected with apoptosis non-blocking bacteria (d). The fluorescence images were acquired after staining with CytoCalcein Violet (Blue) and Apopxin Green (Green). Bacteria are reported in red. The reported images are illustrative of the pipeline adopted

In M1 macrophages, bystander non-infected cells of generalist and other strains showed increased apoptosis (40%) (Figure S4.3, panel a). Despite not statistically significant, also macrophages infected with generalist strains showed a similar increase in the percentage of apoptosis. Stratification of the data by mycobacterial burden showed higher apoptotic levels in bystander non-infected cells or in cells infected with lower mycobacterial load (Figure S4.3, panel b). In M2 macrophages, only bystander non-infected cells of other strains showed a slight, statistically not significant increase in apoptosis (30%) (Figure S4.3, panels c and d).

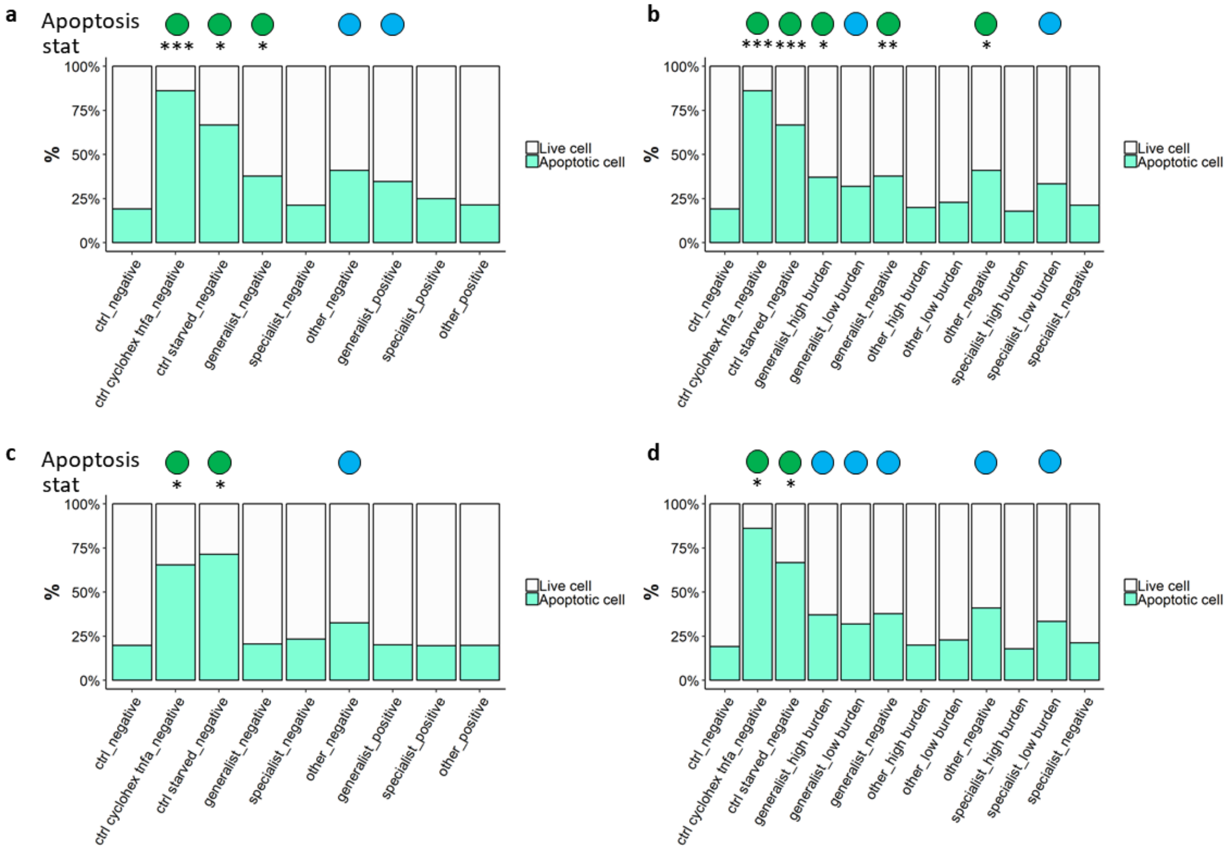


**Figure S4.3:** Apoptosis at 24h p.i. Percentage of live/apoptotic M1 (panel a) and M2 (panel c) macrophages. Panels b (M1) and d (M2) report percentages of live/apoptotic cells stratified by mycobacterial burden. Stat = statistical analysis compared to non-infected controls. In the “apoptosis” line a visual interpretation is reported as follow: green circle = apoptosis observed (statistically significant difference compared to non-infected controls); blue circle = acidification trend observed but not statistically significant.

These findings indicate that MTB infection with isolates other than specialists can trigger apoptosis mainly in M1 macrophages and related bystander non-infected cells.

The role of apoptosis on the MTB pathogenesis is debatable with reports supporting a protective effect for the host contributing to infection control, while others suggesting a detrimental effect increasing mycobacterial dissemination [2-5]. In the present study, apoptosis was observed only in bystander non-infected cells of more virulent, modern and generalist isolates. Induction of apoptosis by virulent strains seems contradictory to its protective role against MTB. However, previous findings highlighted that apoptosis signals at day 1 p.i. progressed to necrosis at later timepoints; furthermore, exposure of phosphatidylserine is observed in different forms of programmed cell death (namely apoptosis, necroptosis and ferroptosis) [6]. Apoptosis data are in line with what we observed with the cell survival assay, where in M1 macrophages, more virulent strains induce a major cells death rate compared to less virulent strains, while in M2 cells we found lower induction of apoptosis, with no statistical differences among MTB categories. Furthermore, the correlation between apoptosis and lower bacterial burden observed further support the protective role of apoptosis.

**REFERENCES**

1. Lo K, Hahne F, Brinkman RR, Gottardo R. flowClust: a Bioconductor package for automated gating of flow cytometry data. BMC Bioinformatics **2009**; 10:145.

2. Porcelli SA, Jacobs WR. Tuberculosis: unsealing the apoptotic envelope. Nat Immunol **2008**; 9:1101-2.

3. Davis JM, Ramakrishnan L. The role of the granuloma in expansion and dissemination of early tuberculous infection. Cell **2009**; 136:37-49.

4. Aguiló N, Marinova D, Martín C, Pardo J. ESX-1-induced apoptosis during mycobacterial infection: to be or not to be, that is the question. Front Cell Infect Microbiol **2013**; 3:88.

5. Behar SM, Briken V. Apoptosis inhibition by intracellular bacteria and its consequence on host immunity. Curr Opin Immunol **2019**; 60:103-10.

6. Amaral EP, Costa DL, Namasivayam S, et al. A major role for ferroptosis in. J Exp Med **2019**; 216:556-70.
