## Supplementary 6 for "*Mycobacterium tuberculosis* diversity and macrophage heterogeneity dictate phagosomal acidification"

***MTB INTRAMACROPHAGE SURVIVAL***

**Intramacrophage MTB survival**

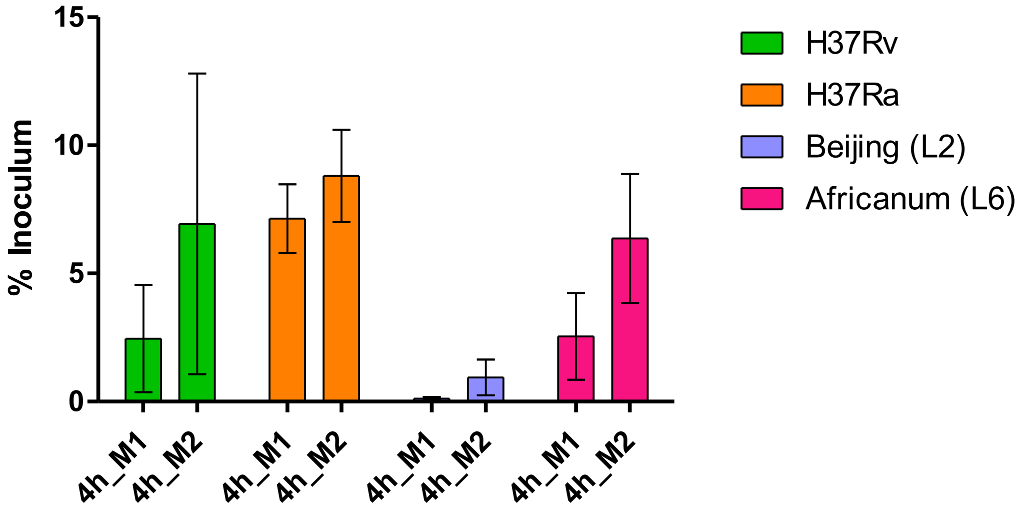


**Figure S5.1**: CFU results at 4h p.i. (% of the inoculum). Beijing: Generalist/L2; Africanum: Specialist/L6

Differences are present considering M1/M2 cell phenotype: entry of bacteria is increased in M2 cells in all MTB strains selected. Higher CFU levels are found in H37Ra infected cells, while H37Rv and L6 show a similar CFU level. L2 uptake resulted to be very low compared to all other strains, with CFU values close to 1% of the inoculum in M1 cells and to 3% in M2 cells.
